# Lesion environments direct transplanted neural progenitors towards a wound repair astroglial phenotype

**DOI:** 10.1101/2022.01.24.477530

**Authors:** T.M. O’Shea, Y. Ao, S. Wang, A.L. Wollenberg, J.H. Kim, R.A. Ramos Espinoza, A. Czechanski, L.G Reinholdt, T.J. Deming, M.V. Sofroniew

## Abstract

Neural progenitor cells (NPC) represent potential cell transplantation therapies for CNS injuries. To understand how lesion environments influence transplanted NPC fate *in vivo*, we derived NPC expressing a ribosomal protein-hemagglutinin tag (RiboTag) for transcriptional profiling of transplanted NPC. Here, we show that NPC grafted into uninjured CNS generate cells that are transcriptionally similar to healthy astrocytes and oligodendrocyte lineages. In striking contrast, NPC transplanted into serum-exposed CNS lesions after stroke or spinal cord injury generate cells that share transcriptional, morphological and functional features with newly proliferated host astroglia that restrict inflammation and fibrosis and thereby protect adjacent neural tissue. Our findings reveal overlapping differentiation potentials of grafted NPC and proliferating host astrocytes; and show that in the absence of other interventions, non-cell autonomous cues in CNS lesions direct the differentiation of grafted NPC predominantly towards a naturally occurring neuroprotective wound repair astroglial phenotype.

## Introduction

Neural tissue that is lost to injury or disease in the mature mammalian central nervous system (CNS) is not spontaneously replaced. Instead, naturally occurring and conserved CNS wound repair generates lesions in which non-neural lesion cores of fibrotic and inflammatory cells are partitioned from adjacent preserved neural tissue by newly formed astroglial borders^1-10^. Although this wound repair response is effective in clearing debris, limiting infection and protecting nearby viable neural tissue, the resulting lesions often contain large volumes of non-neural fibrotic scar tissue that lacks the specialized neural cells necessary to support axon regeneration or the remodeling of neural circuits^10-14^.

Neural cell transplantation represents one potential therapeutic strategy for replacing lost neural tissue and improving outcome after CNS insults^15-18^. Different types of cell transplantation are being explored for this purpose including fetal cell grafts^19-24^, adult neural progenitor cells (NPC)^25-27^, and NPC derived from lines of embryonic stem cells (ESC) or induced pluripotent stem cells (iPSC)^28-32^. Despite considerable progress in the derivation, production and transplantation of NPC into CNS injuries, many questions remain about the roles of cell autonomous versus non-cell autonomous factors in determining NPC differentiation, and their neural repair support functions, after grafting *in vivo*^33^.

Here, we examined how transplantation of NPC into different CNS environments altered their gene expression and differentiation fate *in vivo*. To do so we derived NPC from mouse embryonic stem cells (ESC) that constitutively express the ribosomal protein Rpl22 with a hemagglutinin (HA) tag (Rpl22-HA), also known as RiboTag, which permits cell-specific transcriptional profiling by RNA sequencing (RNAseq) and immunohistochemical characterization of HA-positive cells ^34,35^. Using Rpl22-HA-expressing NPC (RiboTag-NPC), we selectively evaluated NPC transcriptional profiles and differentiation fates following transplantation into uninjured CNS or into CNS lesions after forebrain stroke or spinal cord injury. We compared NPC transcriptional responses *in vivo* with NPC transcriptional responses *in vitro* to specific non-cell autonomous molecular cues that modified their differentiation; and we compared NPC differentiation phenotypes *in vivo* with the profiles of newly proliferated host astroglia that naturally adopt wound repair functions. We found that non-cell autonomous cues powerfully modify NPC transcription and instruct different differentiation fates both *in vitro* and *in vivo*, and that NPC are directed towards different cell fates by non-cell autonomous cues in uninjured or lesioned CNS tissue. Our findings reveal striking similarities between the transcriptional profiles and cellular morphologies of cells derived from NPC transplanted into CNS lesions and proliferating host astroglia in CNS injuries that are both stimulated into adopting naturally occurring wound repair astroglial phenotypes.

## Results

### Neural induction and expansion of RiboTag mESC derives reproducible and stable NPC lines

Mouse ESC expressing RiboTag through Cre-Lox recombination (Fig. 1a) were used to generate NPC by neural induction and expansion^35,36^ (Fig. 1b). Discrete multicellular ESC colonies (Fig. 1c) were transitioned into spindle-shaped NPC and expanded as adherent monolayer cultures rather than as floating neurospheres (Fig.1d). Transcriptome profiling by bulk RiboTag RNA Sequencing (RNA-Seq) showed gain of NPC phenotype and loss of ESC characteristics assessed using defined panels of canonical genes for each cell type^37^. NPC generation markedly reduced ESC gene expression with median log2Fold Change (FC) of approximately -10 (Fig. 1e,f,h). Concurrently, NPC derivation induced increased expression of canonical neural stem cell genes with a median log2FC of approximately +5 (Fig. 1h). Loss of protein expression of ES markers Dppa4, Oct4, Nanog as well as emergence of NPC markers Nestin, Sox9 and Fabp7 by immunocytochemistry (ICC) and quantitative western blotting (WB) further supported successful NPC generation (Fig.1g, Supplementary Fig. 1h,i). HA-positive ribosome expression was consistently and robustly detected in all cells across all *in vitro* evaluations including in mESC colonies, NPC derivations and differentiated NPC populations (Fig. 1g).

**Fig. 1:**
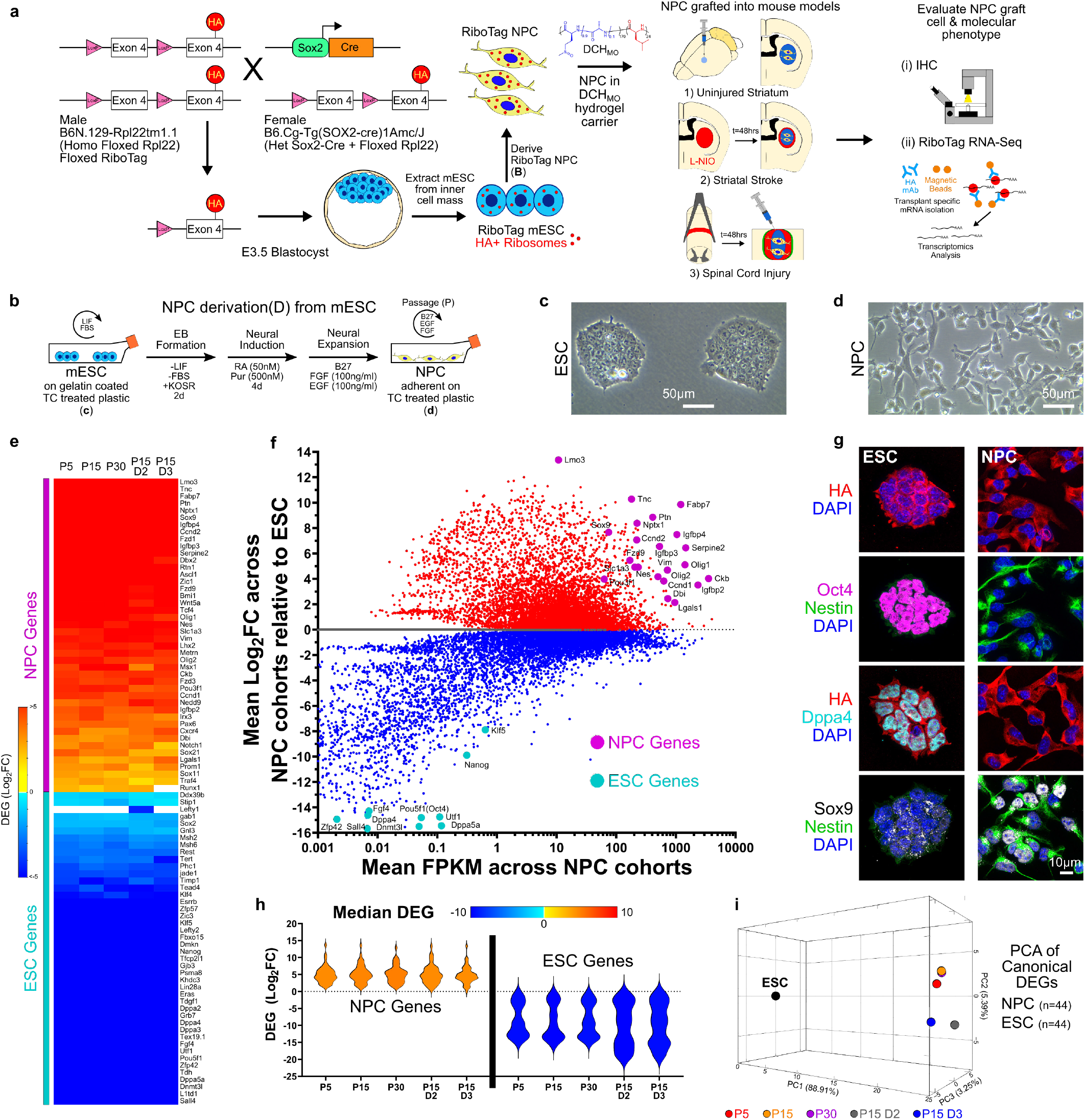
Neural induction and expansion of RiboTag mESC derives reproducible and stable NPC lines. **a** Schematic of RiboTag ESC derivation by Cre-Lox recombination and experimental workflow. **b** Schematic of NPC derivation by sequential neural induction and expansion. **c** Phase contrast image of ESC colonies. **d** Phase contrast image of adherent NPC. **e** Heatmap of DEGs for ESC and NPC gene panels for NPC at different Passages (P) and derivation (D). **f** M-A Plot of mean transcriptome differences averaged across NPC derivations (5 groups, N=4 samples per group) referenced to ESC (N=4 samples) showing increased expression of NPC genes and decreased expression of ESC genes. **g** Immunocytochemistry of ESC and NPC samples showing robust expression of HA-positive ribosomes across both cell types as well as downregulation of ESC markers (Oct4, Dppa4) and upregulation of neural stem markers (Nestin, Sox9) following application of neural induction and expansion procedures. **h** Violin plots of DEGs for the different NPC derivations (D) and passages (P) for the canonical NPC gene and ESC gene panels with color scale based on median DEG for each derivation. **i** 3D PCA of NPC samples from different derivations (D) and passages (P) using canonical NPC gene and ESC gene panels (44 genes for each panel).

To evaluate reproducibility and variability, three unique NPC derivations (D) were generated from different ESC cultures. To assess NPC stability and genetic drift, multiple passages (P) of a single NPC derivation up to passage 30 were generated. Samples from 5 unique NPC groups with different D and P combinations were profiled by bulk RNA-Seq (Fig. 1e,f,h, Supplementary Fig. 1a-g). Principal Component analysis (PCA) of transcriptome differences showed minimal variation amongst NPC groups and comparably large differences between all NPC groups and the ESC state (Fig. 1i, Supplementary Fig. 1a,b). Two independent and complementary metrics, PCA Euclidean distance and Cosine Similarity (CS), showed high correlation amongst the different NPC groups but large and equivalent differences relative to ESC (Supplementary Fig 1c, e-g). Limiting comparisons to 88 DEGs comprising the canonical ESC and NPC gene panels strengthened the similarity amongst NPC groups compared with the full complement of 14,532 DEGs (Supplementary Fig 1e).

These data show that RiboTag-NPC derived from a common ESC stock by standard neural induction and expansion protocols are genetically stable and maintain unaltered RiboTag expression across serial passages and up to 4 months of constant culture with minimal differences between derivation or passage. RiboTag-NPC dominantly express well established canonical neural stem cell genes and significantly downregulate all canonical embryonic stem cell genes making them suitable candidates for further investigation.

### NPC spontaneously differentiate into astrocyte and oligodendroglia lineage cells *in vitro* with phenotype modulated by addition of specific molecules

NPC maintained in serum free medium with mitogens EGF and FGF retain multipotency over serial passages but as expected^38^ undergo spontaneous cell autonomous regulated differentiation (SPONT) over 4 days under reduced EGF/FGF conditions (Fig. 2a). SPONT induced upregulation of astroglial and oligodendroglia lineage genes and downregulation of neural stem and proliferation genes (Fig. 2b). We added CNTF or 1% FBS to promote astrocyte differentiation^36,39,40^; and IGF-1 to promote oligodendroglia differentiation^41^. We compared variations in transcription profiles using reference gene panels for Astrocyte lineage, Oligodendrocyte lineage and Neuronal lineage cells derived from meta-analysis of archival datasets (Supplementary Fig. 2,3 and Source Data). PCA and CS analysis of all 14,017 DEGs (Fig. 2c) or neural cell type specific DEGs (Supplementary Fig. 4a,b) revealed large differences for all differentiated cells compared to the NPC state (defined by PC1) and smaller but measurable variation among the four differentiation conditions (defined by PC2). Astroglia and oligodendroglia lineage phenotypes emerged as the dominant transcriptomic signatures amongst all differentiation conditions, with little overall change in neuronal genes but comprehensive downregulation of NPC and proliferation genes (Fig. 2d). For many astrocyte genes (including *Gfap, Apoe, Pla2g7, Slc1a3, Sparc, Ndrg2, Id3*), the expression levels (by FPKM) after *in vitro* differentiation were comparable to the levels in healthy mouse spinal cord astrocytes *in vivo* (Supplementary Fig. 5 and Source Data) *whereas* other astrocyte genes (*Slc7a10, Myoc, Fam107a, Gjb6, Slc1a2, Aqp4, Aldh1a1, Slc2a4, S100b, Atp1a2*), were not expressed at *in vivo* astrocyte levels but were generally increased compared to NPC. IGF-1 evoked the highest increase in oligodendrocyte gene expression (Fig. 2d, Supplementary Fig. 4a).

**Fig. 2:**
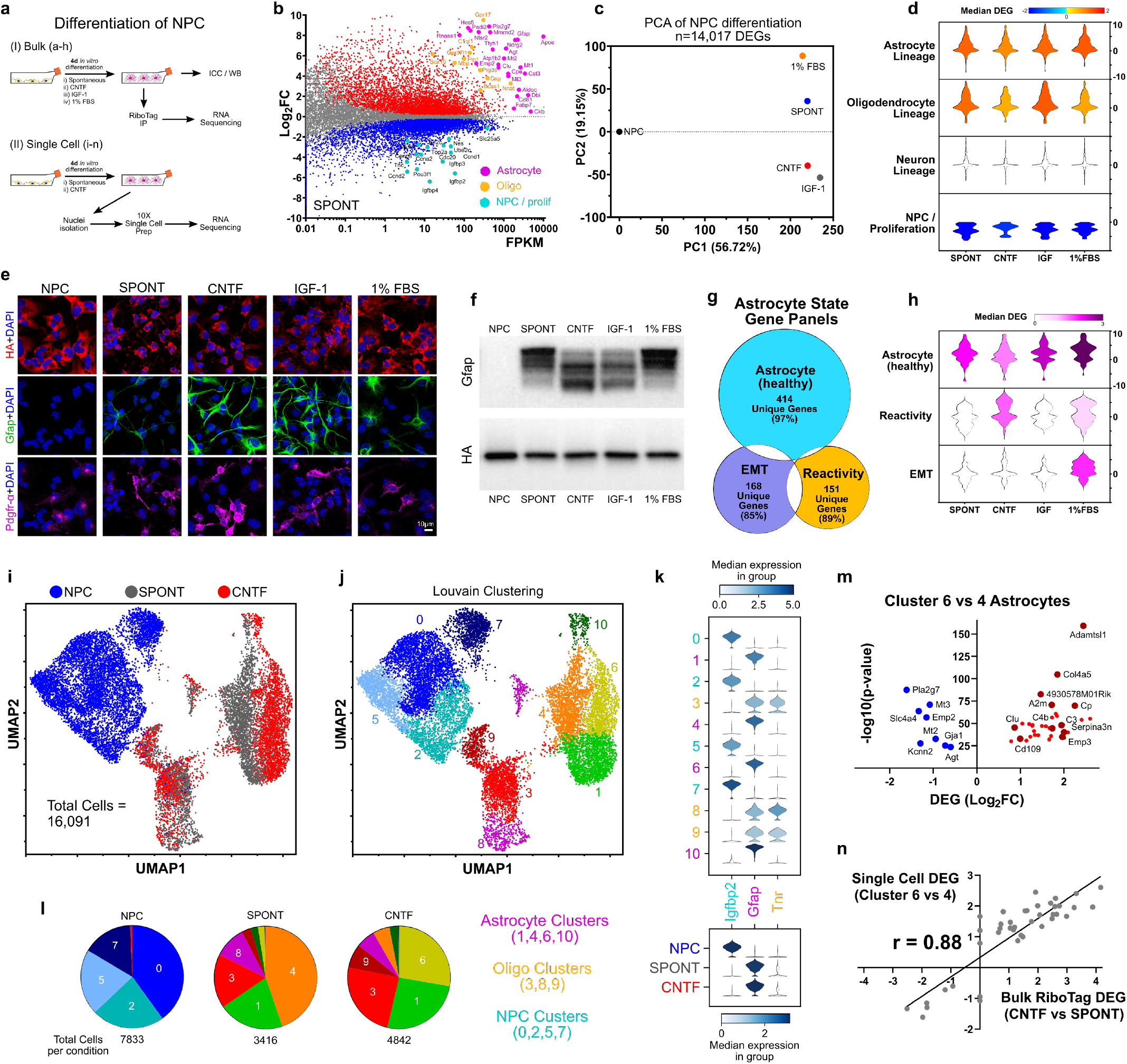
NPC cell autonomously differentiate into astrocyte and oligodendroglia lineage cells *in vitro* with phenotype modulated by specific molecules. **a** Schematic of experimental approach for evaluating *in vitro* NPC differentiation. **b** M-A Plot of Spontaneous (SPONT) differentiation referenced to NPC (N=4 samples) showing increased expression of astrocyte and oligodendroglia lineage genes and decreased expression of NPC/proliferation genes. **c** PCA of all DEGs associated with NPC differentiation across the conditions referenced to NPC showing large differences for all differentiation conditions compared to the NPC state and variation amongst the unique differentiation conditions. **d** Violin plots of DEGs for neural cell specific gene panels. Color of violin plot reflects the value of the median DEG for specified gene panel. **e** ICC staining of the differentiation conditions for astrocyte (Gfap) and oligodendroglia (Pdgfr-α) markers. All DAPI-positive cells across differentiation conditions are RiboTag (HA) positive. **f** Western blot for Gfap and HA showing variation in Gfap isoform bands amongst the various differentiation conditions. HA expression is not altered by differentiation condition. **g** Venn diagram of genes for the different astrocyte states derived through meta-analysis of published data sets showing minimal overlap of healthy astrocyte, reactivity and EMT genes. **h** Violin plots of DEGs from the healthy astrocyte, reactivity and EMT gene panels that are amongst the top 2000 most differentially expressed genes for the differentiation conditions. Color of violin plot reflects the Median DEG for specified gene panel. **i** UMAP of 16,091 single nuclei samples designated by NPC, SPONT and CNTF condition and derived from the top 5000 uniquely expressed genes. **j** UMAP of 16,091 single nuclei samples as in i) segregated by Louvain clustering algorithms into 11 unique clusters. **k** Violin plots of expression values for Igfbp2, Gfap and Tnr for the 11 unique clusters. Igfbp2 is highly expressed in NPC dominant clusters. Gfap is highly expressed in astrocyte clusters. Tnr is highly expressed in oligodendroglia lineage cell clusters. SPONT and CNTF conditions result in predominant astrocyte differentiation. **l** Pie charts for NPC, SPONT and CNTF conditions indicating the predominate cell populations for each condition. Cluster 0 (NPC) predominates in NPC sample. Astrocyte Cluster 4 predominates in SPONT condition whereas Cluster 6 Astrocytes are the dominant cell type in CNTF condition. **m** Volcano plot of DEGs identified by comparing Cluster 6 vs Cluster 4 astrocytes. Cluster 6 astrocytes upregulate numerous genes on the reactivity gene panel and downregulate several canonical healthy astrocyte genes relative to Cluster 4 cells. **n** Correlation analysis of Single nuclei RNA-Seq DEGs for Cluster 6 vs 4 astrocytes compared with bulk RiboTag DEGs for CNTF compared with SPONT. The DEGs identified by both methods show strong correlation (r=0.88).

Gfap-positive astrocytes and Pdgfr-α positive oligodendroglia were detected across all differentiation conditions by ICC, but expression levels and morphology patterns varied markedly (Fig. 2e). Variation in Gfap-positive cell morphology amongst differentiation conditions was associated with differences in Gfap isoform usage ^42^ demonstrated by WB staining bands to mouse monoclonal and rabbit polyclonal Gfap antibodies (Fig. 2f). Lower molecular weight bands for Gfap were predominant in CNTF and IGF-1 conditions and this correlated with the observation of long filamentous morphologies by ICC under the same conditions (Fig. 2e). Higher molecular weight Gfap bands were more concentrated in SPONT and 1% FBS and correlated with weaker Gfap staining by ICC (Fig. 2e,f). Despite displaying similar Gfap bands, FBS treated cells were flat and polygonal shaped whereas SPONT had long filamentous morphologies (Fig. 2e,f). RiboTag expression was maintained in every cell across all differentiation conditions (Fig. 2e,f). Some, but very few, Tuj-1 positive cells were detected in differentiation conditions but often colocalized with Gfap (Supplementary Fig. 4c).

Transcriptome differences in astrocyte phenotypes for the differentiation conditions were evaluated using the top 2000 most differently regulated DEGs and categorized using gene panels defined for healthy, reactive and myofibroblast-like (or epithelial to mesenchymal transition (EMT)) astrocyte states that were generated by meta-analysis of published data sets (Fig. 2g, Supplementary Fig 6). Among the top 2000 most different DEGs for the differentiation conditions, Astrocyte (78/429), Reactivity (40/170) and EMT (49/197) genes were identified. All 4 conditions showed upregulation of various healthy astrocyte genes compared with NPC, but only CNTF and 1% FBS evoked net upregulation of reactivity genes with CNTF causing the largest upregulation (Fig. 2h, Supplementary Fig 4d). Only 1% FBS had a net upregulation of EMT genes. CNTF and FBS conditions stained positively for astrocyte reactivity markers Cd44 and Vimentin whereas SPONT did not (Supplementary Fig. 4e).

To evaluate the relative proportions and phenotypes of astrocyte and oligodendrocyte lineage cells, we performed single-nucleus RNA-Seq on NPC, SPONT, and CNTF-directed differentiation *in vitro*. 16,091 total nuclei from the 3 conditions were separated into 11 clusters by Louvain algorithms (Fig. 2i,j, Supplementary Fig. 4f,g). NPC separated into 4 clusters defined by shared elevated expression of *Igfbp2*, a canonical neural stem cell gene^43^ and other canonical NPC genes (Fig. 2k, Supplementary Fig. 4h), with variation in cell cycle/proliferation gene expression prompting the 4 cluster segregation (Supplementary Fig. 4i). SPONT and CNTF conditions induced comprehensive differentiation with only 0.32% and 0.29% of SPONT and CNTF treated cells partitioning into NPC clusters (Supplementary Fig. 4f,g). SPONT and CNTF conditions generated oligodendrocyte (*Tnr* high) and astrocyte (*Gfap high*) lineage cells (Fig. 2k,l, Supplementary Fig. 4g,j) but no obvious neuronal populations. Oligodendrocyte lineage cells segregated into 3 clusters: (i) oligodendrocyte progenitor cells (OPCs) (Cluster 3), (ii) mature oligodendrocytes (Cluster 8), and (iii) a smaller number of actively proliferating OPCs (Cluster 9), and SPONT and CNTF promoted comparable distributions of these different oligodendrocyte clusters (Fig. 2j,k, Supplementary Fig. 4k,l). Astrocyte lineage cells partitioned into four clusters: immature astrocytes (Cluster 1), and more mature astrocytes (clusters 4, 6 and 10) (Supplementary Fig. 4M). SPONT and CNTF promoted different predominant astrocyte clusters such that SPONT comprised ∼90% of total cluster 4 and CNTF comprised ∼93% of total cluster 6 astrocytes (Fig. 2i, Supplementary Fig. 4f,k). Notably, cluster 6 astrocytes, when compared with cluster 4 astrocytes, exhibited higher expression of diverse reactivity genes from the curated reactivity gene list (Fig. 2m, Supplementary Fig. 4m) and cluster 4 astrocytes exhibited comparatively higher expression of canonical healthy astrocyte genes. Moreover, there was strong correlation (r=0.88) between the DEGs for cluster 6 referenced to cluster 4 astrocytes and the Bulk RiboTag DEGs for CNTF relative to SPONT suggesting that the signature that emerged in the bulk analysis was likely associated with the difference in the predominant astrocyte cell population and not just a global upregulation of those reactivity genes across all cells (Fig. 2n).

These data show that upon mitogen removal, RiboTag-NPC cell autonomously differentiated *in vitro* into a mixed population of astrocyte and oligodendrocyte lineage cells, with few neuronal lineage cells, and that non-cell autonomous molecular factors alters the transcriptome phenotypes in a manner consistent with previous observations^44-48^. The expression of astrocyte specific genes, while upregulated compared to NPC, generally did not reach levels of host astrocytes *in vivo* and certain specialized astrocyte genes associated with maintaining neuronal functions were not significantly expressed in cultures.

### Serum exposure induces concentration dependent myofibroblast-like phenotypes in NPC in vitro

NPC transplanted into CNS lesions are exposed to lesion derived molecular cues that can alter cell fate^33^. Acute CNS lesions have substantial blood-brain barrier leak creating microenvironments with spatially varied concentrations of blood and serum-derived molecules^49^. We therefore evaluated the effects on NPC phenotypes of adding increasing concentrations of serum into media during spontaneous differentiation *in vitro*.

We first tested fetal bovine serum (FBS), a widely used media supplement known to influence astrocyte phenotypes *in vitro* compared to serum-free media ^45,48,50^ and found that FBS induced pronounced concentration dependent changes in the transcriptome profiles of NPC undergoing spontaneous differentiation after mitogen withdrawal (Fig. 3a,b). Notably, PCA on 10,427 DEGs revealed increasing divergence from the SPONT state for higher concentrations of FBS and analysis of healthy, reactive and EMT astrocyte gene panels revealed that FBS concentration dependent effects involved adoption of new cell fates rather than attenuation of NPC differentiation such that a low dose (1%) induced the most pronounced astrocyte phenotype, and increasing serum concentrations evoking augmented myofibroblast-like phenotypes often associated with EMT, together with a concurrent decrease in healthy astrocyte genes (Fig. 3b-e, Supplementary Fig. 7a-c). FBS concentration dependent changes in NPC differentiation were also identified by ICC and WB such that Gfap protein expression decreased at higher FBS concentrations, while α-Sma and F-actin (Phalloidin) markedly increased and cells exhibited increases in cytoplasm and nuclei size, with a flattened morphology and augmented arrangement of actin stress fibers, in a manner consistent with a contractile activated myofibroblast-like state (Fig. 3e,g, Supplementary Fig. 7d)^51^.

**Fig. 3:**
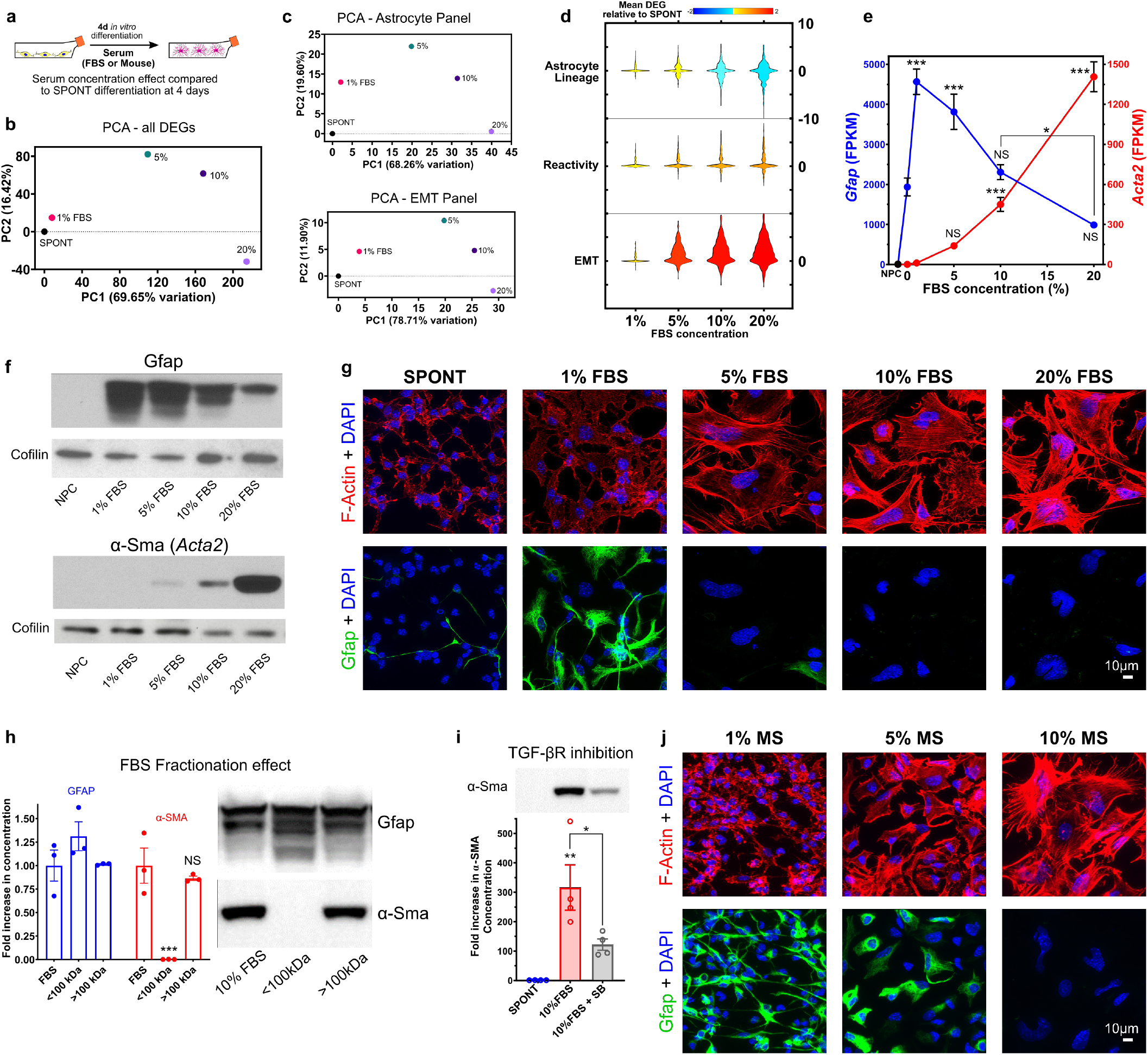
Serum exposure induces concentration dependent myofibroblast-like phenotypes in NPC *in vitro*. **a** Schematic summarizing experimental approach for evaluating serum (Fetal Bovine Serum (FBS) and Mouse Serum (MS)) exposure effects on NPC differentiation *in vitro* by, ICC, WB and Bulk RiboTag. **b** PCA of all DEGs associated with exposure to increasing concentrations of FBS indicating a strong FBS concentration dependent divergence away from SPONT differentiation. **c** PCA of DEGs from healthy Astrocyte and EMT gene panels for FBS treated NPC showing comparable FBS concentration dependent divergence away from SPONT differentiation as observed in b). **d** Violin plot of DEGs for astrocyte state gene panels for FBS treated NPC showing a prominent FBS dependent increase in EMT gene expression and a concurrent decrease in healthy astrocyte gene expression. **e** FBS concentration dependent changes in Gfap and Acta2 gene expression measured in FPKM. N=4 per group *** p-value < 0.0001, * p-value < 0.02, NS = Not significant. One way ANOVA with Tukey multiple comparison test for Gfap and Acta2 run independently, comparisons made with SPONT unless indicated. **f** Western blot (WB) for Gfap and α-Sma showing FBS concentration dependent expression changes. Gfap protein expression decreases with higher FBS concentrations and is associated with loss of lower molecular weight isoform bands. α-Sma protein expression increases with FBS concentration exponentially. Cofilin used as WB and protein concentration reference. **g** ICC staining of FBS treated NPC for Phalloidin (F-Actin), which shows myofibroblast-like phenotype, and Gfap. ICC staining shows similar FBS concentration dependent effects as in e) and f). **h** Quantification of WB for Gfap and α-Sma for FBS and fractionated FBS treatment showing higher molecular weight species (>100 kDa) in FBS stimulated α-Sma expression. **i** Quantification of WB for FBS induced α-Sma expression following TGF-βR inhibition by SB431542 (SB). Comparisons made between SPONT and 10% FBS with and without concurrent TGF-βR inhibition by SB. **j** ICC staining of Mouse serum (MS) treated NPC for Phalloidin (F-Actin) and Gfap showing a conserved serum dependent effect on myofibroblast-like phenotype with increased serum concentration exposure.

To begin identifying the molecular cues in FBS responsible for inducing EMT in NPC, we fractionated FBS by molecular weight. Removing high molecular weight molecules from FBS via 100 kDa molecular weight cut off (MWCO) ultrafiltration attenuated α-Sma expression and increased Gfap expression (Fig. 3h, Supplementary Fig. 7e,f). Treating cells with only the higher molecular weight components of FBS retained after ultracentrifugation induced α-Sma expression comparable to that for the equivalent concentration of FBS (Fig. 3h, Supplementary Fig. 7e,f). Treating NPC with an established EMT inhibitor, the Tgf-β receptor inhibitor SB-431542^52^, significantly reduced α-Sma expression at 10% FBS suggesting that FBS-induced EMT may involve this pathway (Fig. 3i)^53^. CNTF combined with FBS also attenuated the upregulation of EMT genes and α-Sma protein expression while sustaining the full complement of Gfap isoforms including isoforms induced uniquely by FBS or CNTF alone (Supplementary Fig. 7g,h). We also treated NPC with freshly prepared mouse serum, which in high concentrations induced a comparable phenotype to FBS, including reduction in Gfap expression, increased cytoplasm and nuclei size, flattened morphology, and increased arrangement of actin stress fibers (Fig. 3j, Supplementary Fig. 7i), ruling out non-specific effects of FBS.

These data demonstrate that serum derived factors that are present in CNS lesions can non-cell autonomously modify NPC in a concentration dependent manner such that low serum concentrations promote astrocyte differentiation, but high concentrations drive NPC towards myofibroblast phenotypes via EMT-like processes. We found that this effect was associated with the serum fraction with molecular weights greater than 100 kDa that includes candidate molecules such as globulins, transferrin, Complement proteins and assembled lipoproteins. We also identified small molecule EMT inhibitors that can attenuate the EMT-like phenotype in NPC *in vitro*.

### NPC grafted into healthy mouse striatum differentiate into astrocyte and oligodendroglia lineage cells

To assess NPC transplantation outcomes *in vivo* we first grafted NPC into healthy mouse striatum. As carriers, we used two different formulations of our previously developed synthetic polypeptide hydrogel (DCHMO) that differed in polymer weight fraction and mechanical stiffness^35^, and compared these with grafting in media only (Fig. 4a). Cells prepared in media at a density of 100K cells/μL showed an increased tendency to clump and separate in the pipette over the course of an injection session whereas both hydrogel groups maintained uniform suspensions of cells throughout injections. NPC transplanted in media showed reduced cell survival compared to the 2.5% hydrogel but comparable survival numbers to the 5% hydrogel (Fig. 4b).

**Fig 4:**
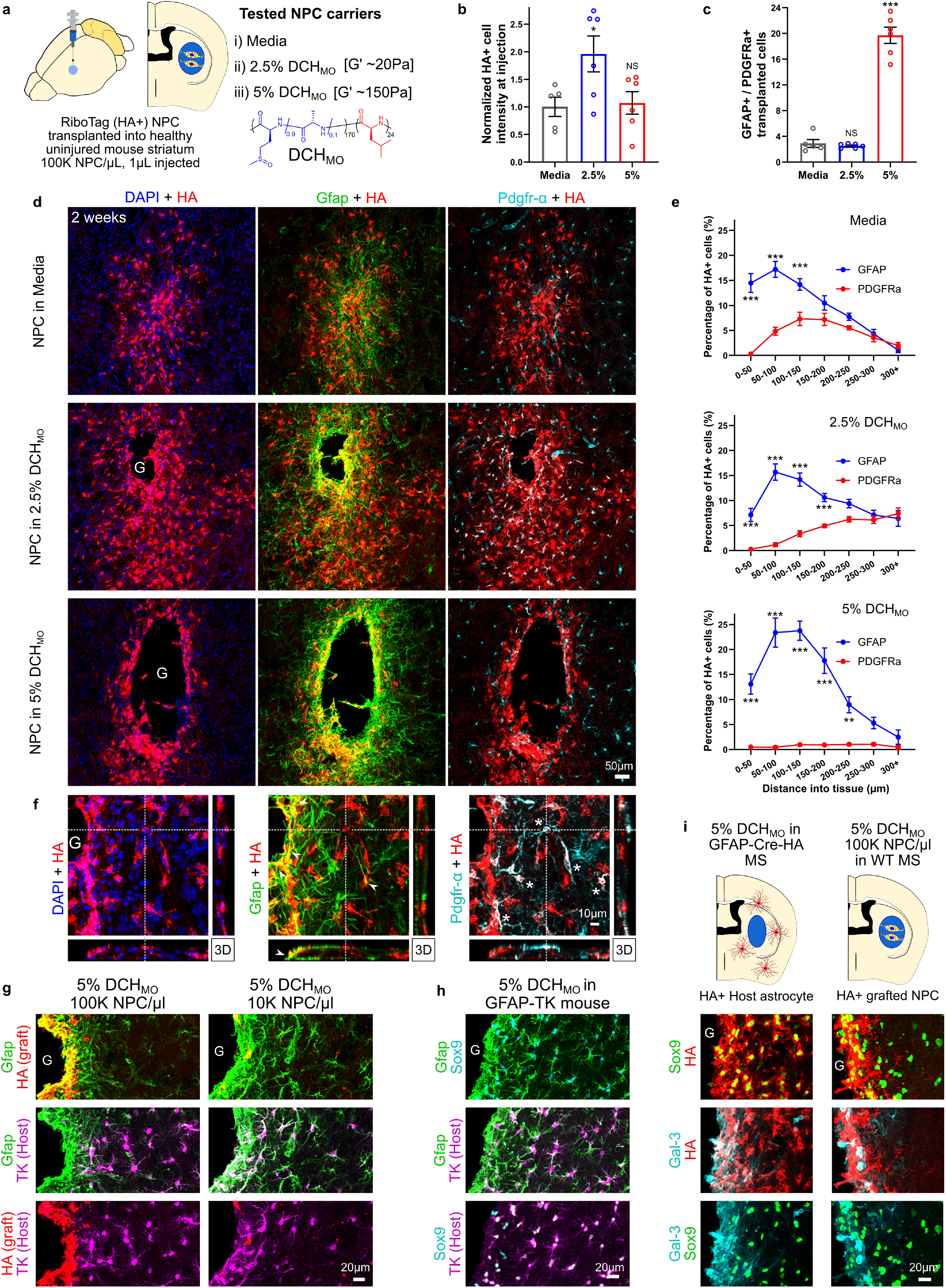
NPC grafted into healthy mouse striatum differentiate into astrocyte and oligodendroglia lineage cells in a spatial and carrier dependent manner. **a** Schematic summarizing experimental approach for evaluating NPC grafting in uninjured mouse striatum and evaluating different hydrogel (DCHMO) transplant carrier formulations. **b** Quantification of normalized NPC number at striatal injection site by HA-positive cell intensity on IHC sections. (N=5 for Media, N=6 for hydrogels). * p-value < 0.05, NS = Not significant. One way ANOVA with Tukey multiple comparison test. **c** Quantification of proportions of Gfap-positive to Pdgfr-α-positive (Gfap+/Pdgfr-α+ ratio) transplanted HA-positive cells (N=5 for Media, N=6 for hydrogels). *** p-value < 0.0001, NS = Not significant. One way ANOVA with Tukey multiple comparison test. **d** Survey Immunohistochemistry (IHC) images of grafted HA-positive NPC in healthy striatum stained for astrocytes (Gfap) and oligodendroglia lineage (Pdgfr-α) at 2 weeks after NPC transplantation using media or hydrogel (G) carriers. **e** Quantification of the percentage of Gfap-and Pdgfr-α-positive grafted cells as a function of distance into neural parenchymal tissue for each carrier. (N=5 for Media, N=6 for hydrogels). *** p-value < 0.0001, **p-value < 0.002. Two way ANOVA with Bonferroni multiple comparison test. **f** High magnification 3D view showing HA-positive grafted NPC in 2.5% DCHMO hydrogel (G) carrier showing formation of Gfap-positive borders (➤) and Pdgfrα-positive cells in neural parenchyma (*) by grafted NPC. **g** Detailed IHC images showing the contributions of host astrocytes (TK-positive) and grafted NPC (HA) to the astrocyte border formed at the NPC hydrogel carrier-neural tissue interface. Contribution from NPC to the astrocyte border is dependent on concentration of cells grafted. **h** Detailed IHC images showing the contributions of host astrocytes (TK-positive) to astrocyte borders formed at hydrogel-neural tissue interface for hydrogel injection alone. **i** Schematic and detailed IHC images comparing HA positive cell responses from host or NPC graft to hydrogel injections. HA-positive host astrocytes derived using Gfap-Cre-HA transgenic mice show prominent HA-positive borders expressing Sox9 and Galectin-3 (Gal-3) at hydrogels injected without NPC. Transplanting HA-positive NPC using the same hydrogel carrier provokes astrocyte border formation by grafted NPC and comparable expression of Sox9 and Galectin-3 (Gal-3).

Grafting vehicle properties markedly influenced the spatial distribution and differentiation phenotype of transplanted cells (Fig. 4c,d). Hydrogel carriers augmented the dispersal of grafted cells over a larger tissue volume compared to media and this effect was influenced by hydrogel formulation (Supplementary Fig. 8a,b). For all conditions, NPC differentiated into a mix of Gfap and Pdgfr-α positive cells at two weeks after transplantation. Notably, graft-derived progeny residing closest to the injection epicenter predominately expressed Gfap whereas Pdgfr-α positive cells were more prevalent in neural tissue several hundred μm from the injection center (Fig. 4d, Supplementary Fig. 8c,d). The 2.5% hydrogel and media showed a similar number ratio of Gfap/Pdgfr-α positive cells of between 2.5-2.9, and remarkably, this ratio was similar to the proportions of astrocyte and oligodendrocyte lineage cells characterized by single cell transcriptome analysis of spontaneous NPC differentiation *in vitro*. In contrast, the stiffer 5% hydrogel carrier showed a predominant Gfap-positive cell phenotype with a Gfap/Pdgfr-α positive cells ratio of ∼20 (Fig. 4c). No neuronal differentiation was observed in any of the animals across the different transplant groups, although grafted NPC interacted directly with healthy neurons (Supplementary Fig. 8e).

Graft derived cells that had migrated into neural tissue tended to adopt individual cell domains reminiscent of host glia (Fig. 4e). Graft-derived cells that persisted at the injection site adopted border forming astrocyte phenotypes similar to those of host astrocytes at interfaces between host neural tissue and biomaterials^1^. This border-forming glia limitans-like phenotype was most apparent around the 5% hydrogel, which formed larger and stiffer deposits. Notably, when present in sufficient numbers, grafted cells assumed most of this border-forming role as demonstrated by using a GFAP-TK transgenic reporter to label host astrocytes, whereas at lower NPC concentrations, host cells predominated at these borders (Fig. 4g). Grafted cells forming these astrocyte limitans borders expressed molecular markers, Gfap, galectin-3 (Gal-3) and fibronectin (Fn), in a manner comparable to host border-forming astrocytes (Fig. 4f-h, Supplementary Fig. 8f).

To assess effects of the transplantation injection on NPC fate, we performed RiboTag RNA-Seq analysis on (i) NPC passed through glass pipettes in media or 2.5% hydrogel, and (ii) cells allowed to dwell in hydrogel for 4 hours before injection, the maximum potential time course of an injection session (Supplementary Fig. 9a). Compared to NPC in culture, there were small but detectable differences in the transcriptome for samples passed through pipettes and most of these changes were shared between these three conditions (carrier and time independent) (Supplementary Fig. 9b-c).

These data show that hydrogel carrier formulation and mechanical properties act as non-cell autonomous modifying cues on NPC, which influenced survival and differentiation outcomes for transplanted cells. The mechanically stiffer hydrogel stimulated a predominant glia limitans-like border phenotype consistent with transplanted NPC responding to the hydrogel as a foreign body and contributing to host astroglial border formation associated with the classic CNS foreign body response^1^. The hydrogel carrier with lower mechanical stiffness but comparable chemistry showed the best NPC survival and interestingly, using this carrier the relative proportions of astrocyte and oligodendrocyte lineage cells detected after NPC grafting into healthy, uninjured tissue were similar to the proportions identified by single nucleus transcriptome analysis of spontaneous NPC differentiation *in vitro*. This optimal hydrogel formulation was used for subsequent experiments.

### NPC grafted into striatal stroke or SCI generate astroglia that reduce fibrotic scar and bridge lesions

To examine NPC transplanted into CNS injuries, we grafted NPC into either L-NIO induced stroke lesions in the striatum or into crush spinal cord injury (SCI) (Fig. 5,6). These models generate two common types of CNS lesions, ischemic lesions due to loss of blood flow after stroke or hemorrhagic lesions caused by severe traumatic injury. NPC were grafted using the 2.5% hydrogel carrier and were injected at two days after either injury (Fig. 5a,6a), a time point after acute CNS injuries when most neural cell death has occurred and multicellular repair processes such as debris clearance and glial proliferation have begun, including the onset of host astrocyte proliferation to form protective borders that separate fibrotic scar from adjacent viable neural tissue (Supplementary Fig. 10a)^4,10,54^. Using separate cohorts of mice, we compared host-astrocytes transgenically labeled with RiboTag via Gfap-Cre, with grafted RiboTag-NPC (Fig. 5, Supplementary Fig. 9c-e). Host astrocytes expressing transgene-derived RiboTag-HA under Gfap-Cre were readily detected in borders around stroke lesions by IHC staining for HA (Fig. 5b), indicating that the HA expressed by NPC-derived cells should also be detectable via HA IHC and that death of HA-positive cells does not result in non-specific HA deposits.

**Fig 5:**
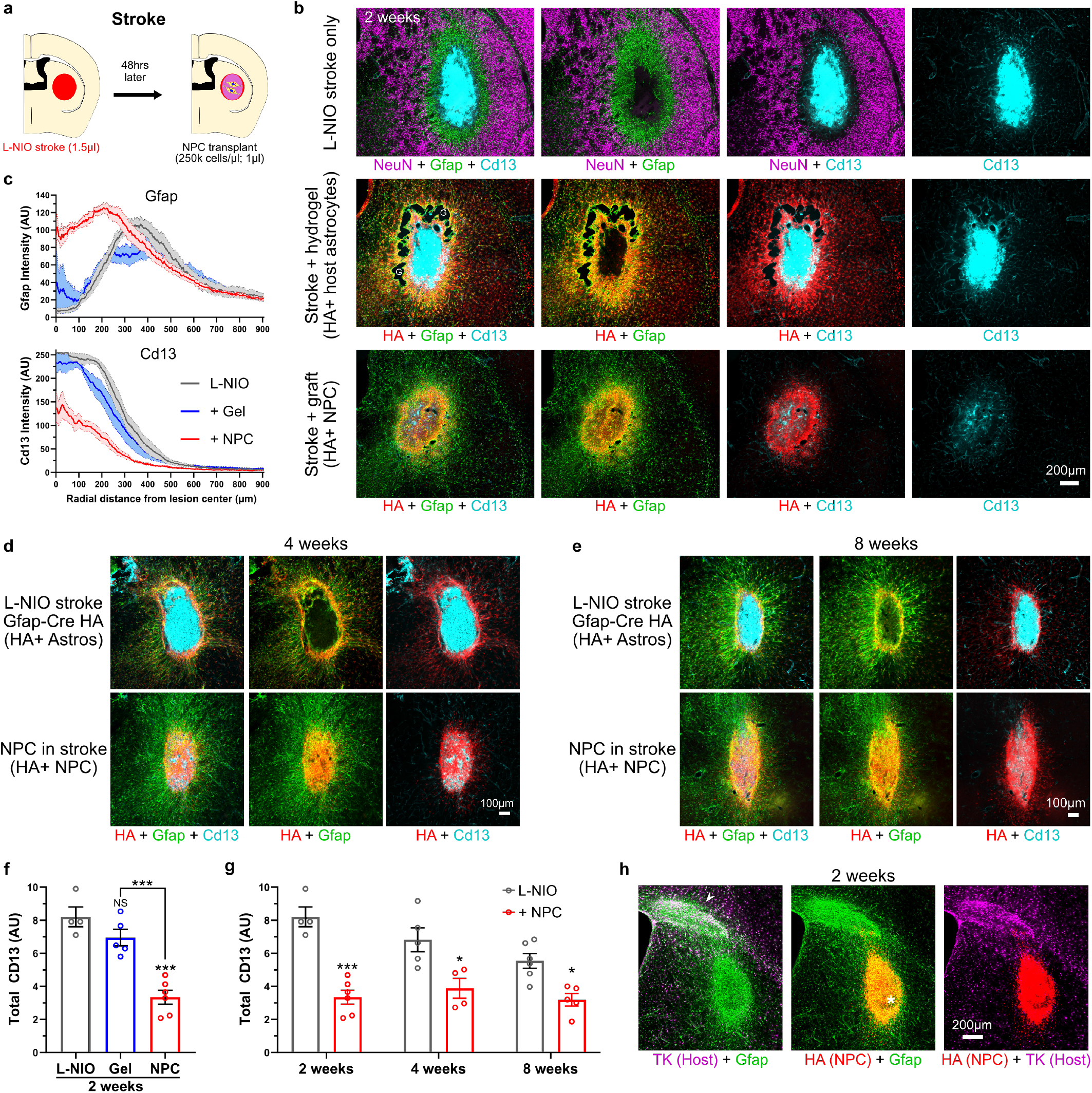
NPC grafted into striatal stroke generate astroglia that reduce fibrotic scar and bridge lesions. **a** Schematic summarizing experimental approach for evaluating NPC grafting in mouse striatal stroke. **b** Survey IIHC images of stroke lesions at 2 weeks comparing effect of hydrogel (DCHMO) alone or grafted HA-positive NPC on stroke lesion phenotype. NPC grafted lesions show Gfap-positive lesions with reduced Cd13-positive tissue in non-neural lesion core. **c** Quantification of Gfap and Cd13 intensity at stroke lesions as a function of the radial distanced from the center of the lesion. **d & e** Survey IHC images of stroke lesions at 4 weeks (**d**) and 8 weeks (e) comparing the effect of grafted HA-positive NPC on stroke lesion phenotype. **f** Quantification of total Cd13 at stroke lesion cores at 2 weeks after L-NIO injection for stroke only, hydrogel (DCHMO) alone, and NPC grafting. (N=4 for L-NIO only, N=5 for gel, N=6 NPC graft). *** p-value < 0.0005. One way ANOVA with Tukey multiple comparison test. **g** Quantification of total Cd13 at stroke lesion cores at 2, 4 and 8 weeks after L-NIO injection for stroke only and NPC grafting. (N=4, 5, 6 for L-NIO only at 2, 4 and 8 weeks, N=6, 4, 5 for NPC grafts at 2, 4 and 8 weeks). *** p-value < 0.0001, * p-value < 0.05. Two way ANOVA with Tukey multiple comparison test. **h** Survey IHC images of NPC grafted stroke lesions at 2 weeks in GFAP-TK mice showing NPC graft contribution to stroke lesion remodeling (*). Host astrocytes have minimal contribution to stroke lesion remodeling but have responded to tissue disruption from pipette insertion (➤) located away from the lesion.

**Fig 6:**
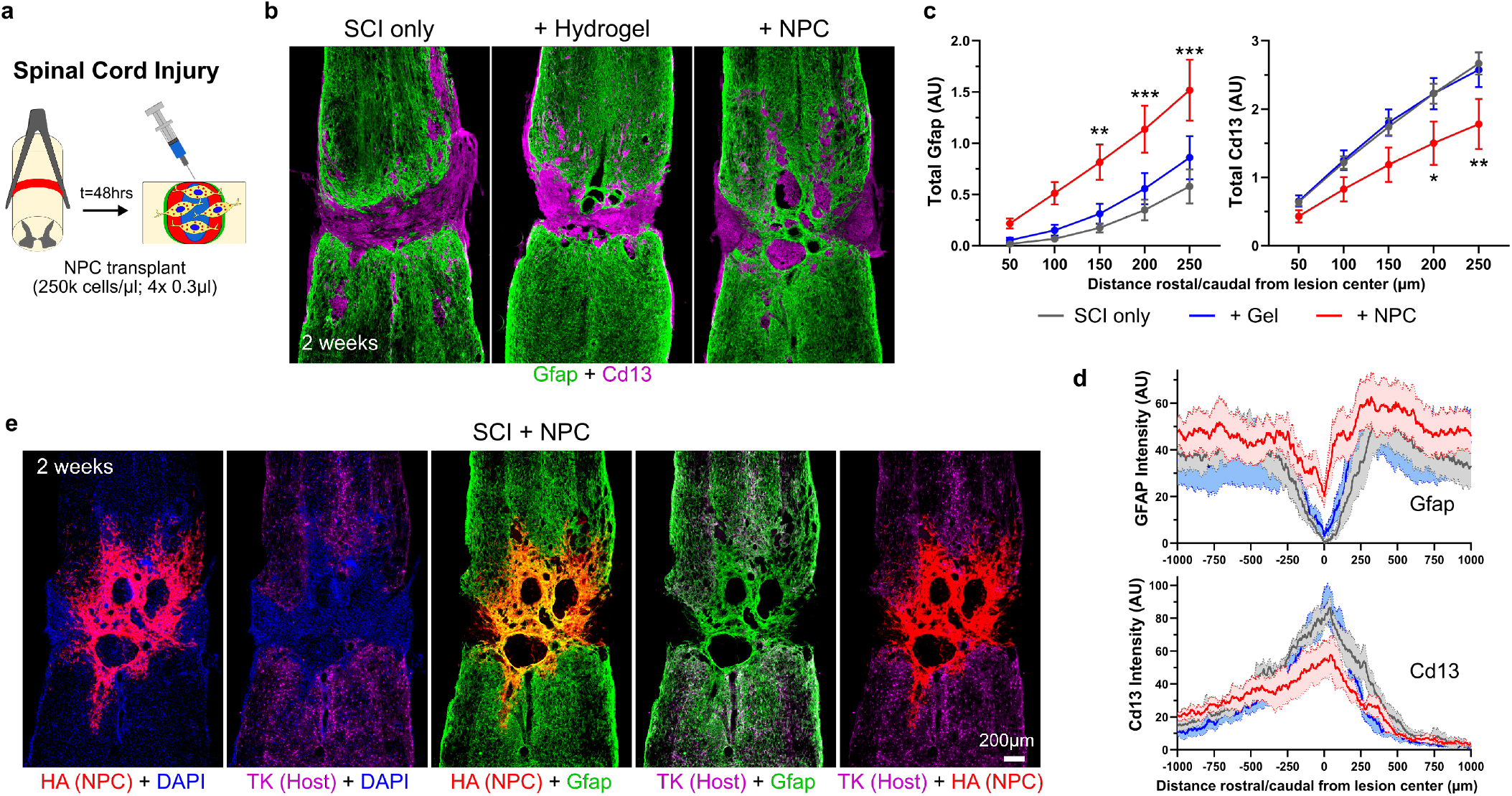
NPC grafted into SCI generate astroglia that reduce fibrotic scar and bridge lesions. **a** Schematic summarizing experimental approach for evaluating NPC grafting in mouse spinal cord injury (SCI). **b** Survey IHC images of SCI lesions at 2 weeks comparing effect of hydrogel alone or grafted HA-positive NPC on SCI phenotype. NPC grafted lesions show Gfap-positive bridges with reduced Cd13-positive tissue in lesion cores. **c** Quantification of total Cd13 at SCI lesion cores at 2 weeks showing increased Gfap and decreased Cd13 at lesion cores, N=6 for SCI only, N=5 for gel, N=6 NPC graft, *** p-value < 0.0005, ** p-value < 0.006, * p-value < 0.05. Two way ANOVA with Tukey multiple comparison test. Gfap and Cd13 intensity plots show mean as darken lines and SEM as shaded area. **d** Survey IHC images of NPC grafted SCI lesions at 2 weeks in GFAP-TK mice showing contribution of grafted NPC to the Gfap-positive tissue at the lesion core.

At two days after L-NIO injection, untreated stroke lesion cores exhibited complete loss of viable neurons and glia, accumulation of axon damage markers including amyloid precursor protein (App), pronounced BBB leak and onset of acute inflammation (Supplementary Fig. 10a). App breakdown into Aβ demarcated the nascent lesion border at this timepoint, suggesting a potential role for Aβ in normal wound responses (Supplementary Fig. 10a). NPC grafts were placed directly into the center of stroke lesions and tissue was harvested after 2, 4 and 8 weeks (Fig. 5b-e). By 2 weeks, untreated L-NIO strokes formed focal compartmentalized lesions with a dense core of Cd13-postive non-neural cells consisting mainly of infiltrating macrophages and fibrotic scar that was surrounded by a border of newly proliferated astrocytes, which isolated the lesion from immediately adjacent viable neural tissue^1^ (Fig. 5b). Injection of empty DCHMO hydrogel into stroke lesions caused no significant alteration to this lesion phenotype (Fig. 5b). In contrast, NPC grafts survived robustly to markedly alter the cellular organization of stroke lesions such that by two weeks, HA-positive grafted cells that were strongly Gfap-positive were widespread throughout the entire lesion core (Fig. 5b). Lesion remodeling facilitated by NPC grafts resulted in significant reduction in total number of Cd13-positive cells and fibrotic lesion size (Fig. 5b-g) which is consistent with the anti-inflammatory and neuroprotective roles performed by host reactive astrocytes^5,10,55^. Grafted HA-positive cells persisted robustly for the duration of all experiments and were abundantly present at 4 and 8 weeks after grafting resulting in a long lasting reduction of infiltrating Cd13-positive cells (Fig. 5d,e,g).

Interestingly, ischemic stroke lesions with NPC grafts lacked the defined non-neural/neural compartmentalization characteristic of untreated stroke and instead exhibited a contiguous bridge of Gfap-positive tissue that filled the entire lesion core (Fig. 5 b,d,e,h). To examine this phenomenon further, we grafted NPC into transgenic mice expressing Gfap-TK in host astrocytes to compare the contributions of host-derived and graft-derived Gfap-positive cells. This comparison revealed that Gfap-positive cells throughout the lesion core were derived exclusively from transplanted cells, which remained entirely within the stroke site (Fig. 5h). NPC grafted into strokes exhibited negligible expression of mature astrocyte markers such as Aldh1l1 at 2 weeks (Supplementary Fig. 10b), but this expression increased by 4 and 8 weeks, particularly in graft-derived cells located at the immediate host neural tissue interface (Supplementary Fig. 10c). NPC treated stroke lesions transiently contained P2yr12-positive CNS derived microglia at 2 weeks, but not at 4 or 8 weeks, which together with the significantly reduced Cd13-staining and visibly reduced Cd68 suggested an NPC graft induced shift in the immune cell milieu within the lesion core (Supplementary Fig. 10d-e). Nevertheless, NPC grafts did not show any neuronal repopulation or ingrowth of neurofilament positive axons by 8 weeks (Supplementary Fig. 10f).

We next examined the effects of NPC grafts into SCI (Fig. 6a). As expected^1,56^, untreated crush SCI caused focal compartmentalized lesions similar to untreated L-NIO stroke, with a dense core of Cd13-postive non-neural cells consisting mainly of infiltrating macrophages and fibrotic scar surrounded by a border of newly proliferated astrocytes (Fig. 6b). SCI treated with DCHMO hydrogel only also exhibited similar discretely compartmentalized lesions with Cd13-positive cores surrounded by host-derived border forming astrocytes (Fig. 6b). NPC transplanted at two days after SCI and evaluated at 2 weeks survived robustly and differentiated into Gfap-positive cells that populated lesion cores and formed contiguous cell clusters and tracts throughout the lesions (Fig. 6b). NPC grafting also significantly reduced total Cd13-postive cells and increased Gfap-positive cells within SCI lesions (Fig. 6c,d), but remodeling of lesions with graft-derived cells was not as comprehensive after SCI as after stroke injury. In stroke, grafted NPC-derived cells essentially filled the entire lesion core and there were few isolated volumes of infiltrating inflammatory cells and fibrosis, whereas in SCI, there were discrete, and sometimes large, pockets of Cd13-positive immune cells surrounded by grafted cell borders (Fig. 6b,e). These differences could be due to differences in lesions generated by ischemic stroke injury with little or no bleeding or by crush SCI with clear evidence of bleeding and pronounced hemorrhagic necrosis^57^. Nevertheless, even after SCI, lesions with NPC grafts exhibited significantly reduced infiltration of peripheral inflammatory cells and reduced amounts of fibrotic tissue (Fig. 5c,d), and there were prominent Gfap-positive cell tracts that surrounded inflammatory and fibrotic cells throughout the lesion core in all animals, and these tracts provided bridges of contiguous Gfap-positive cells that spanned the SCI lesions across their rostral to caudal borders (Fig. 5b,e). Comparisons using separate molecular markers of host (TK) and graft (HA) derived Gfap-positive astroglia in the same animals demonstrated that the bridges spanning the lesion core were comprised exclusively of grafted cells and that the grafted cells integrated seamlessly with host astrocytes (Fig. 5e). Notably, although these NPC-derived astroglial bridges did not encourage significant spontaneous infiltration of neurofilament positive axons, previous studies show that appropriately stimulated and chemoattracted host axons can grow along host astrocytes bridges in lesions^56,58^, suggesting that NPC-derived astroglial bridges warrant future testing for this capacity.

These findings show that NPC grafted into forebrain stroke or thoracic SCI lesions at two days after injury survive well, persist and populate the lesion cores with Gfap-positive cells. Notably, graft-derived Gfap-positive cells spread throughout the lesion cores and significantly reduced the amount of tissue exhibiting inflammation and fibrosis after both stroke and SCI. In addition, grafted NPC-derived cells formed contiguous tracts of Gfap-positive cells that spanned across lesions and integrated seamlessly with host astrocytes, thereby forming astroglial bridges that interconnected preserved neural tissue on opposite sides of lesions after both stroke and SCI.

### Astroglia derived from NPC grafted into subacute SCI mature over time, are altered by non-cell autonomous cues in the lesion environment and exhibit features of naturally occurring border-forming astroglia

Host astrocytes that form borders around CNS lesions and thereby corral infiltrating inflammatory cells and fibrosis are newly proliferated, derive largely from local mature astrocytes, express high levels of Gfap and exhibit pronounced differences in cellular appearance, organization and gene expression compared with mature astrocytes in healthy tissue^5,59,60^. Using IHC, we found that NPC graft-derived cells also expressed high levels of Gfap and organized into contiguous borders along the entire interfaces between viable neural tissue and the stromal and inflammatory cells of lesion cores after both stroke and SCI in a manner similar to host border-forming astrocytes (Fig. 5,6, 7a-c). These graft-derived borders also expressed various additional markers associated with host-derived astrocyte borders including Lcn2, Clu and Cpe (Supplementary Fig. 11a-d). Notably, lesions that received NPC grafts exhibited a reduced contribution of host astrocytes to astrocyte borders and graft-derived astroglia seemed to adopt this role. Moreover, lesions with NPC-derived astroglia exhibited significantly less inflammatory and fibrotic tissue indicating that NPC-derived astroglia shared functional similarities with host border-forming astrocytes and could in some cases substitute for them (Fig. 5, 6, 7c).

**Fig. 7:**
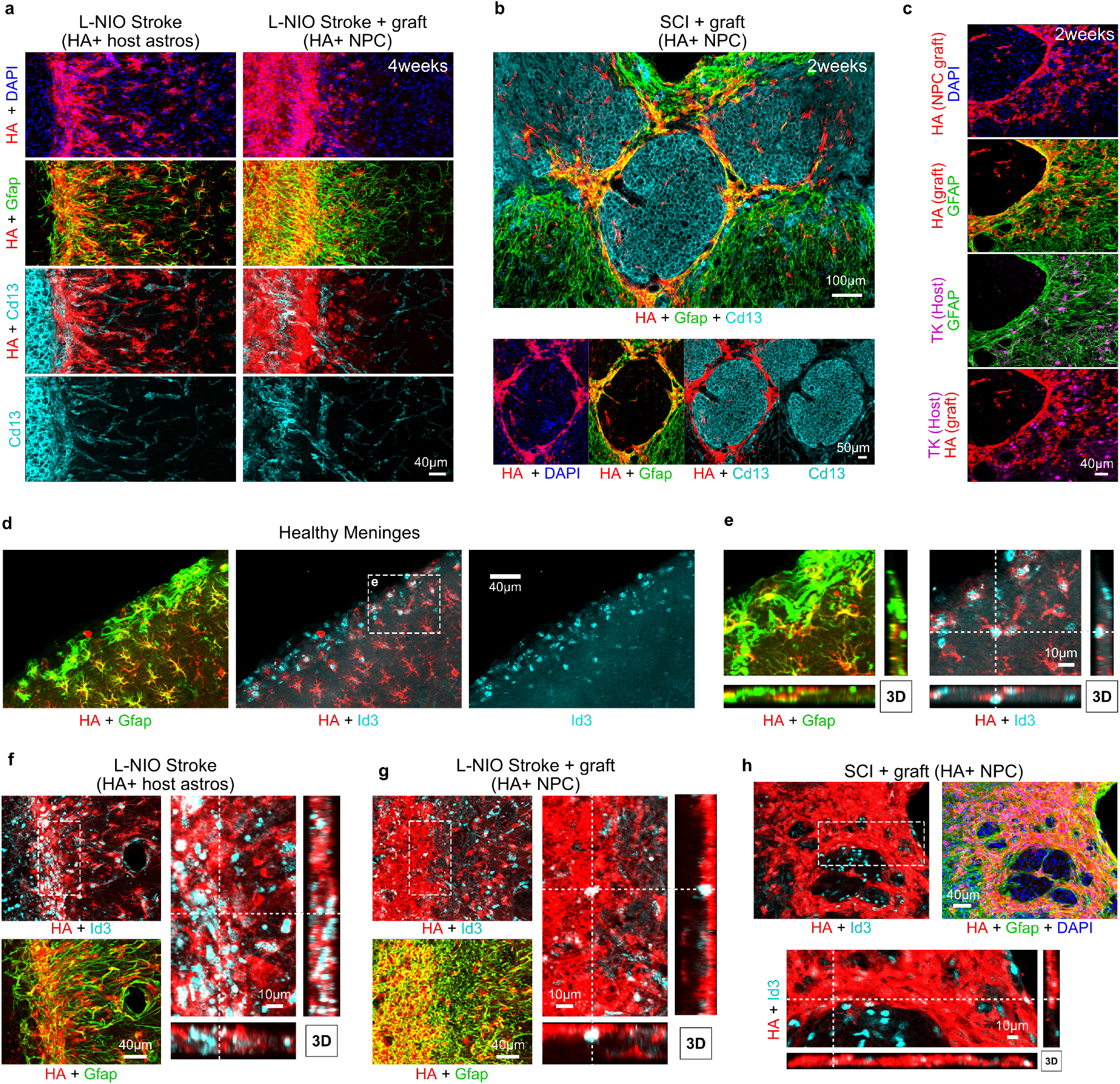
Astroglia derived from NPC grafted into CNS lesions exhibit features of border-forming astroglia. **a** Detailed IHC images showing the astrocyte border phenotype of stroke lesions at 4 weeks in untreated stroke only and NPC grafted strokes. **b** Survey IHC of NPC grafted SCI lesions at 2 week showing NPC derived astrocyte borders formed around Cd13-postive inflammatory cells. **c** Detailed IHC images showing the astrocyte border phenotype in SCI derived from transplanted NPC and not from host astrocytes (TK-positive). **d, e** Survey and detailed, higher magnification 3D IHC imaging of healthy mouse cortex and meninges showing higher expression of transcriptional regulator Id3 (inhibitor of DNA binding 3) in host astrocytes (HA+ Gfap-Cre-RiboTag) that form glia limitans at the meninges and penetrating blood vessels. **f** High magnification 3D IHC images showing astrocyte borders formed at stroke lesions upregulate expression of Id3. **g** High magnification 3D IHC images showing grafted NPC that remodel stroke lesions also express Id3. **h** NPC grafted into SCI lesions that form astrocyte borders also express Id3.

Host astrocytes that form borders around lesions exhibit characteristics and functions similar to glia limitans astrocytes along all meningeal borders in the healthy CNS ^5^. Glia limitans astrocytes that abut the fibroblast lineage cells of the meninges or penetrating blood vessels with meningeal borders selectively express the transcription factor Id3^61^ (Fig. 7d,e, Supplementary Fig. 11e-f). We found that this Id3 expression is shared not only by host border forming astrocytes around lesions, but also by NPC graft-derived astroglia that similarly form such borders (Fig. 7f-h) further supporting similarities between these cell types. In contrast, Id3 expression is sparce among other astrocytes located distant to borders with either meninges, large vessels or injury lesions (Figure 7d-h, Supplementary Fig. 11e-f).

To dissect transcriptional characteristics of grafted NPC *in vivo*, we focused on SCI in order to make direct comparisons with existing datasets on host astrocytes responses to SCI derived using the same RiboTag IP method^56,60^, and for ease of collecting high quality tissue restricted to the lesion site from SCI versus forebrain stroke. The RiboTag procedure enabled recovery of mRNA selectively from small populations of transplanted NPC and their progeny with high yield and specificity (Fig. 8a, Supplementary Fig. 12a,b). NPC grafted at 2 days after SCI and harvested at 14 days showed upregulation of astrocyte, reactivity and EMT genes with a concurrent downregulation of NPC genes when compared with pre-graft NPC profiles (Fig. 8b). Host border-forming astrocytes exhibited a comparable upregulation of reactivity and EMT genes at 14 days after SCI (Fig. 8c), with a downregulation of genes associated with healthy astrocyte functions, consistent with their new primary interactions with non-neural lesion core cells rather than with neural cells. PCA and CS analyses revealed that from very different transcriptional starting states, NPC and host astrocytes at SCI lesions both converged to remarkably similar astroglial reactivity states despite large differences in their overall transcriptional state changes (all genes) (Fig. 8d-l). The gene panel for reactivity in particular showed remarkable congruency in the responses of host astrocytes and NPC with comparable vectors in Euclidean PCA space and a CS = 0.843 (Fig. 8h,i). Reactivity genes such as *Tgm1, Cd22, Serpina3n, Timp1* showed highly correlated elevated expression in both NPC and host astrocytes in SCI (Fig. 8l). Other reactivity genes associated with astrocyte proliferation after injury were inversely correlated because NPC downregulate these genes from high levels upon grafting whereas host cells upregulated them from very low levels after injury (Fig. 8 g,j,l). Restricting transcriptome analysis to the healthy astrocyte gene panel revealed convergent changes whereby NPC and host astrocytes increased or sustained expression of astrocyte genes associated with glia limitans borders ^61^ (e.g. *Gfap, Id3, Vim, Clu, Chil1, Ccn1, C4b*) (Fig. 8k) and reduced expression of genes associated with specialized neuronal support functions such as synapse maintenance (*Sparc, Sparcl*) and molecular transport (*Atp1a2, Slc1a2, Slc6a11, Slc6a1, Slc7a10, Dio2, Agt*) (Fig.8k).

**Fig. 8:**
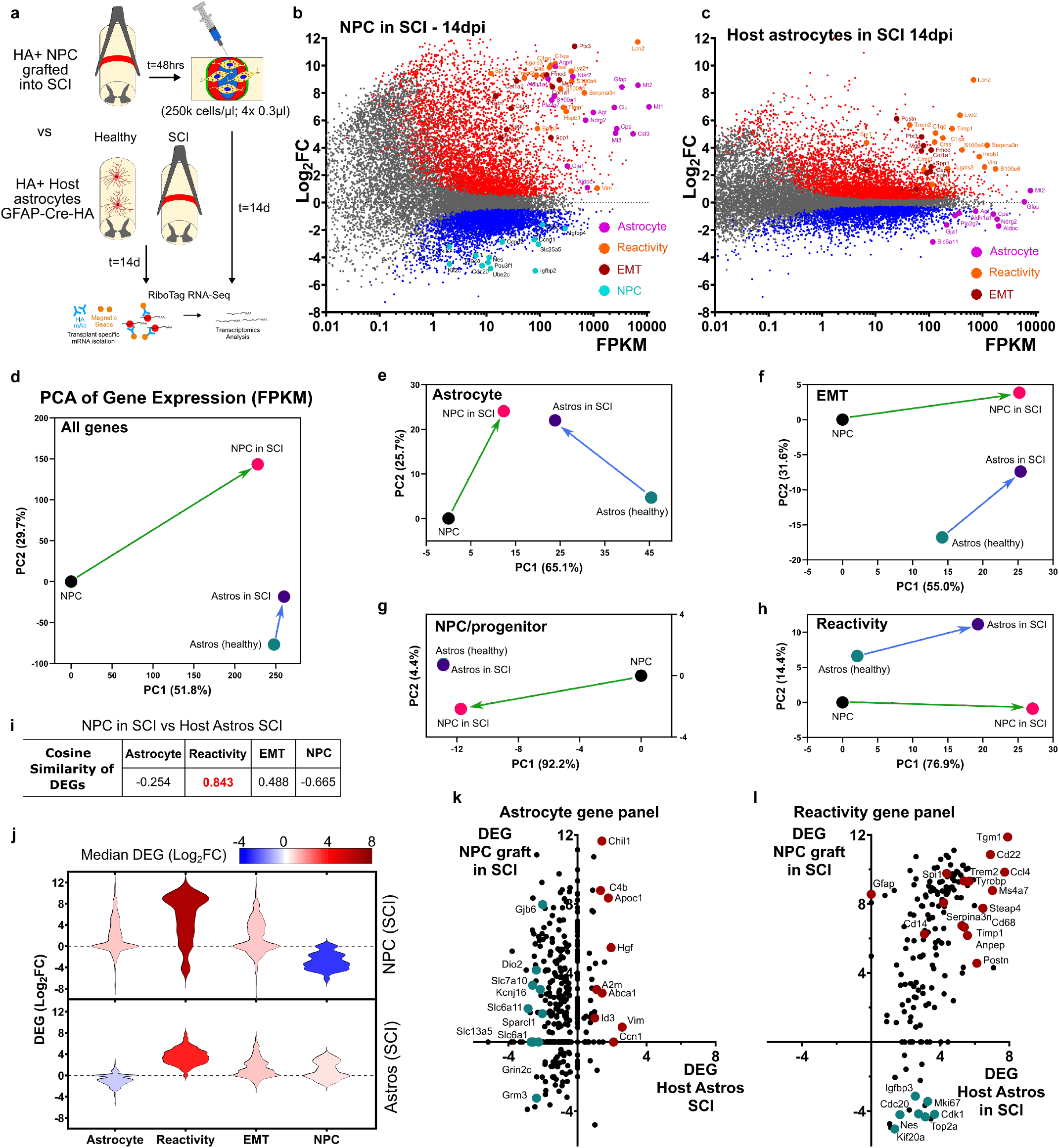
Astroglia derived from NPC grafted into SCI exhibit similar astroglial reactivity states to host astrocytes responding to SCI. **a** Schematic summarizing experimental approach for evaluating transcriptomes of NPC grafted into SCI with host astrocytes responding to SCI. **b** M-A Plot of grafted NPC (N=5 samples) recovered at 2 weeks post injury (14dpi) compared to pre-grafted NPC (N=5 samples) showing increased expression of healthy astrocyte, reactivity and EMT genes and decreased expression of NPC/proliferation genes. **c** MA-Plot of host astrocytes at 14 days after SCI showing comparable upregulation of reactivity and EMT genes but decreased expression of healthy astrocyte genes. **d-h** PCAs using FPKM values comparing state changes between NPC before and after grafting into SCI (recovered at 14dpi) and host astrocytes before and 14 days after SCI for all expressed genes (d) as well as using gene panels for healthy astrocytes (e), EMT (f), progenitors (g) and reactivity genes (h). **i** Cosine similarity of DEGs comparing state changes in NPC after grafting with that of host astrocytes after SCI. NPC and astrocytes show similar reactivity and EMT gene changes but divergent changes in astrocyte and NPC/proliferation genes. **j** Violin plots of DEGs for NPC grafted into NPC and host astrocytes in SCI for astrocyte, reactivity, EMT and NPC/proliferation gene panels. **k**,**l** Plot of DEGs from the astrocyte (k) and reactivity (l) gene panels for NPC after grafting compared with astrocytes after SCI showing strong correlation of reactivity genes and genes associated with glia limitans borders. Genes from the reactivity gene list but that are also associated with proliferation or progenitor phenotype (cyan) such as *Mki67, Top2a*, and *Nes*, show an inverse correlation.

Host astrocyte border formation around CNS lesions is a dynamic process that evolves over time during the 14 days after SCI and is likely to be influenced by molecular cues in the lesion environment^54^. To evaluate NPC graft transcriptome at an earlier timepoint during border formation we harvested grafts at 5 days after transplantation (Fig. 9a-c, Supplementary Fig. 12c). Compared with the NPC grafts recovered at 14 days, grafts at 5 days showed a significantly elevated EMT signature, lower astrocyte characteristics, and similar reactivity and progenitor gene expression profiles (Fig. 9c, Supplementary Fig. 12c,d). Key EMT regulatory genes more highly expressed in 5-day grafts included canonical markers *Snai2, Postn, Fbln2* and *Acta2* (Fig. 9b, Supplementary Fig. 12d). The downregulation of EMT gene expression from 5 to 14 days coincides with consolidation of wound repair functions and stabilization of astroglial borders as well as reduced blood and serum exposure. Concurrently, healthy astrocyte gene expression increased from 5 to 14 days including genes required for forming and maintaining astrocyte border functions such as *Chil1, Clu, C4b* and *Myoc*^61^ as well as specialized astrocyte functional genes such as *Aqp4, Agt, Hepacam, Slc6a1* (Fig. 9b, Supplementary Fig. 12d).

**Fig. 9:**
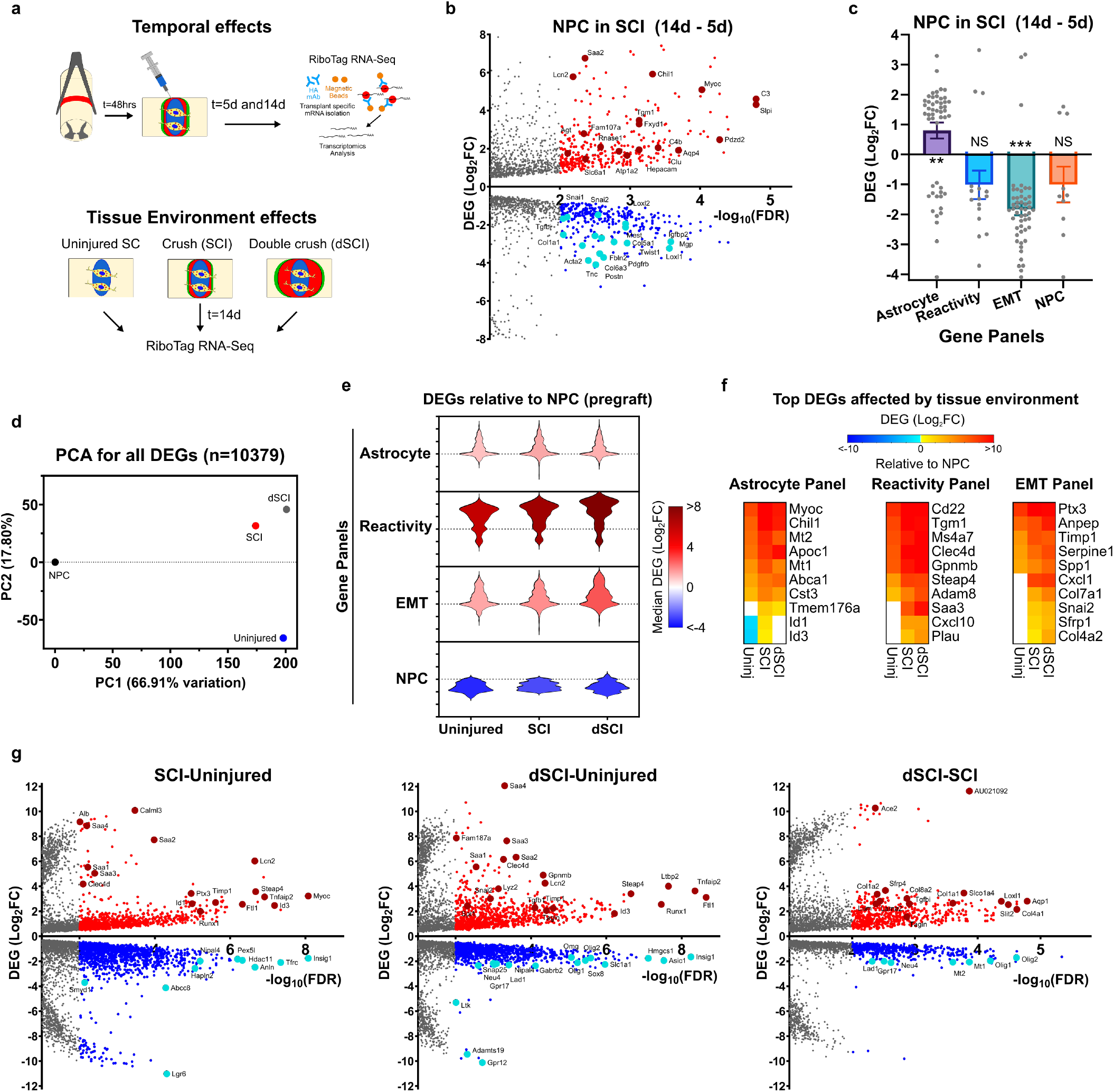
Astroglia derived from NPC grafted into subacute crush spinal cord injury mature over time and are altered non-cell autonomously by the lesion environment. **a** Schematic summarizing experimental approach for evaluating temporal and tissue environment effects of NPC grafting into mouse SCI. **b** Volcano Plot comparing DEGs for NPC grafted into 2d SCI and recovered at 14 days after grafting versus samples recovered at 5 days after grafting. **c** Bar graph of DEGs for astrocyte, reactivity, EMT and NPC/progenitor gene panels for NPC in SCI recovered at 14 days versus 5 days. Astrocyte gene expression in NPC increases whereas EMT gene expression decreases temporally. *** p-value < 0.0005, ** p-value < 0.004, One sample t-test for hypothetical mean = 0. Individual DEGs are overlayed on the graft. **d** PCA for all DEGS for NPC grafted into uninjured, single crush SCI (SCI) and double crush SCI (dSCI) referenced to NPC (pre-graft) showing large differences for all differentiation conditions compared to the NPC state and the variation amongst the unique tissue environments. **e** Violin plots of DEGs for NPC grafted into different tissue environments for astrocyte, reactivity, EMT and NPC/proliferation gene panels. **f** Heat maps showing the top 10 DEGs for the astrocyte, reactivity and EMT gene panels most altered in grafted NPC by tissue environment. **g** Volcano Plot comparing DEGs for NPC grafted into different tissue environments.

Individual grafted NPC phenotypes, as detected by IHC, correlated with the relative spatial proximity to preserved neural tissue or pockets of immune and fibrotic cell laden lesion cores such that NPC within or adjacent to neural tissue expressed Gfap, whereas NPC surrounded only by non-neural cells did not, suggesting a strong environmental influence on NPC functions (Fig. 7b,c). To further dissect the effect of molecular environment on non-cell autonomous regulation of NPC phenotype we grafted NPC into three different conditions in the spinal cord that would direct unique predominant differentiation phenotypes that could be readily evaluated by bulk RiboTag transcriptomics. The three environments were: (i) uninjured spinal cord, (ii) single forceps crush SCI (SCI) as done previously, and (iii) double forceps crush SCI (dSCI) that created substantially larger non-neural cores and decreased the influence on NPC grafts of preserved neural tissue at the margins of the lesion (Fig. 9a, d-g, Supplementary Fig. 12e,f). NPC transcriptome analysis at 14 days revealed both common changes in NPC grafts that were not altered by tissue environment, and lesion environment specific effects (Fig. 9 e-g). Notable astrocyte genes that were upregulated by at least a Log_2_FC>5 upon grafting versus pre-graft NPC gene expression regardless of lesion size included *Gfap, Atp1a2, Pla2g7, S100b, Slc1a2, Slc6a11, Rnase1*. Comparing NPC grafted into SCI versus into uninjured spinal cord tissue (Fig. 9g) showed: (i) a significant increase in expression of astrocyte border genes *Id3, Myoc, Chil1, Vim* consistent with a higher proportion of grafted cells adopting astroglia border functions in response to injury, and (ii) a concurrent significant decrease in oligodendrocyte lineage expression including genes such as *Anln, Hapln2, Pex5l, Nipal4, Gpr37, Plp1, Mag, and Mbp*. Consistent with IHC observations, NPC grafted into more severe injury environments with less exposure to neural tissue (dSCI) exhibited increased expression of reactivity as well as EMT genes (Fig. 9g, Supplementary Fig. 12f). Astrocyte reactivity genes were most altered by tissue environment (Supplementary Fig. 12f) with immune and inflammation response related reactivity genes such as Lyz2, *Cd22, Adam8, Saa3, Steap4, Clec4d*, and *Tgm1* most prominently varied by environment (Fig. 9g).

These data show that grafted NPC differentiation is strongly influenced by non-cell autonomous cues in lesion microenvironments. The findings reveal striking similarities between the transcriptional profiles, cell morphologies and functions of NPC transplanted into CNS lesions to that of host proliferating astroglia that generate a naturally occurring wound repair astroglial phenotype that functions to form borders around CNS lesions.

## Discussion

Directing the fate of transplanted NPC into desired cell phenotypes to serve specific therapeutic purposes after transplantation *in vivo* will require a detailed understanding of the relative roles of non-cell autonomous cues and cell autonomous programs in determining NPC fate. Although NPC can be matured into neuronal or glial restricted progenitors *in vitro* that tend to give rise preferentially to neurons or glia after grafting *in vivo*^33^, the pronounced degree to which different host cell environmental cues modulate NPC differentiation is only beginning to be recognized, specified and addressed^62^. Our findings here demonstrate the power of the Rpl22-HA (RiboTag) tool for transcriptionally profiling and morphologically characterizing NPC and their progeny after grafting under different conditions *in vivo*. Using this approach, we compared the transcriptional states of NPC grafted into uninjured or lesioned CNS tissue with the transcriptional changes induced in NPC by exposure to specific molecular cues *in vitro* and with transcriptional profiles of endogenous CNS wound repair cells. Our findings show that (i) NPC fate is strongly influenced by different non-cell autonomous cues *in vitro* and in different host CNS environments *in vivo*, and (ii) NPC grafted into CNS subacute lesions after stroke or SCI adopt a phenotype similar to newly proliferated host astroglial that participate in wound repair (Fig. 10).

**Fig. 10:**
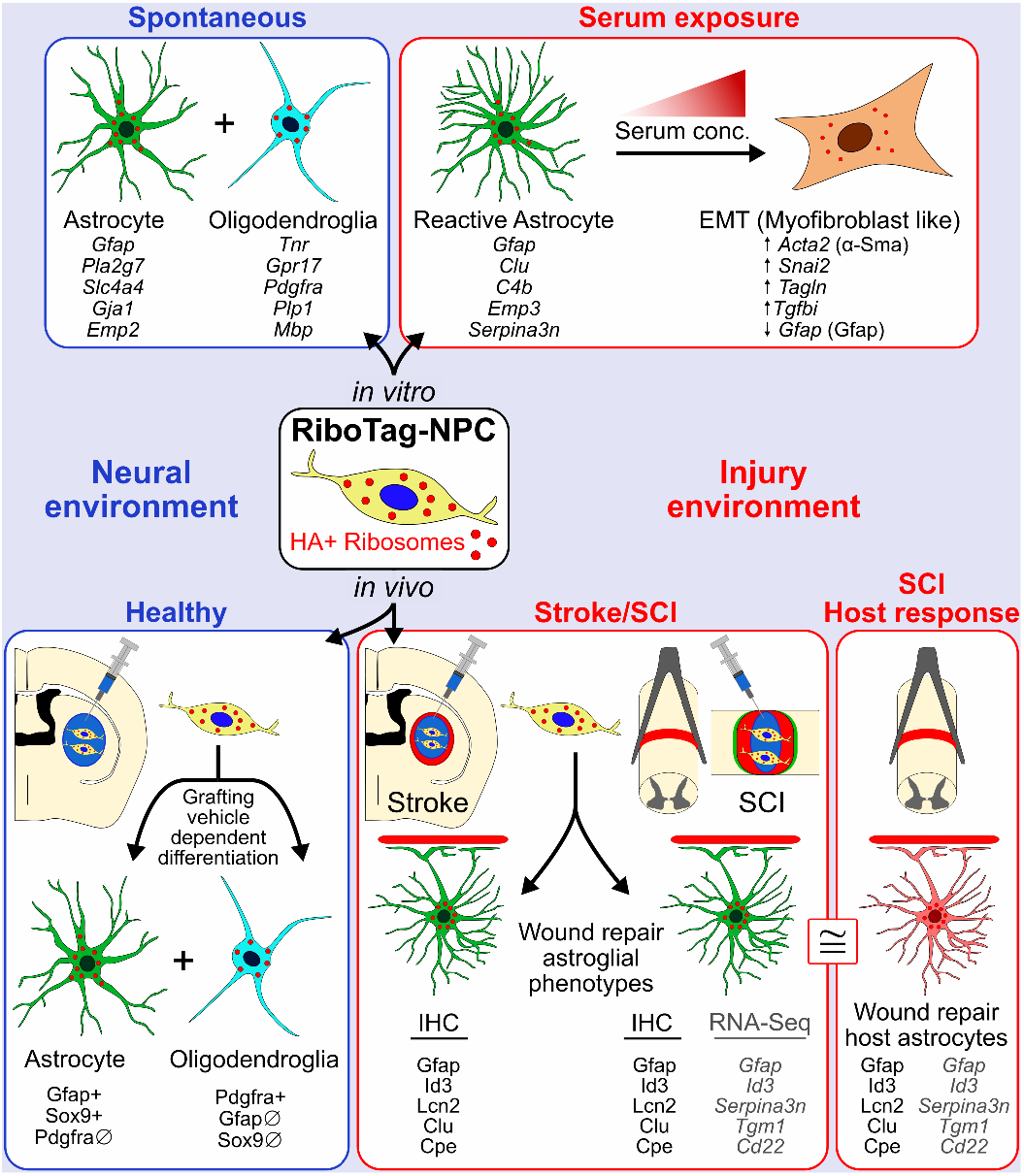
Schematic summarizing the main findings. RiboTag-NPC were derived by neural induction of mESC and were used for RNA-Seq and IHC studies *in vitro* and *in vivo*. When placed into *in vitro* or *in vivo* neural environments, RiboTag-NPC spontaneously differentiated into mixed populations of astrocyte lineage and oligodendrocyte lineage cells. Glial cell differentiation was verified based on gene expression determined by bulk RiboTag RNA-Seq and single nuclei RNA-Seq. Introducing NPC into CNS injury-like environments *in vitro* (via exposure to increasing concentrations of serum) or grafting *in vivo* into models of stroke and spinal cord injury, instructed cells to adopt wound repair astroglial phenotypes through non-cell autonomous cues in a manner consistent with serum concentration and lesion properties (temporal, spatial, mode of injury). High serum concentrations *in vitro*, or grafting into larger and more severe lesions (not depicted), increased myofibroblast-like phenotypes in NPC characteristic of EMT. The neuroprotective wound repair astroglial phenotype adopted by grafted NPC in CNS injuries shared numerous protein and transcriptome similarities (≅) with the host astrocyte wound repair response. Example proteins and genes (*italicized*) shared by these wound repair phenotypes are listed. EMT, epithelial mesenchymal transition; HA, haemagglutinin; IHC, immunohistochemistry.

### Non-cell autonomous cues instruct NPC differentiation in vitro and in vivo

After mitogen withdrawal *in vitro*, RiboTag-NPC spontaneously differentiated primarily into astrocyte and oligodendrocyte lineage cells as reported for NPC by others^36^. This cell autonomous differentiation *in vitro* was modified as expected by known non-cell autonomous cues such that exposure to CNTF or 1% serum generated more astroglia^36,39,40^, whereas IGF1 generated more oligodendroglia^41^. NPC grafted into healthy CNS tissue generated cells with transcriptional phenotypes similar to astroglia and oligodendroglia present in healthy adult CNS tissue or to cells generated after NPC differentiation *in vitro*. Notably, NPC grafted into uninjured CNS tissue intermingled with local neurons and upregulated genes associated with neuronal interactions. In striking contrast, these NPC when grafted into subacute CNS lesions and exposed to serum from the blood-brain barrier leak, differentiated primarily into cells with transcriptional and morphological features similar to newly proliferated host astrocytes that surround lesions and corral inflammatory and fibrotic cells. Notably, NPC progeny did not intermingle with nearby neurons and did not upregulate genes associated with neuronal interactions, but instead interacted with inflammatory and fibrotic tissue and upregulated genes associated with wound repair. To look for potential non-cell autonomous cues underlying these differences in grafted NPC fate, we examined NPC responses to serum proteins *in vitro*. We found that serum proteins at lower concentrations fostered NPC acquisition of astroglial features but at higher concentrations attenuated astroglia features and promoted EMT-like changes. *In vivo*, we noted compatible changes with these *in vitro* findings such that NPC located in the center of large lesions and thereby exposed to high serum and inflammatory/fibrotic cells but not to neural cells exhibited more EMT-like features, whereas NPC located along borders between inflammatory/fibrotic cells and neural cells and thereby exposed to mixed cues from both cell populations were directed towards a border-forming astroglial phenotype, indicating that different micro-environments within lesions can differentially influence NPC cell fate. Thus, non-cell autonomous cues in different host tissue environments can powerfully modulate grafted NPC transcription and instruct differentiation into cells with different phenotypes. This differential responsiveness of NPC to different non-cell autonomous cues after grafting *in vivo* strongly supports the notion that understanding these cues will foster the development of bioengineered interventions that can intentionally provide desired, or block unwanted, cues and thereby direct grafted NPC into therapeutically beneficial phenotypes^62^.

### CNS lesion environments instruct grafted NPC towards a naturally occurring wound repair astroglial phenotype

During naturally occurring wound repair in the adult CNS, neural tissue that has been lost to injury or disease is replaced by non-neural fibrotic tissue that is partitioned away from adjacent viable neural tissue by protective astroglial borders^1-10^. This stereotypic organization of CNS lesions is conserved across rodents and humans, and the newly proliferated astroglia that form narrow borders between neural and non-neural tissue during wound repair share similarities with astroglial that form narrow ‘limitans’ borders separating neural from non-neural tissue along meninges in the healthy CNS^5^.

Here, we found that NPC grafted into subacute CNS lesions differentiated into an astroglial phenotype with striking similarities to that adopted by proliferating host astrocytes that generate naturally occurring astroglial borders around non-neural lesion core tissue. Starting from two very different transcriptional states, transplanted NPC and proliferating host astroglia in serum-exposed lesions exhibited convergent differentiation into astroglia with similar transcriptional, morphological and functional features. In this regard, it deserves mention that the peak time of host-derived astrocyte proliferation for border formation is from 2 to 7 days after injury^54^ so that NPC grafted at 2 days after stroke or SCI, as done here, were placed into an environmental niche potentially primed to direct them into such border forming cells. It is interesting to speculate that NPC and proliferating astrocytes share similar differentiation potentials and that the same molecular cues in lesions might have similar effects on their differentiation. Others have shown that adult astroglia induced to proliferate after injury and then placed into neurogenic culture conditions exhibit differentiation potentials similar to NPC^59,63^, providing further evidence for similarities between these cells. A likely source of differentiation modifying cues is serum-derived molecules that are present in high concentrations in lesions during the timepoint of astroglial proliferation and border formation. We show here that serum exposure *in vitro* alters the transcription of NPC differentiating *in vitro* to become similar to that of reactive astrocytes in CNS lesions. In addition, we show that border forming astroglia derived from either grafted NPC or host proliferating astrocytes, as well as serum-exposed NPC *in vitro*, all upregulate the transcription regulator Id3, which is also expressed by astroglial limitans borders along meninges and large vessels in healthy tissue but is rare among other astrocytes not detectably present at these neural borders. Interestingly, others have shown that blood-derived fibrinogen can instruct endogenous periventricular NPC to upregulate Id3 and adopt a wound repair astroglial phenotype^64^. Together, these findings support a model whereby newly proliferated astroglia that form borders around CNS lesions represent a naturally occurring neuroprotective wound repair phenotype^65^ and that non-cell autonomous cues in serum-exposed CNS lesions activate similar intrinsic potentials in grafted NPC and proliferating host astroglia towards this naturally occurring wound repair astroglia phenotype.

NPC derived from ESC or iPSC and grafted into CNS lesions are well known to bias towards astroglial-like differentiation with formation of few neurons in the absence of exogenous interventions but why this occurs and the degree to which it is cell autonomously or non-cell autonomously determined have not been well understood^62^. Because NPC generate primarily astroglia instead of neurons when transplanted into CNS lesions, the lesion environment is often regarded as hostile. Our findings challenge the notion that lesions are ‘hostile’ to grafted NPC and instead provide an explanation that NPC are responding to cues in the lesion environments that instruct NPC and host proliferating astrocytes to adopt a naturally occurring wound repair astroglial phenotype. Understanding how and why different non-cell autonomous cues instruct NPC transplanted in CNS host tissue to generate different cell types will be fundamental to achieving mechanism-based approaches to therapeutic NPC transplantation. Monitoring transcriptional profiles of transplanted NPC *in vivo* over time as described here can facilitate the dissection of such mechanisms.

## Methods

### Cells

Mouse embryonic stem cells (mESC) were derived from the inner cell mass of E3.5 blastocyst stage embryos generated from crosses of male homozygous B6N.129-Rpl22tm1.1 Psam/J(RRID: IMSR_JAX: Stock No: 011029) “Ribotag” mice to females hemizygous for a dominant, maternal effect cre allele, B6.Cg-Tg(SOX2-cre)1Amc/J (RRID: IMSR_JAX: 008454) and heterozygous for the “RiboTag” allele. Multiple male and female mESC lines were derived, and each was karyotyped and genotyped to confirm sex and homozygosity for the cre-exised, Ribotag allele ^35^. A single female mESC line was used for all experiments in this work. These RiboTag mESC express a modified ribosomal protein L22 (Rpl22) with hemagglutinin HA epitope tag ^34^.

### Animals

All *in vivo* experiments involving the use of mice were conducted according to protocols approved by the Animal Research Committee (ARC) of the Office for Protection of Research Subjects at University of California Los Angeles (UCLA). All *in vivo* animal experiments were conducted within approved UCLA facilities using wildtype or transgenic C57/BL6 female and male mice that were aged between 8 weeks and four months old at the time of craniotomy or spinal cord injury surgery. B6N.129-Rpl22tm1.1Psam/J (RRID: IMSR_JAX: 011029) were bred with B6.Cg-Tg(Gfap-cre)73.12 Mvs/J (RRID: IMSR_JAX: 012886) from an in-house colony to generate Transgenic 73.12 GFAP Cre -RiboTag mice. B6.Cg-Tg(Gfap-TK)7.1Mvs/J (RRID: IMSR_JAX: 005698) were bred from an in-house colony to generate GFAP-TK mice. Mice were housed in a 12-hour light/dark cycle in a specific pathogen-free facility with controlled temperature and humidity and were provided with food and water ad libitum. Mice that received surgical procedures were administered post-surgical analgesia (buprenorphine, 0.1mg/kg) for at least 2 days after each surgery. Spinal cord injury mice received twice daily bladder expression until voluntary voiding returned. No animals in the study that received NPC transplants were administered with any immunosuppression drugs.

### NPC derivation from mESC

Neural progenitor cells (NPC) were derived from mouse embryonic stem cells (mESC) by adapting and refining existing protocols used by us and others^35,36,66^. mESCs were maintained and expanded on 0.1% gelatin-coated flasks (EmbryoMax™ 0.1% Gelatin Solution, Millipore Sigma, Cat# ES006B) in KnockOut™ DMEM media (Cat# 10829018, ThermoFisher Scientific) containing 15% Fetal bovine serum (FBS) (ES cell tested) (ThermoFisher Sci cat # 10439-016), MEM Non-Essential Amino Acids Solution (100X) (Cat# 11140050, ThermoFisher Scientific), 100X EmbryoMax® Nucelosides (EMD Millipore cat # ES-008-D), Beta-Mercaptoethanol (Sigma Aldrich M3148), Antibiotic-Antimycotic (100X) (Cat# 15240096, ThermoFisher Scientific) and Leukemia inhibitory factor (LIF) (1 million units/mL stock) (EMD Millipore cat # ESG1106). NPC lines were derived from mESCs using a standard 2-/4+ induction protocol followed by a neural expansion protocol ^35,36,66^. The 2-step of embryoid body (EB) formation involved removing LIF and FBS supplements from the cell media and changing media to a differentiation media consisting of Advanced DMEM/F12 (ThermoFisher Sci cat # 12634-010) supplemented with L-Glutamine (Thermofisher Scientific Cat#25-030-081) (2mM) and Knockout serum replacement (ThermoFisher Sci cat # 10828010) plus the other constituents described above for 2 days. Neural induction (4+) of embryoid bodies (EB) was performed by culturing isolated EB in differentiation media supplemented with retinoic acid (RA) (R2625-50MG, Sigma) (50nM) and purmorphamine (PUR) (sonic hedgehog (SHH) agonist) (SML0868-5MG, Sigma) (500nM) for 4 days. To propagate the neurally induced mESCs, the cells were grown in neural expansion media consisting of Advanced DMEM/F12 supplemented with B27 (no Vitamin A) (50X) (ThermoFisher Sci cat # 12587010) and growth factors EGF (Cat# AF-100-15-100UG, Peprotech) and FGF (Cat# 100-18B-100UG, Peprotech) (100ng/ml for each) for a period of 2-6 weeks resulting in a passagable line of adherent NPC. Samples at defined passages after neural induction (P) were collected for analysis. NPC stocks were maintained so that all samples in the study were less than P30.

### *In vitro* NPC assays

NPC were differentiated spontaneously (Spont) by lowering the EGF/FGF concentration in the neural expansion media from 100ng/ml to 1ng/ml for 4 days. CNTF (Cat# 450-13-100UG, Peprotech) and IGF-1 (Cat#100-11-100UG, Peprotech) differentiation involved adding 100ng/ml of the respective growth factor into the Spont media and culturing for 4 days. FBS differentiation involved adding FBS (Cat# 10437010, ThermoFisher Scientific) at 1, 5, 10 or 20% vol/vol to the Spont media and culturing for 4 days. For ICC evaluations cells were seeded onto 10mm round glass coverslips that had been coated with 0.1% gelatin and placed into wells of a 24 well plate at 30,000 cells per well and cultured with 1ml of media. At 4 days, cells on coverslips were fixed with 4% paraformaldehyde for 30 minutes. For RNA-Seq and Western blotting samples, cells were seeded at 30,000 cells per cm^2^ on T75 flasks (total of 2.25 million cells per flask) and recovered by gentle trypsinization and centrifugation before being frozen as cell pellets and stored in -80C cold storage until processing. TGF-βR inhibition of NPC cultured in 10% FBS was performed by preincubating NPC in media supplemented with the small molecule SB 431542 hydrate (S4317-5MG, Sigma) at a concentration 1μM for 2 hours prior to adding the FBS and then culturing for 4 days. Fractionation of FBS was performed using Amicon Ultra-0.5 Centrifugal Filter Unit (100 kDa MWCO, UFC510024, Sigma) spinning on a fixed angle rotor at 14,000g for 30 minutes to concentrate the >100kDa fraction. Fractionation was performed in 12 separate filter units and the >100kDa and <100kDa fractions from each were pooled. The concentration factor for the >100kDa sample was estimated from the pooled sample and diluted to equivalent concentration of the pre-fractionated FBS for use in the *in vitro* cell cultures. The <100kDa fraction was used in the *in vitro* experiments as collected without any further dilution. Mouse serum was generated from pooled blood collected fresh from 12-16 week C57/BL6 female and male mice via cardiac puncture using a 3mL syringe with a 18 gauge needle. Collected blood was transferred to a microcentrifuge tube and allowed to clot for 30 minutes at room temperature. Serum was isolated from the clotted blood by gentle centrifugation at 1000 RPM for 10 minutes in a refrigerated centrifuge (Model 5415R, Eppendorf).

### Synthesis and formulation of hydrogels

Physically crosslinked hydrogels were formulated from polypeptides that were synthesized using procedures developed previously by our group ^1,35^. Polypeptide synthesis was performed in a N2 filled glove box using anhydrous solvents. Amino acid N-carboxyanhydride (NCA) monomers L-methionine (M) NCA, L-leucine (L) NCA and L-alanine (A) NCA were prepared by phosgenation in a tetrahydrofuran (THF) solution under inert conditions. NCA monomers were purified by either recrystallization (for L and A NCA) or column chromatography in the glove box (M NCA). Copolypeptides were prepared at 100 mg scale, by adding a solution of initiator, Co(PMe3)4, (3.5 mg, 0.001 mmol) in THF (20 mg/mL) to a mixture of L-methionine NCA (Met NCA; 100 mg, 0.57 mmol) and L-alanine NCA (Ala NCA; 7.3 mg, 0.063 mmol) in THF (50 mg/mL). After 2 hours, L-leucine NCA (Leu NCA; 14 mg, 0.088 mmol) in THF (50 mg/mL) was then added and after a further 1 hour the reaction mixture was removed from the glove box. The block copolypeptide solution was precipitated in deionized water, filtered, and dried under reduced pressure. Next, a volume of 70 wt. % *tert*-butyl hydroperoxide (TBHP) (16 molar equivalents per L-methionine residue) was added to the copolypeptide in DI water to convert L-methionine residues to L-methionine sulfoxide residues over 24 hours at ambient temperature. To aid oxidation, a catalytic amount of camphorsulfonic acid (CSA) (0.2 molar equivalents per Met residue) was added to the solution. Polypeptide was dialyzed in 2000 MWCO dialysis bags against: (i) pyrogen free deionized milli-Q water (3.5 L) containing sodium thiosulfate (1.2 g, 2.2 mM) for 1 day to neutralize residual peroxide, (ii) pyrogen free deionized milli-Q water (3.5 L) containing ethylenediaminetetraacetic acid tetrasodium salt hydrate (1.0 g, 2.63 mmol) to aid cobalt ion removal, and (iii) pyrogen free deionized milli-Q water (3.5 L) for 2 days to remove residual ions. For each step dialysate was changed every 12 hours. The copolypeptide solution was then freeze dried to yield a white fluffy solid. Hydrogels were prepared by solubilizing lyophilized copolypeptide in phosphate buffered saline (PBS) or supplemented cell culture media and stored at 4°C for 24 hours to assemble, without stirring, before use. DCH formulations mixed with cells were prepared at either 10 or 5 wt% and then diluted 1:1 with cells in media via gentle mixing with a pipette. DCH alone formulations were prepared at 5% or 2.5% in PBS.

### Surgical procedures

All surgical procedures were approved by the UCLA ARC and conducted within a designated surgical facility at UCLA. All procedures were performed on mice under general anesthesia achieved through inhalation of isoflurane in oxygen-enriched air.

#### Craniotomy procedure

Shaved mice heads were stabilized and horizontally leveled in a rodent stereotaxic apparatus using ear bars (David Kopf, Tujunga, CA). A small craniotomy over the left coronal suture was performed using a high-speed surgical drill and visually aided by an operating microscope (Zeiss, Oberkochen, Germany). As small rectangular flap of bone encompassing sections of the frontal and parietal bone was removed to expose the brain in preparation for injection.

#### Stroke lesion model

To create focal ischemic strokes, 1.5 μL of L-NIO (N5-(1-Iminoethyl)-L-ornithine) (Cat. No. 0546, Tocris solution) (27.4 mg/ml (130 μM) in sterile PBS) was injected into the caudate putamen nucleus at 0.15 μL/min using target coordinates relative to Bregma: +0.5 mm A/P, +2.5 mm L/M and −3.0 mm D/V. A standard micropipette injection protocol was used to make all injections into the brain or spinal cord using pulled borosilicate glass micropipettes (WPI, Sarasota, FL, #1B100-4) that were ground to a 35° beveled tip with 150–250 μm inner diameter. Glass micropipettes were mounted to the stereotaxic frame via specialized connectors and attached, via high-pressure polyetheretherketone (PEEK) tubing, to a 10 μL syringe (Hamilton, Reno, NV, #801 RN) controlled by an automated syringe pump (Pump 11 Elite, Harvard Apparatus, Holliston, MA). After injection the surgical incision site was sutured closed and the animals were allowed to recover for 48 hours before undergoing a second surgery involving injection of hydrogel with and without cells as described below.

#### Spinal Cord Injury crush model

A single incision was made along the back of mice and back musculature attached to the vertebrae cut and removed using small spring scissors and sterile cotton swabs. A laminectomy was made at the 10th thoracic (T10) spinal vertebrae level using spring scissors and 2SP laminectomy forceps (FST) to expose the dorsal surface of the spinal cord. A timed (5 second) lateral compression crush injury at the laminectomy site was made using No. 5 Dumont forceps beveled to a tip width of 0.5 mm.

#### NPC and hydrogel injections

Cells and hydrogel formulations were backloaded into the pulled glass micropipettes prior to connecting to the stereotaxic frame. For brain injections, a volume of 1 μL of hydrogel and/or cells (loaded at 10,000, 100,000 or 250,000 cells/μL) were injected into the caudate putamen nucleus at 0.15 μL/min using target coordinates relative to Bregma: +0.5 mm A/P, +2.5 mm L/M. The pipette was lowered to −3.5 mm D/V for the start of the injection and moved up +0.5mm twice over the course of the injection to a final location of −2.5 mm D/V to improve deposition of cells in brain tissue. The micropipette was allowed to dwell in the brain at the injection site for an additional 4 minutes at the end of the injection. The micropipette was then removed from the brain slowly and incrementally over a 2-minute period. The same procedures were used for injections into L-NIO stroke lesions at 2 days after stroke induction. For hydrogel and/or cell injections into SCI lesions, at two days after SCI, NPC formulations were transplanted at a concentration of 250,000 cells/μL with four individual injections of 0.3 μL (4x 0.3μL = 1.2μL total) made at the lesion site at a depth of 0.6mm from the dorsal surface of the spinal cord.

### Transcardial perfusions

After terminal anesthesia by intraperitoneal injection of pentobarbitol, mice were perfused transcardially with heparinized saline (10 units/ml of heparin) and 4% paraformaldehyde (PFA) that was prepared from 32% PFA Aqueous Solution (Cat# 15714, EMS), using a peristaltic pump at a rate of 7 mL/min. Approximately, 10 mL of heparinized saline and 50 mL of 4% PFA was used per animal. Brains and spinal cords were immediately dissected after perfusion and post-fixed in 4% PFA for 6-8 hours. After PFA post-fixing, brains and spinal cords were cryoprotected in 30% sucrose in Tris-Buffered Saline (TBS) for at least 3 days with the sucrose solution replaced once after 2 days and stored at 4°C until further use. For RiboTag RNA-Seq experiments mice were perfused transcardially with cold heparinized saline (10 units/ml of heparin) for 2 minutes (7mL/min) and a 4mm segment of spinal cord centered on the lesion was immediately dissected after perfusion on ice and snap frozen in microcentrifuge tubes on dry ice. Frozen spinal cords were kept at -80°C until RiboTag Immunoprecipitation (IP) processing.

### Immunohistochemistry and immunocytochemistry

Coronal brain sections (40 μm thick) or horizontal spinal cord sections (30 μm thick) were cut using a Leica CM3050 cryostat. Tissue sections were stored transiently in TBS buffer at 4°C or in antifreeze solution of 50% glycerol/30% Sucrose in PBS at -20°C for long term storage. Tissue sections were processed for immunofluorescence using free floating staining protocols described in comprehensive detail previously using donkey serum to block and triton X-100 to permeabilize tissue ^1,56,60^. The primary antibodies used in this study were: rabbit Hemagglutinin (HA) (1:1000, Sigma #H6908); goat HA (1:800, Novus, NB600-362); rabbit alpha-smooth muscle actin (α-Sma) (1:200, Novus, NB600-531); rabbit anti-Gfap (1:1000, Dako/Agilent, GA524); rat anti-Gfap (1:1000, Thermofisher, #13-0300); rabbit anti NeuN (1:1000, Abcam, ab177487); guinea pig anti-NeuN (1:1000, Synaptic Systems, 266 004); goat anti-Cd13 (1:200, R&D systems, AF2335); rat anti-Galectin-3 (1:200, Invitrogen, 14-5301-82); rabbit anti-Fibronectin (1:500, Millipore, Cat#AB2033); rat anti-Cd68 (1:1000, AbDserotec-BioRad, MCA1957); rabbit anti-Iba-1 (1:800, Wako, 019-19741); guinea pig anti-Iba-1 (1:800, Synaptic systems, 234 004); rabbit anti-P2ry12 (1:500, Anaspec, AS-55043A); goat anti-Pdgfr-α (1:500, R&D systems, AF1062); goat anti-Nestin (1:500, R&D, AF2736); goat anti-Oct4 (1:500, R&D systems, AF1759); goat anti-Sox9 (1:500, R&D systems, AF3075); rabbit Aldh1l1 (1:1000, Abcam, Ab87117); rabbit anti-Amyloid Beta (Aβ) (1:200, Abcam, Ab2539); rabbit anti-Amyloid precursor protein (App) (1:200, abcam, ab32136); goat anti-Carboxypeptidase E/CPE (Cpe) (1:200, R&D systems, AF3587); goat anti-Lipocalin-2 (Lcn2) (1:200, R&D systems, AF1857), goat anti-Clusterin (Clu) (1:200, R&D systems, AF2747); Rabbit anti-Tuj-1 (1:500, Sigma, T2200-200UL); rat anti-Vimentin (1:500; R&D Systems, MAB2105); rat anti-Cd44 (IM7) (1:200; Thermofisher Scientific, #14-0441-82); goat anti-Dppa4 (1:200; R&D Systems, AF3730), Rabbit anti-Id3 (1:200; Cell Signaling Technology, #9837); rabbit anti–HSV-TK (1:1000) (generated by M. Sofroniew and validated previously^67^). All secondary antibodies used in this study were purchased from Jackson ImmunoResearch (West Grove, PA). All secondary antibodies were affinity purified whole IgG(H+L) antibodies with donkey host and target species dictated by the specific primary antibody used. All secondary antibodies were stored in 50% glycerol solution and diluted at a concentration of 1:250 in 5% normal donkey serum in TBS when incubated with brain histological sections. Nuclei were stained with 4’,6’-diamidino-2-phenylindole dihydrochloride (DAPI; 2ng/ml; Molecular Probes). Acti-stain 555 phalloidin (Cytoskeleton Inc. Cat. # PHDH1-A) staining was performed in conjunction with DAPI nuclei staining using manufacturer’s instructions. Sections were coverslipped using ProLong Gold anti-fade reagent (InVitrogen, Grand Island, NY). Sections were examined and photographed using epifluorescence microscopy, deconvolution widefield fluorescence microscopy and scanning confocal laser microscopy (Zeiss, Oberkochen, Germany). Tiled scans of individual whole sections were prepared using a x20 objective and the scanning function of a Leica Aperio Versa 200 Microscope (Leica, Wetzlar, Germany) available in the UCLA Translational Pathology Core Laboratory.

### RiboTag Immunoprecipitation, RNA purification and RNA-Sequencing

Frozen 4mm segments of SCI tissue containing HA-positive NPC transplants or pellets of cells collected from *in vitro* experiments were processed by RiboTag Immunoprecipitation (IP) using established methods^34,60^. SCI tissue or cell pellet samples were homogenized in supplemented homogenization buffer (50 mM Tris pH 7.4, 100 mM KCl, 12 mM MgCl2, 1% NP-40, 1 mM Dithiothreitol (DTT), 1X Protease inhibitors, 200 U/ml RNAsin, 100 mg/ml Cyclohexamide, 1 mg/ml Heparin) using 2mL glass Dounce tissue grinders (D8938-1SET, Sigma). Tissue homogenates were centrifuged to remove tissue debris before being incubated with Anti-HA.11 Epitope Tag Antibody (Biolegend, Cat# 901515) for 4 hours in a microcentrifuge tube on a microtube rotator kept at 4°C. At the end of 4 hours, IP solutions were combined with Pierce A/G Magnetic Beads (Thermofisher, #PI88803) and incubated overnight on a microtube rotator at 4°C. On the second day, magnetic beads were washed three times with high salt solution (50 mM Tris pH 7.4, 300 mM KCl, 12 mM MgCl2, 1% NP-40, 1 mM Dithiothreitol (DTT), 100 mg/ml Cyclohexamide). Unpurified RNA was collected from the magnetic beads by addition of RLT Plus buffer with BME and vigorous vortexing. RNA was then purified using RNeasy Plus Mini (for *in vitro* cell pellets) or Micro Kits (for spinal cord tissue) (QIAGEN Cat# 74134 and 74034). Total mRNA derived from the RiboTag IP was quantified using an automated electrophoresis bioanalyzer. RNA samples with RNA integrity numbers (RIN) greater than seven were processed for RNA-Sequencing. Sequencing was performed on poly-A selected libraries using Illumina NovaSeq S2 (housed in the UCLA Technology Center for Genomics & Bioinformatics) using pair end reads (2×50 – 50bp length) with an average of 50-100M reads per sample. We performed sequence alignment and transcript counting on raw FASTQ files using standardized Galaxy workflows described in detail below.

### Nuclei preparation from *in vitro* cell cultures and single nuclei RNA-Sequencing

Cultures of NPC and NPC differentiated using the Spont or CNTF conditions (∼2.25 × 10^6^ cells per condition) were trypsinized and pelleted by centrifugation. Nuclei were extracted from cell pellets by gentle resuspension in ice-cold lysis buffer (10mM Tris buffer, 10mM NaCl, 3 mM MgCl2, 0.1% Nonidet P40 Substitute). Nuclei were pelleted by centrifugation (500RPM (Model 5415R, Eppendorf) for 5 min at 4°C) and then resuspended in Nuclei Wash and Resuspension Buffer (NWRB) (1xPBS, 1% BSA, 0.2U/uL RNAse inhibitor). Nuclei were assessed by trypan blue staining before being washed once more in NWRB and then filtered using a 5 mL Polystyrene Round-Bottom Tube with 35 μm Cell-Strainer Cap and concentrated to a nuclei concentration of 1000 nuclei/μL (1 × 10^6^ nuclei/mL). Dissociated single nuclei from the three conditions were processed using Chromium Next GEM Single Cell 3’ v3.1 kits using manufacturer’s instructions and libraries processed for RNA-Sequencing using Illumina NovaSeq S2 (housed in the UCLA Technology Center for Genomics & Bioinformatics) using pair end reads (2×50 – 50bp length). Raw data files were processed using Scanpy workflows conducted using Galaxy and Galaxy Single Cell Omics platforms described in detail in the quantification section below.

### Western Blotting

Protein was extracted from frozen cell pellets by homogenization in RIPA lysis buffer supplemented with PMSF, protease inhibitor and sodium orthovanadate using a handheld grinder and attached pestle. Homogenized samples were kept on ice for 30 minutes after homogenization and then centrifuged to remove insoluble debris. Supernatants containing protein were decanted into fresh microcentrifuge tubes and stored at -80°C until use. Total protein in solutions was quantified prior to gel loading using standard BCA assay (Pierce™, Cat#23225, Thermofisher) following manufacturer’s instructions. SDS-PAGE was performed using NuPAGE Novex Bis-Tris Pre-Cast Gels in XCell SureLock Mini-Cell Electrophoresis System. Protein samples were prepared in NuPAGE LDS Sample Buffer and heated at 70C for 10 minutes prior to gel loading. For all runs, 15-30 μg protein per sample was used. Protein gel electrophoresis was run using 1X NuPAGE SDS Running Buffer and conducted at 150 volts for one hour. Separated proteins in the gel were transferred to PVDF membranes using 1X NuPAGE Transfer Buffer and XCell Blot Module by applying 30 volts for 1 hour following manufacturer’s instructions. Total protein across sample lanes transferred to the PVDF membrane was quantified using SYPRO Ruby. For western blotting, membranes were activated with methanol and blocked using non-fat milk in Tris Buffer Saline with Tween20 (TBST) for one hour followed by primary antibody incubation overnight at 4°C. Primary antibodies were prepared in 5%BSA/TBST. Primary antibodies used for WB included: rabbit anti-Fabp7 (1:2000, PA5-31864 Thermofisher); goat anti-Nanog (1:1000, AF2729-SP R&D Systems); rabbit anti-Oct4 (1:1000, MA5-14845, Invitrogen); goat anti-Sox9 (1:1000, R&D systems, AF3075); goat anti-Dppa4 (1:2000, Novus, AF3730); goat anti-Nestin (1:2000, R&D Systems, AF2736); rabbit alpha-smooth muscle actin (α-Sma) (1:2000), Novus, NB600-531; mouse anti-Gfap (1:3000, Sigma, Cat#3893). After primary incubation membranes were washed 3 times with TBST on a shaker before incubation with species appropriate HRP-conjugated secondary antibody in 5% non-fat milk/TBST for 1 hour. Secondary antibodies used were goat anti-rabbit (1:10000, A27036, ThermoFisher); donkey anti-goat (1:5000, A15999, ThermoFisher); goat anti-mouse (1:20000, Cat#62-6520, ThermoFisher). After additional washing bands were detected using SuperSignal West Pico PLUS Chemiluminescent Substrate and imaged on a ChemiDoc XRS+ Imaging System (BioRad).

### Statistics, power calculations, group sizes and reproducibility

Graph generation and statistical evaluations of repeated measures were conducted by one-way or two-way ANOVA with post hoc independent pair wise analysis as per Tukey, or by Student’s t-tests where appropriate using Prism 9 (GraphPad Software Inc, San Diego, CA). Statistical details of experiments can be found in the figure legends including the statistical tests used and the number of replicative samples. Across all statistical tests significance was defined as p-value <0.05. Power calculations were performed using G*Power Software V 3.1.9.2. For immunohistochemical quantification analysis and RNA-Sequencing, group sizes were calculated to provide at least 80% power when using the following parameters: probability of type I error (alpha) = .05, a conservative effect size of 0.25, 2-5 treatment groups with multiple measurements obtained per replicate. All graphs show mean values plus or minus standard error of the means (S.E.M.) as well as individual values as dot plots. All bar graphs are overlaid with dot plots where each dot represents the value for one animal to show the distribution of data and the number (N) of animals per group. Injections of NPC and hydrogel formulations were repeated independently at least three times in different colonies of mice across a two-year period with similar results.

### Principal Component Analysis (PCA) and Cosine similarity (CS)

Principal Component Analysis (PCA), Euclidean distance and Cosine Similarity (CS) analysis was performed using XLStat (Addinsoft Inc, Long Island City, NY). For PCA, 2 and 3 dimensions were used. PCA data was represented in both Euclidean distance and correlation plots as appropriate. Euclidean distance calculations were derived by assessing the vector magnitude in PCA space of a specific sample referenced to another sample as an initial point. Cosine similarity (CS) measures similarity of samples by comparing the angle between the sample vectors of n-genes and is reported on a scale of 0-1, with 1 being the most similar and 0 being orthogonal.

### Quantification of Immunohistochemistry

Immunohistochemical staining intensity quantification was performed on whole brain and spinal cord images derived from the slide scanner or tiled images prepared on the epifluorescence microscope. All images used for each comparable analysis were taken at a standardized exposure time and used the raw/uncorrected intensity setting. Quantification of antibody staining intensity was performed using NIH Image J (1.51) software using the Plot profile (for spinal cord sections) or radial profile angle (for brain sections) plugins using methods developed previously^1^. Total values for IHC stainings were determined by taking the integral (area under the curve) of the plot profile or radial intensity profile. Quantification of co-staining for HA and cell specific markers Gfap and Pdgfr-α was performed using the RG2B colocalization plugin. Orthogonal (3D) images were prepared using Imaris 9.2 (Bitplane) or Zen 3.1 (Blue Edition) (Zeiss).

### Generation of Gene Panels

To generate a list of genes enriched in healthy astrocytes we mined eight published healthy astrocyte specific and enriched gene expression datasets from mouse brain and spinal cord. Astrocyte enriched genes (AEG) were defined as having a Log2FC >2 compared to all other cells in the sample population or were defined as being detected specifically in astrocytes if derived from single cell data. Genes designated to the healthy astrocyte gene panel appeared in at least 5 of the 8 lists resulting in a total of 429 genes. The Oligodendrocyte lineage gene (n= 112 genes) and neuron gene (n=221 genes) lists were derived from a publicly available list generated from the aggregation of published single cell datasets (PanglaoDB ^68^). We derived the astrocyte reactivity gene panel by analyzing six unique data sets of astrocyte responses to injury in the acute setting available in the literature. The analysis included 3 spinal cord data sets and 3 brain data sets that all derived astrocyte specific transcriptomic information at acute injury timepoints of less than 7 days after injury. Using the principal criterion that genes needed to be included in at least 3 of the 6 lists, we generated a curated list of 170 genes deemed to be associated with an acute astrocyte response to injury. we also generated an EMT gene panel using the EMT Gene Set Enrichment Analysis (GSEA) dataset from the Molecular Signatures Database (MSigdb), which is a publicly available list derived from rodent and human studies. For the EMT gene panel there were 197 genes whose expression level could be assessed in mouse. Comprehensive gene panel information is provided in the Source data files.

### Transcriptomics analysis of RiboTag RNA-Sequencing data

Analysis of RNA-Seq raw data was performed in Galaxy using standardized workflows. The R1 and R2 FASTQ files from lanes 1 and 2 that were obtained directly from the Illumina NovaSeq S2 run were concatenated and cleaned-up using the Trimmomatic tool. Data was then aligned to the M. musculus (mm10) reference genome using the HISAT2 tool applying default parameters. Gene counts from the aligned datasets was performed using the featureCounts tool applying default parameters. Fragments Per Kilobase of transcript per Million mapped reads (FPKM) values were calculated for each gene directly in Excel (Microsoft) using standardized lists of gene lengths and normalization of the count data. Differential expressed gene (DEG) analysis on raw gene count data was conducted using Edge-R in Galaxy applying Benjamini and Hochberg p-value adjustment and TMM normalization. Across all studies we used a conservative false discovery rate (FDR) cut off < 0.01 to define significance of DEGs and evaluated at least 4 unique samples per experimental group. Gene ontology analyses were performed using Enrichr tool (https://maayanlab.cloud/Enrichr/). Differences in transcript expression across samples were evaluated using data-dimensionality reduction techniques including Principal Component Analysis, Euclidian distance and Cosine Similarity Analysis as described above. Heatmaps of DEG data were generated using NG-CHM BUILDER (https://build.ngchm.net/NGCHM-web-builder/). Violin plots of DEGs were generated using Prism 9.

### Analysis of single Nuclei RNA-Seq data

Raw single nuclei sequencing data was processed using RNA StarSolo on the Galaxy Single Cell Omics platform using M. musculus (mm10) reference genome, 3M-february-2018 barcode whitelist, the gencode vM25 annotation list, and Cell Ranger v3 configure chemistry options. Scanpy tools were used through the Galaxy platform to convert genes, barcodes and matrix files derived from RNA StarSolo into an AnnData matrix h5ad format. Individual h5ad files for NPC, CNTF and SPONT samples were generated separately and concatenated (merged) into a single data set in h5ad format using the Manipulate AnnData tool and applying unique batch designations. The combined data set was preprocessed to remove mitochondrial genes, filter out genes that were detected in less than 200 cells and filter out low quality cells that had less than 333 attributed genes. Using Scanpy tools we normalized the dataset, identified the 5000 most variable genes across the total population and generated the UMAP projection using the 5000 gene list. Cell clustering was performed on the UMAP data via Scanpy using Louvain clustering algorithms with a resolution of 0.5 which resulted in 11 discrete clusters. Marker genes that defined the clusters were determined using the Scanpy FindMarkers tool applying default parameters. Clusters were assigned NPC, astrocyte or oligodendroglia lineage designation on the basis of expression of known marker genes for those specific cells. Violin plots of gene expression distribution within clusters was also performed using Scanpy. The percentage of cells from each condition detected within each cluster was determined from the batch designations assigned upon merging of the datasets.

### Data availability statement

RiboTag RNA-Seq and Single-cell RNA-seq data have been deposited at Gene Expression Omnibus (GEO) and are publicly available as of the date of publication with Accession numbers XXXX. All data generated for this study are included in the main and supplementary figures. For all quantitative figures, files of source data of individual values as well as the results of statistical tests are provided with the paper. Other data that support the findings of this study are available on reasonable request from the corresponding author.

## Acknowledgments

Thanks to the UCLA Neuroscience Genomics Core for assistance with sequencing. This work was supported by US National Institutes of Health (NS084030 to M.V.S.); Dr. Miriam and Sheldon G. Adelson Medical Foundation (M.V.S. and T.J.D); Craig H. Neilsen Foundation (381357 to T.M.O.); Paralyzed Veterans Foundation of America (RF3170 to T.M.O.); American Australian Association (T.M.O.), Wings for Life Spinal Cord Research Foundation (M.V.S. and T.M.O.); US National Institutes of Health (OD010921 to L.G.R.); and Microscopy Core Resource of UCLA Broad Stem Cell Research Center.

## Author contributions

T.M.O carried out biomaterial preparations, *in vitro* cell culture, surgical procedures, RiboTag IP, tissue processing for IHC and RNA-Seq, microscope imaging, IHC image analysis, and processed and analyzed bulk and single-nuclei RNA-seq data. Y.A. carried out tissue processing, IHC staining and microscope imaging. S.W. carried out Western blotting, ICC, RiboTag IP and Single Nuclei isolations. A.L.W synthesized hydrogel biomaterial, J.H.K performed cell culture, RiboTag IP and IHC image analysis. R.R.E prepared single nuclei analysis plots, A.C and L.G.R generated the RiboTag mESC line, T.J.D. derived the plan for, and supervised, the biomaterial synthesis. M.V.S. and T.M.O. conceived, designed, and directed the project and guided data analyses. T.M.O analyzed data and assembled the figures with feedback from M.V.S. T.M.O and M.V.S wrote the paper and all authors commented and edited it.

## Statement about Competing interests

The authors declare no competing interests.

