## Supplemental Information for "Lesion environments direct transplanted neural progenitors towards a wound repair astroglial phenotype"

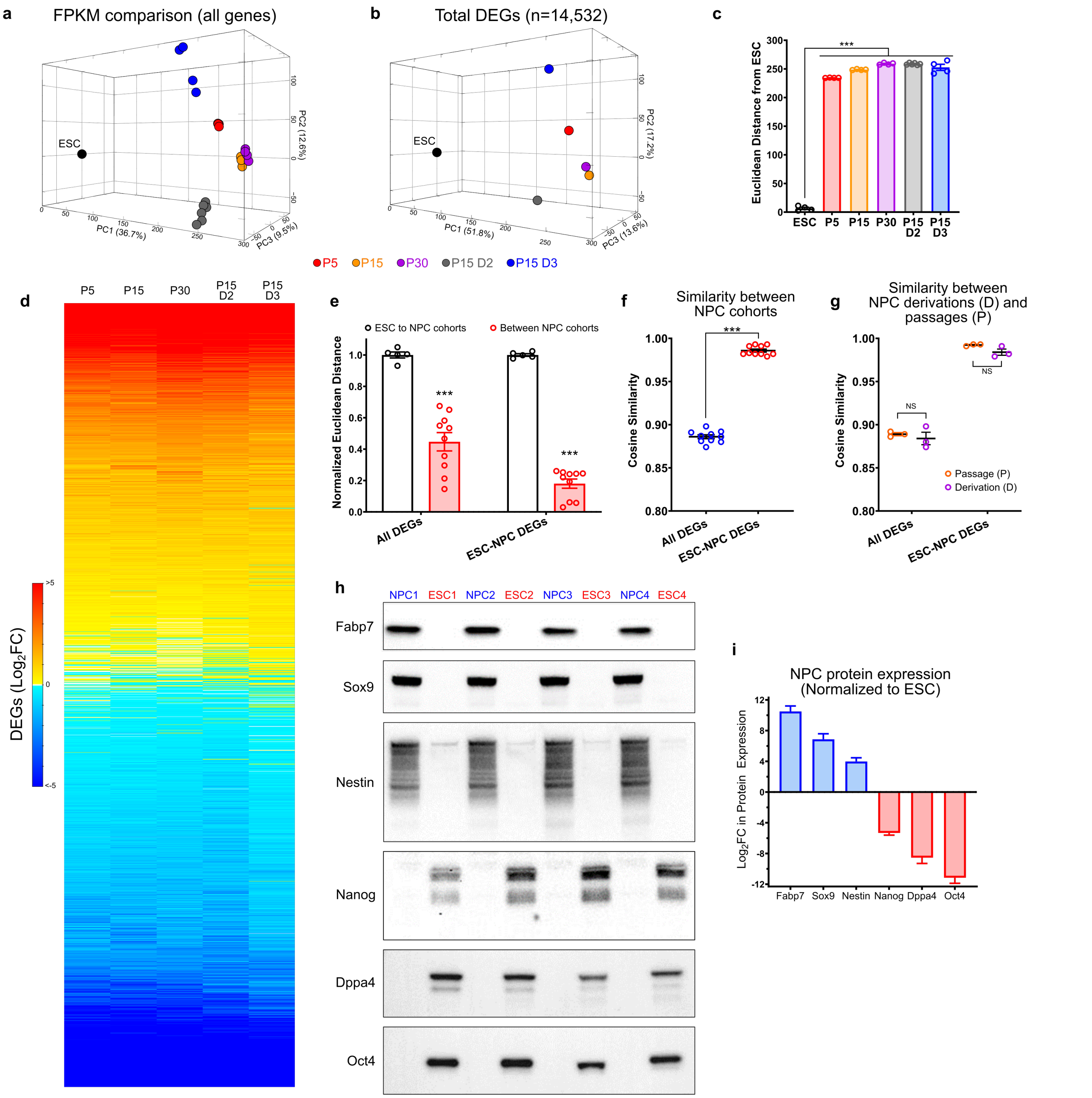


**Supplementary Fig. 1 | Neural induction and expansion of RiboTag mESC derives a reproducible and stable NPC line. a** PCA of NPC samples from different derivations (d) and passages (P) comparing all expressed genes (genes measured with an FPKM >0.1 in at least one sample) referenced to ESC sample average. n=4 per group for all groups. **b** PCA of NPC samples from different derivations (d) and passages (P) comparing all DEGs (DEG defined as having an FDR≤0.01) for NPC samples referenced to ESC. n=4 per group for all groups. Total number of DEGs used in PCA = 14,352. **c** Euclidean distance comparisons for ESC and NPC samples. Euclidean distance equates to the vector magnitude in PCA space of NPC samples referenced to mean of ESC samples as the initial point, using FPKM values for all detected genes. n=4, *** p-value < 0.0001, One way ANOVA with Tukey multiple comparison test. **D** Heatmap of all DEGs referenced to ESC for NPC at different Passages (P) and derivation (D). **e** Normalized Euclidean distance evaluations comparing the scaled vector lengths between ESC and NPC samples with the scaled vector lengths between individual NPC samples from different Passages (P) and derivation (D) for all DEGs (n=14,532 genes) or for ESC-NPC gene panel DEGs (n=88 genes). N=5 and 10, *** p-value < 0.0001, Two way ANOVA with Tukey multiple comparison test. **f** Cosine similarity comparisons for NPC samples from different Passages (P) and derivation (D) using all DEGs (n=14,532 genes) or ESC-NPC gene panel DEGs (n=88 genes). N=10 *** p-value < 0.0001, two-tailed, unpaired Student's t-test. **g** Cosine similarity comparisons for NPC samples comparing by different Passage number (P) or derivation (d). N=3 per group, Not significant (NS), Two way ANOVA with Tukey multiple comparison test. **h** Western blot of NPC and ESC samples (N=3 samples per cell type) for canonical NPC markers Fabp7, Sox9, Nestin as well as canonical ESC makers Nanog, Dppa4 and Oct4. **i** Quantification of Western blots measuring protein expression of canonical NPC and ESC markers. Expression levels in all samples were referenced to total protein that was quantified by SYPRO Ruby Protein Blot Stain. Mean NPC expression levels were referenced to mean ESC levels (Mean + SEM).

### Astrocyte Gene Panel

| | | | | | |
|---|---|---|---|---|---|
| Aldoc | Papss2 | Hist1h2ac | Aass | Fam181a | Tmem176b |
| Apoe | Paqr6 | Fzd9 | Lfng | Atp13a4 | Trp53bp2 |
| Aldh1l1 | Hyal1 | Cyr61 | Gli2 | Rfx4 | Acsm5 |
| Htra1 | Vcam1 | Cpq | Fam181b | BC064078 | Ctso |
| Slc1a3 | Pld2 | Aldh2 | Dmp1 | Car5b | Vim |
| Sox9 | Ppp1r3c | Acaa2 | Mmp14 | Etv4 | S100b |
| Ntsr2 | Tmem51 | Myh15 | Ppp1r3g | P4ha3 | Fam107a |
| Gja1 | Scg3 | Exoc3l4 | Hsd11b1 | Dag1 | Apln |
| S1pr1 | Elf5 | Tmco4 | Kctd14 | Slc4a4 | Mapkapk3 |
| Fgfr3 | Phka1 | Bdh2 | Ezr | Dio2 | Stk17b |
| Pla2g7 | Cideb | Rbm46 | Fgfrl1 | Fabp7 | Lrp10 |
| Slc1a2 | Emp2 | Etnppl | Slc2a10 | Igsf1 | Pnpla7 |
| Lcat | Ednrb | Pdlim5 | Pbxip1 | Fkbp10 | Tmem100 |
| Cldn10 | Npas3 | Irak2 | Rapgef3 | Slc12a4 | Acadm |
| Acsl6 | Pon2 | Smox | Steap3 | Mfge8 | Itga6 |
| Entpd2 | Slc15a2 | Renbp | Fxyd1 | Gdf10 | Ech1 |
| Agt | Sash1 | Aldh4a1 | Cyp2j9 | Lgr6 | Gpc6 |
| Gpr37l1 | Adhfe1 | Mtss1l | Nwd1 | Serhl | Elovl5 |
| Slc6a11 | Gm2a | 2310022B05Rik | Id3 | Itga7 | Dapp1 |
| Acsbg1 | Bmpr1b | Sugct | Sdc4 | Oplah | Gm17455 |
| Fads2 | Scd2 | Myo10 | Acss1 | Ip6k3 | Kank1 |
| Plxnb1 | Cd81 | Phyh | Cth | Slc13a3 | Apoc1 |
| Clu | Nat1 | Plin2 | Sardh | Pygm | Hist1h4j |
| Slc39a12 | Smpdl3a | Pdlim4 | Kirrel2 | Ltbr | Cables1 |
| Cbs | Oat | Prss50 | Pm20d1 | Cyp2d22 | Acad11 |
| Slc27a1 | Rnase1 | Id1 | Ltbp1 | Sdsl | Lonrf3 |
| Gfap | Pxmp2 | Erbb2 | Msi1 | Gstt1 | Decr1 |
| Atp1a2 | Tnfrsf19 | Loxl3 | Fam69c | Heph | Acacb |
| Srebf1 | Fjx1 | Itpr2 | Alpl | Selenbp1 | Baalc |
| Pygb | Als2cl | Cnn3 | Axl | Fzd2 | Fgd6 |
| Slc7a10 | Eno1b | Aldh7a1 | Hspa2 | Tdgf1 | Smo |
| Itih3 | Gm973 | Hrh1 | Mt2 | Cdc42ep4 | Scd1 |
| Gstm1 | Ptprz1 | Notch2 | Sox2 | Myoc | As3mt |
| Sparcl1 | Tlr3 | Abhd4 | Slc13a5 | Plcd1 | Lrrc8a |
| Ndrg2 | Myo6 | Tmem176a | Phkg1 | Plcb3 | Inppl1 |
| Cpe | Luzp2 | Aldh1a1 | Slc25a18 | Klhdc7a | Cybrd1 |
| Rgma | Timp4 | Bcar3 | Pard3b | Asrgl1 | Stat5a |
| Tril | Pdpn | Thnsl2 | 2810459M11Rik | Hes5 | Tead1 |
| Tst | Myom3 | Necap2 | Tnc | Fzd10 | Ston2 |
| Nkain4 | Mgst1 | Acad12 | Fam20a | Tnfrsf1a | Add3 |
| Prodh | Serpine2 | Rftn2 | Rlbp1 | Arhgef19 | Gpr179 |
| Dbx2 | Acsl3 | Hgf | F3 | Mfsd2a | Yap1 |
| Sfxn5 | Ntrk2 | Epx | Pdk4 | Cntfr | Tspan7 |
| Mmd2 | Mt1 | Btd | Ephx2 | Gstt3 | 4921530L21Rik |
| Abhd3 | Rarres1 | Arhgef26 | Lgi4 | Smpd2 | Rnf182 |
| Timp3 | Tnfaip8 | Oaf | Agl | Gpam | Ptch1 |
| Atp1b2 | Nrarp | Eci1 | Psd2 | Gpc5 | Kcnj16 |
| Hepacam | Gabrg1 | Acads | Grin2c | Idh2 | Hbegf |
| Gjb6 | Rgcc | Egfr | Sema4b | Slc8b1 | Zfp36l2 |
| Slc1a4 | Soat1 | Lrrc2 | Paqr8 | Lgals3bp | A730056A06Rik |
| Aqp4 | Adk | Adcyap1r1 | Slc25a34 | S100a1 | Laptm4a |
| Lxn | Lix1 | Tns3 | Palld | Chil1 | Gm266 |
| Glud1 | Gdpd2 | Slc2a12 | Maob | Plekho2 | Bbox1 |
| Ncan | Ckb | Grm3 | Cst3 | Dspp | Ptn |
| Ttyh1 | Ndp | Lpar4 | Tlcd1 | Sfrp5 | A330048O09Rik |
| Lpin3 | Smyd1 | Fam213a | Cd63 | Btbd17 | Rgl3 |
| Slc9a3r1 | Rorb | Acadl | Egfl6 | Gstk1 | Mr1 |
| Slc38a3 | Hadhb | Abca1 | Prdx6 | Suclg2 | Mdk |
| Kcne1l | 4932438H23Rik | Hacl1 | Ppara | Eps8 | Frem2 |
| Cyp4f14 | Mamdc2 | 3110082J24Rik | Amot | Fads1 | Gli3 |
| A2m | Sox21 | Gprc5b | Mertk | Padi2 | Rgs20 |
| Lrp4 | Calr4 | 5930403L14Rik | Slc14a1 | Olfr287 | Id4 |
| AI464131 | Nek8 | Eva1a | Lrig1 | Tekt4 | Aqp9 |
| Mlc1 | Msx2 | Evc | Cd38 | Sparc | Ttpa |
| Rdh5 | Rbpms2 | Fmo5 | Kcnj10 | Snta1 | Pax6 |
| Bcan | Echdc3 | Elovl2 | Ucp3 | Cpt1a | Cbr3 |
| Acsf2 | D630039A03Rik | Dock1 | Mcc | Sh3pxd2b | Glycam1 |
| C4b | Slc6a1 | A730036I17Rik | Ddo | Igdcc4 | Nqo1 |
| Gldc | Gm11627 | Thrsp | Adora2b | Cyp4v3 | Prkd1 |
| Acot11 | Micalcl | Ddah1 | Slc7a2 | Glipr2 | Cxcl14 |
| Slc2a4 | Megf10 | Mid1ip1 | Cyp4f15 | Ccdc8 | Nat8 |
| Pttg1ip | Mt3 | Arhgap5 | | | |

### Oligodendroglia

| | |
|---|---|
| Gpr17 | Tmem88B |
| Neu4 | Bace1 |
| Fa2H | Vcan |
| Ninj2 | Olig3 |
| Tmem163 | Gpr62 |
| C1Ql1 | Slc25A38 |
| Opalin | S1Pr5 |
| Enpp6 | Aspa |
| Gpr37L1 | Zdhhc9 |
| Gjc3 | Tmem125 |
| Sox10 | Hapln2 |
| Plp1 | Gjb1 |
| Tmem100 | Ermn |
| Dusp15 | Klk6 |
| Omg | Cldn14 |
| Pcdh15 | Sp7 |
| Mbp | Prkcz |
| Nkx6-2 | Klhl2 |
| Tnr | Olig2 |
| Sgk2 | Kif13B |
| Ptgds | Sec14L5 |
| Hepacam | Galc |
| Cldn11 | Fyn |
| St18 | Cspg5 |
| Cdo1 | Slc25A47 |
| Ephb1 | Fasn |
| Nfe2L3 | Eml1 |
| Gpr37 | Tmem63A |
| Cnp | Pde8A |
| Myrf | Dpy19L1 |
| Enpp2 | Il33 |
| Arsg | Plekhh1 |
| Tspan15 | Slc25A29 |
| Nnat | Nipal4 |
| Cntf | Vldlr |
| Pllp | Epn2 |
| Mag | Creb5 |
| Aldoc | Anln |
| Bcas1 | Etv5 |
| Fgfr2 | Trf |
| Pnpla2 | Pou3F1 |
| Kif6 | |
| Lhfpl3 | |
| Itgb4 | |
| Mog | |
| Efnb3 | |
| Kcnip3 | |
| Dbndd2 | |
| Tyro3 | |
| Hdac11 | |
| Elovl7 | |
| Matn4 | |
| Ascl1 | |
| Sox8 | |
| Sema4D | |
| Cspg4 | |
| Arrdc2 | |
| Olig1 | |
| Itpr2 | |
| Myo1D | |
| Mcam | |
| Adamts4 | |
| Pex5L | |
| Npc1 | |
| Pdgfra | |
| Slc25A19 | |
| Sgk3 | |
| Gjc2 | |
| Gamt | |
| Lad1 | |
| Ddc | |

### Neuronal

| | | |
|---|---|---|
| Gabbr2 | Nicn1 | Atp6V1H |
| Miat | Cxcr5 | Cxcl1 |
| Cck | Frrs1L | Nrsn2 |
| Scrg1 | Magee1 | Zcchc7 |
| Fmo1 | Atp6V0A1 | Cbarp |
| Cpne6 | Scn8A | Spock3 |
| Elavl3 | Gprasp2 | Cntnap2 |
| Rbfox3 | Safb2 | Arhgdig |
| Ppp1R1B | Disp2 | Zcchc12 |
| Slc11A1 | Rhbdd2 | Tmem130 |
| St8Sia3 | Flywch1 | Slc30A9 |
| Dnm3 | Cnrip1 | Tmem59L |
| Cd59A | Rb1Cc1 | Slc12A5 |
| Pcsk2 | Mecp2 | Vsnl1 |
| Slc1A1 | Wdr60 | Resp18 |
| Npy | Tro | Snhg11 |
| Nwd2 | Myo5A | Snap25 |
| Hap1 | Rnpc3 | Meg3 |
| Chodl | Dipk1b | Pnisr |
| Slc6A17 | Dtd1 | Safb |
| Abat | Cdk5 | Sfswap |
| Dlgap1 | Rere | Rnf220 |
| B3Gat2 | Zmynd11 | Dlg4 |
| Sgip1 | Prph | Ptpn11 |
| Syne1 | Syt5 | Ogfod1 |
| Dner | Tubb3 | Gpatch8 |
| Prune2 | Car10 | Necab3 |
| Cacna2D2 | Icam5 | Zfp445 |
| Dusp26 | Shisa2 | Chd6 |
| Luzp2 | Ndn | Nbea |
| Rasgrf1 | Ptk2B | Fam181B |
| Vgf | Aqp1 | Map7D2 |
| Pcp4 | Pnck | Gnai1 |
| Eno2 | Necab2 | Gria4 |
| Dzank1 | Bex1 | Trip11 |
| Stmn2 | Ccdc28B | Scn1A |
| Pclo | Stx1A | Cspp1 |
| Tceal3 | Pqbp1 | Ppp1R14C |
| Ifit3 | Igf1 | Napg |
| Ddn | Epo | Copg1 |
| Cacnb3 | Ghsr | Arid4A |
| Tmem179 | Chat | Gnl3L |
| Ppp1R9A | Omp | Fbxw7 |
| Rgs17 | Agrp | Pabpn1 |
| Celf6 | Th | Cacna2D1 |
| Tmem191C | Aim2 | Gad1 |
| Rora | Csf3 | Scn2B |
| Eml5 | Avp | Zic1 |
| Cplx1 | Hoxc8 | Gng2 |
| Cacnb4 | Sacs | Kcnma1 |
| Matk | Tac2 | Kcnk1 |
| Hdac11 | Nefh | Rgs7Bp |
| Akap8L | Tmem91 | Aak1 |
| Ncald | Mirg | Tesc |
| Scamp1 | Ube2D2A | Ckmt1 |
| Dgkb | Cacng2 | Slc36A4 |
| Ryr2 | Ctxn2 | Acsl4 |
| Cmip | Tceal6 | Chn1 |
| Zrsr1 | Hpca | Rims3 |
| Pacsin1 | Slc17A6 | Synpr |
| Prox1 | Pnmal1 | Kcnc1 |
| Prmt2 | Nrn1 | Fbln2 |
| Camk2N1 | Tff3 | Pde1A |
| Rin1 | Dync1l1 | Matn2 |
| Gpr162 | Gabrg2 | Calb1 |
| Clstn3 | Tceal5 | Gria3 |
| Kmt2C | Slc32A1 | Gabra1 |
| Bex2 | Nefm | Grm5 |
| Slc17A7 | Nmt1 | Isl1 |
| Cttnbp2 | Rit2 | Nrgn |
| Calb2 | | |

### Progenitor/ Proliferation

| |
|---|
| Fzd9 |
| Lmo3 |
| Wnt5a |
| Bmi1 |
| Slc1a3 |
| Traf4 |
| Efnb1 |
| Serpine2 |
| Zic1 |
| Metrn |
| Lhx2 |
| Hjurp |
| Sox11 |
| Nes |
| Mki67 |
| Kif2c |
| Nptx1 |
| Cenpf |
| Ube2c |
| Top2A |
| Fzd1 |
| Cenpk |
| Cdk1 |
| Lgals1 |
| Ccnd1 |
| Cenpa |
| Igfbp3 |
| Tnc |
| Pou3f1 |
| Ccnd2 |
| Igfbp2 |
| Igfbp4 |

**Supplementary Fig. 2 | Gene panels for Astrocyte lineage, Oligodendrocyte lineage, Neuronal lineage cells and genes associated with progenitor/proliferation state.** Astrocyte gene panel (n=429) was derived from meta-analysis of archival datasets (see Supplementary Fig. 3 and Source data for details). Oligodendrocyte lineage and Neuronal lineage gene panels were derived using PanglaoDB single cell datasets.

**Step 1** **Access astrocyte RNA-Seq Data (n=8 data sets)**

**A** Our work - **Sofroniew Lab (UCLA)** - Healthy mouse Spinal Cord from 73.12 GFAP (**1**) and Aldh1l1-CreERT2 - RiboTag (**2**)

**B** **Khahk Lab (UCLA)** - Healthy mouse Striatum (**3**), Hippocampus (**4**), Cortex (**5**) Aldh1l1-CreERT2 - RiboTag

**C** **Holt Lab (Leuven)** - Healthy mouse Brain - single cell RNA-Seq (**6**)

**D** **Barres Lab (Stanford)** - Healthy mouse Brain - immunopanned (**7**)

**E** **PanglaoDB** gene list for Astrocytes - aggregated database of single cell datsets (**8**)

**Step 2** **Define astrocyte enriched genes (AEG)**

**A** AEG = $Log_2FC$ of RiboTag IP vs flow-through > 2.0 (FDR < 0.01)

**B** AEG = $Log_2FC$ of RiboTag IP vs Input > 2.0 (FDR < 0.01)

**C** AEG = Genes that appear on "common" astrocyte gene list

**D** AEG = $Log_2FC$ > 2.0 for Astrocytes compared with average expression for all other immunopanned cell types (FDR < 0.01)

**E** AEG = all genes on currated Astrocyte gene list

| Data Set | 1 | 2 | 3 | 4 | 5 | 6 | 7 | 8 |
|---|---|---|---|---|---|---|---|---|
| Total number of genes | 1386 | 1456 | 1096 | 873 | 899 | 345 | 944 | 62 |

**Step 3** **Identify genes appearing in at least 5 of 8 lists**

**Total genes in Astrocyte gene list = 429 (Full gene list in source data)**

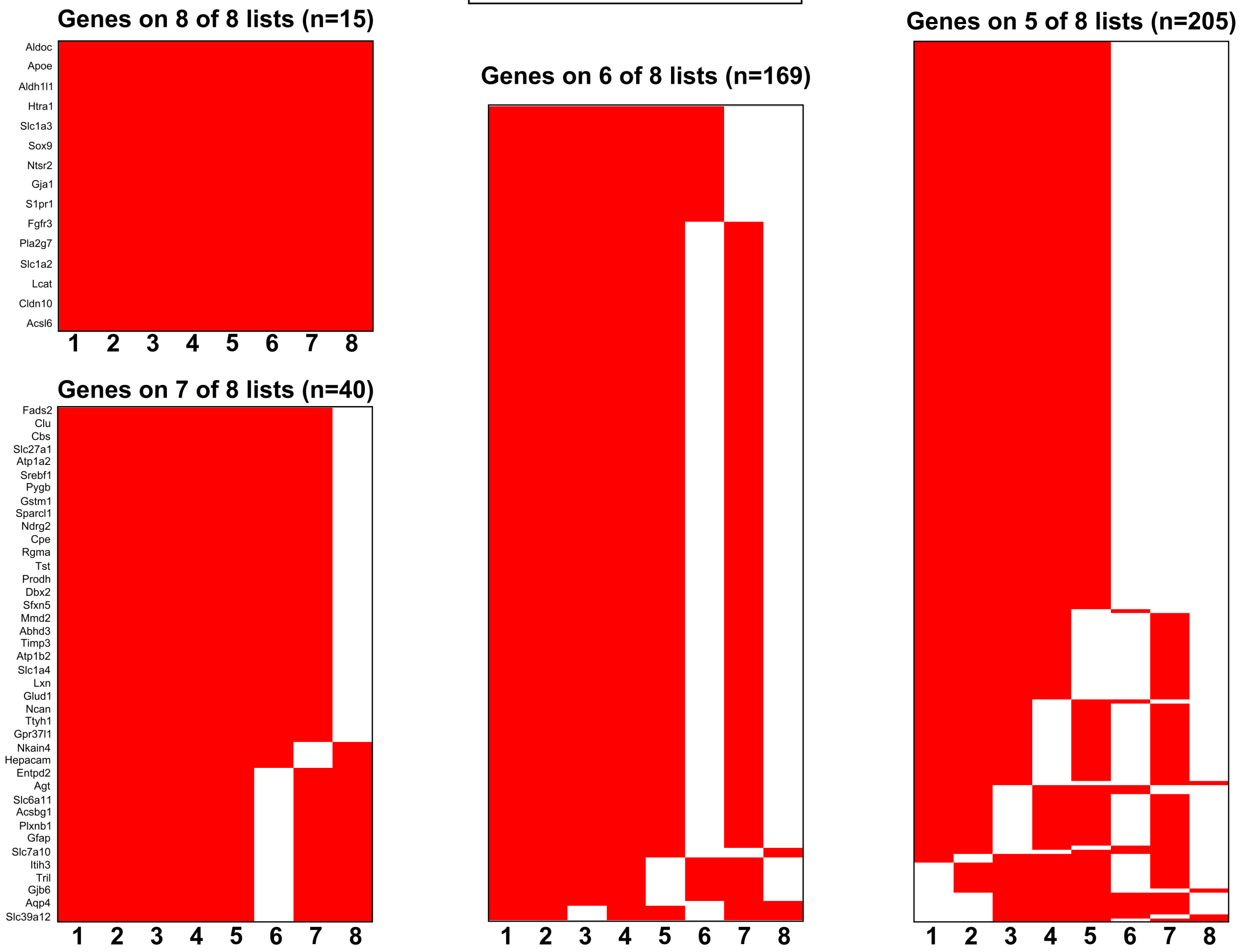


**Supplementary Fig. 3 | Overview of workflow for generating Astrocyte gene panel by meta-analysis of archival datasets.**

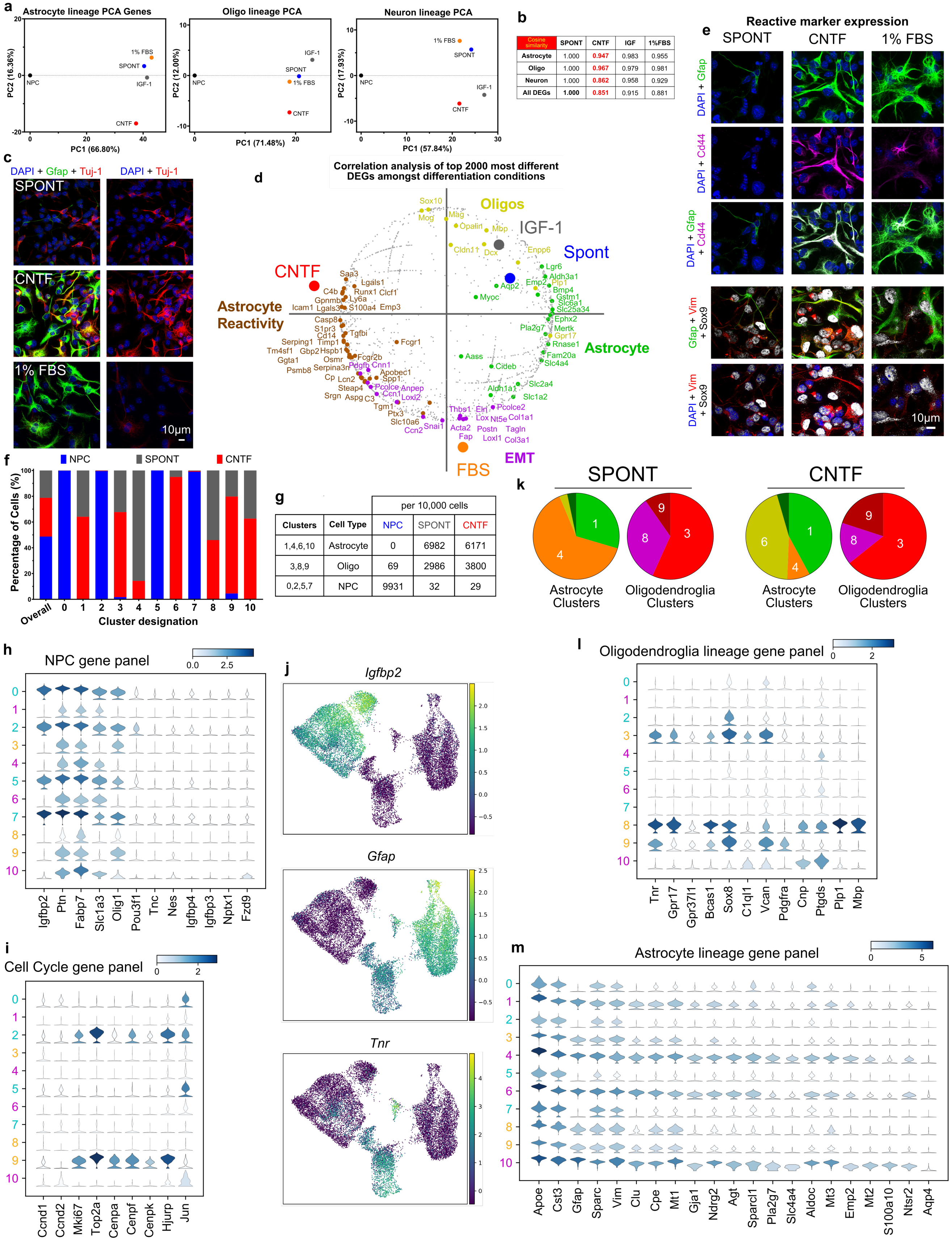


| Cosine similarity | SPONT | CNTF | IGF | 1%FBS |
|---|---|---|---|---|
| Astrocyte | 1.000 | 0.947 | 0.983 | 0.955 |
| Oligo | 1.000 | 0.967 | 0.979 | 0.981 |
| Neuron | 1.000 | 0.862 | 0.958 | 0.929 |
| All DEGs | 1.000 | 0.851 | 0.915 | 0.881 |

| | | per 10,000 cells | | |
|---|---|---|---|---|
| Clusters | Cell Type | NPC | SPONT | CNTF |
| 1,4,6,10 | Astrocyte | 0 | 6982 | 6171 |
| 3,8,9 | Oligo | 69 | 2986 | 3800 |
| 0,2,5,7 | NPC | 9931 | 32 | 29 |

**Supplementary Fig. 4 | NPC differentiate into astrocyte and oligodendroglia lineage cells *in vitro* with phenotype guided by specific molecules. a** PCA for the differentiation conditions using the healthy Astrocyte, Oligodendroglia lineage and Neuron gene panels. **b** Table of cosine similarity comparisons for the differentiation conditions referenced to SPONT differentiation and analyzed using the neural cell type gene panels or all DEGs. CNTF treated samples were least similar to the SPONT differentiation across all analyzes. **c** ICC staining of differentiated cell conditions SPONT, CNTF and 1%FBS for pan neuronal marker Tuj-1 and counter stained with Gfap and DAPI. **d** Correlation analysis of the top 2000 most differential regulated DEGs amongst the 4 differentiation conditions by correlation PCA. CNTF treated samples correlated strongest with astrocyte reactivity genes, FBS treated samples with EMT genes and SPONT and IGF-1 correlated with higher expression of Oligodendroglia lineage and healthy astrocyte lineage genes. **e** ICC staining of differentiated cell conditions SPONT, CNTF and 1%FBS for reactivity markers Cd44 and Vimentin (Vim) and counter stained with Gfap and DAPI. **f** Percentage of cells from NPC, CNTF and SPONT conditions in each of the 11 clusters (0-10) derived from single cell analysis. **g** Table of cell type groupings for the single cell analysis indicating the number of cells per 10,000 for each cell type from each condition. **h** Violin Plots displaying distribution of expression of canonical NPC genes across all cells in the each of cluster designations. Color scale indicates median expression values. **i** Violin Plots displaying distribution of expression of canonical cell cycle or proliferation genes across all cells in the each of cluster designations. Color scale indicates median expression values. **j** Gene expression values across all cells arranged in UMAP space for the genes that most uniquely define NPC (*Igfpb2*), Astrocyte (*Gfap*) and Oligodendroglia (*Tnr*) lineage cells. **k** Pie charts displaying the percentage of each astrocyte and oligodendroglia lineage cluster for the SPONT and CNTF treated samples. While the distribution of oligodendroglia lineage clusters was comparable across the two differentiation conditions, SPONT showed predominant cluster 4 astrocytes whereas CNTF showed predominant cluster 6 astrocytes. **l** Violin Plots displaying distribution of expression of oligodendroglia lineage genes across all cells in the each of cluster designations. Color scale indicates median expression values. **m** Violin Plots displaying distribution of expression of astrocyte lineage genes across all cells in the each of cluster designations. Color scale indicates median expression values.

| Astrocyte Genes | SC Astros | SCI Astros | Spont | CNTF | IGF-1 | 1%FBS | NPC |
|---|---|---|---|---|---|---|---|
| Apoe | 14805.1 | 18634.8 | 9949.9 | 10067.7 | 13946.7 | 12167.7 | 30.7 |
| Mt1 | 7030.8 | 14600.0 | 2645.2 | 2353.3 | 4383.2 | 4926.3 | 48.7 |
| Aldoc | 5902.4 | 2015.2 | 2404.7 | 2141.3 | 2317.1 | 2149.9 | 231.1 |
| Gfap | 5173.0 | 6064.0 | 1935.1 | 5423.3 | 3361.7 | 4564.8 | 3.7 |
| Atp1b2 | 4396.0 | 1999.7 | 732.4 | 973.4 | 635.5 | 1384.2 | 107.5 |
| Ndrg2 | 3894.9 | 1879.8 | 2244.4 | 1456.8 | 2014.3 | 2485.4 | 26.8 |
| Cst3 | 3182.3 | 2648.9 | 3417.4 | 3068.8 | 4205.8 | 4473.9 | 153.5 |
| Clu | 2346.4 | 2862.8 | 726.7 | 1772.0 | 1108.5 | 1187.5 | 8.8 |
| Atp1a2 | 2315.2 | 872.8 | 174.6 | 35.0 | 174.1 | 260.9 | 5.0 |
| Sparc | 1729.4 | 1238.8 | 1088.8 | 1107.8 | 970.7 | 1301.0 | 314.1 |
| S100b | 1544.7 | 1495.3 | 71.7 | 43.5 | 144.1 | 64.6 | 9.0 |
| Agt | 1031.7 | 747.5 | 1140.8 | 587.9 | 1495.3 | 1077.7 | 11.8 |
| S100a1 | 888.4 | 1382.5 | 5.8 | 9.3 | 9.8 | 6.3 | 1.2 |
| Slc1a2 | 851.2 | 271.8 | 14.9 | 5.1 | 9.7 | 33.3 | 3.9 |
| Gja1 | 581.2 | 213.4 | 267.7 | 114.3 | 212.9 | 339.9 | 21.6 |
| Aldh1a1 | 545.8 | 362.9 | 18.0 | 14.6 | 31.8 | 71.7 | 0.7 |
| Aqp4 | 535.1 | 483.0 | 13.0 | 24.5 | 14.1 | 28.6 | 0.3 |
| Pla2g7 | 530.6 | 306.6 | 427.1 | 107.3 | 658.2 | 563.2 | 1.1 |
| Gjb6 | 291.8 | 80.5 | 0.9 | 0.4 | 0.7 | 1.9 | 0.4 |
| Slc7a10 | 247.5 | 45.3 | 0.4 | 0.4 | 0.4 | 0.3 | 0.6 |
| Id3 | 229.3 | 508.4 | 131.1 | 169.1 | 219.7 | 248.8 | 94.0 |
| Slc4a4 | 216.0 | 80.9 | 28.7 | 4.9 | 17.1 | 33.0 | 2.1 |
| Fam107a | 212.7 | 57.6 | 0.2 | 0.1 | 0.2 | 0.7 | 0.2 |
| Mdk | 189.2 | 140.0 | 0.3 | 0.2 | 0.3 | 0.1 | 0.1 |
| Myoc | 180.5 | 39.4 | 0.5 | 0.7 | 2.3 | 1.0 | 0.2 |
| Slc1a3 | 161.2 | 67.3 | 121.7 | 85.8 | 82.6 | 193.1 | 160.8 |
| Etnppl | 65.4 | 23.8 | 0.0 | 0.0 | 0.0 | 0.1 | 0.1 |
| Luzp2 | 59.8 | 24.7 | 0.1 | 0.0 | 0.1 | 0.1 | 0.0 |
| Slc2a4 | 25.8 | 22.1 | 1.2 | 0.5 | 1.7 | 3.6 | 0.2 |

**Supplementary Fig. 5 | Comparison of gene expression levels (average FPKM) for selected astrocyte genes in:** (i) astrocytes from mouse spinal cord (SC) *in vivo*, (ii) astrocytes after SCI *in vivo*, (iii) cells derived from NPC after spontaneous cell autonomous regulated differentiation *in vitro*, or (iv) cells derived from NPC treated with CNTF, FBS, or IGF-1 to promote differentiation *in vitro*.

**a**

### Reactive Astrocyte Gene Panel

#### Acutely Injured astrocyte transcriptomics Data (n=6 data sets)

1 **Sofroniew Lab (UCLA)** - Spinal Cord astrocytes, 5d SCI in 73.12 GFAP-Cre RiboTag, (log2FC>2, FDR<0.01, CPM >10)

2 **Götz lab (Helmholtz)** - Cortical astrocytes, 5d Stab in GFAP-eGFP (Top 50 injury associated genes)

3 **Petzold lab (DZNE)** Brain astrocytes, 3d MCAO in Cx43-CreERT-RiboTag (log2FC>2, p<0.05)

4 **Barres Lab (Stanford)** - mouse Brain, MCAO 24 hrs - immunopanned for astrocytes (Top 50 injury associated genes)

5 **Wu Lab (Houston)** - Spinal Cord astrocytes, 7d SCI in 73.12 GFAP-Cre RiboTag, (log2FC>2)

6 **Voskuhl Lab (UCLA)** - Spinal Cord astrocytes, EAE in 73.12 GFAP-Cre RiboTag, (log2FC>2, FDR <0.1)

**Identify genes appearing in at least 3 of 6 lists**

**Total genes in Reactive gene list = 170 (Full gene list in source data)**

**b**

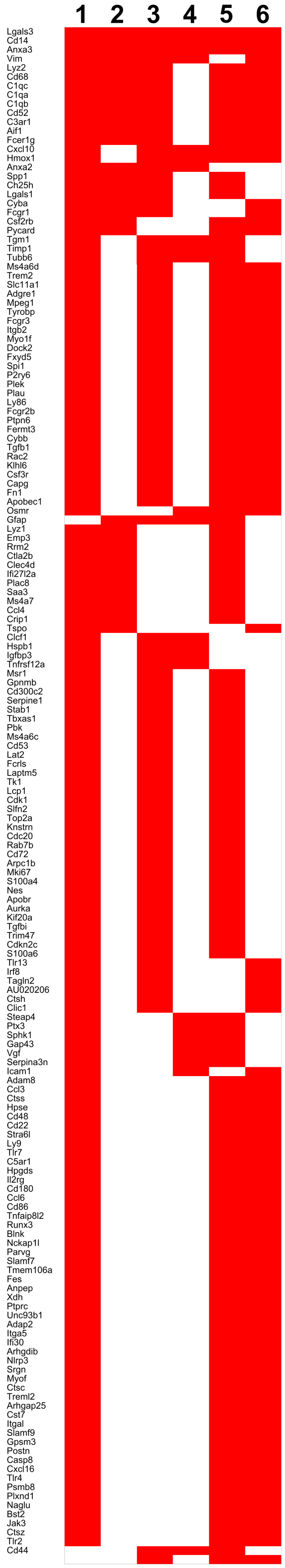


**c**

### EMT Gene Panel

Derived from EMT Gene Set Enrichment Analysis (GSEA) dataset from Molecular Signatures Database (Msigdb) with human specific genes removed

| **EMT gene panel** | | | |
|---|---|---|---|
| Abi3Bp | Dst | Lama1 | Ptx3 |
| Acta2 | Ecm1 | Lama2 | Pvr |
| Adam12 | Ecm2 | Lama3 | Qsox1 |
| Anpep | Edil3 | Lamc1 | Rgs4 |
| Aplp1 | Efemp2 | Lamc2 | Rhob |
| Areg | Eln | Lgals1 | Sat1 |
| Basp1 | Emp3 | Lox | Scg2 |
| Bdnf | Eno2 | Loxl1 | Sdc1 |
| Bgn | Fap | Loxl2 | Sdc4 |
| Bmp1 | Fas | Lrp1 | Serpine1 |
| Cadm1 | Fbln1 | Lrrc15 | Serpine2 |
| Cald1 | Fbln2 | Lum | Serpinh1 |
| Calu | Fbln5 | Magee1 | Sfrp1 |
| Cap2 | Fbn1 | Matn2 | Sfrp4 |
| Capg | Fbn2 | Matn3 | Sgcb |
| Ccn1 | Fermt2 | Mcm7 | Sgcd |
| Ccn2 | Fgf2 | Mest | Sgcg |
| Cd44 | Flna | Mfap5 | Slc6A8 |
| Cd59A | Fmod | Mgp | Slit2 |
| Cdh11 | Fn1 | Mmp13 | Slit3 |
| Cdh2 | Foxc2 | Mmp14 | Snai2 |
| Cdh6 | Fstl1 | Mmp2 | Sntb1 |
| Col11A1 | Fstl3 | Mmp3 | Sparc |
| Col12A1 | Fuca1 | Msx1 | Spock1 |
| Col16A1 | Fzd8 | Myl9 | Spp1 |
| Col1A1 | Gadd45A | Mylk | Tagln |
| Col1A2 | Gadd45B | Nid2 | Tfpi2 |
| Col3A1 | Gas1 | Nnmt | Tgfb1 |
| Col4A1 | Gem | Notch2 | Tgfbi |
| Col4A2 | Gja1 | Nt5E | Tgfbr3 |
| Col5A1 | Glipr1 | Ntm | Tgm2 |
| Col5A2 | Gpc1 | Oxtr | Thbs1 |
| Col5A3 | Gpx7 | P3H1 | Thbs2 |
| Col6A2 | Grem1 | Pcolce | Thy1 |
| Col6A3 | Htra1 | Pcolce2 | Timp1 |
| Col7A1 | Id2 | Pdgfrb | Timp3 |
| Col8A2 | Igfbp2 | Pdlim4 | Tnc |
| Colgalt1 | Igfbp3 | Pfn2 | Tnfaip3 |
| Comp | Igfbp4 | Plaur | Tnfrsf11B |
| Copa | Il15 | Plod1 | Tnfrsf12A |
| Crlf1 | Il6 | Plod2 | Tpm1 |
| Cthrc1 | Inhba | Plod3 | Tpm2 |
| Cxcl1 | Itga2 | Pmepa1 | Tpm4 |
| Cxcl12 | Itga5 | Pmp22 | Vcam1 |
| Cxcl5 | Itgav | Postn | Vcan |
| Dab2 | Itgb1 | Ppib | Vegfa |
| Dcn | Itgb3 | Prrx1 | Vegfc |
| Dkk1 | Itgb5 | Prss2 | Vim |
| Dpysl3 | Jun | Pthlh | Wipf1 |
| | | | Wnt5A |

**d**

### Overlap of Astrocyte State Gene Panels

| Astrocytes & Reactivity | Reactivity & EMT | Astrocytes & EMT | In all 3 lists |
|---|---|---|---|
| C4b | Anpep | Ccn1 | Vim |
| Gfap | Capg | Gja1 | |
| | Cd44 | Htra1 | |
| | Emp3 | Mmp14 | |
| | Fn1 | Notch2 | |
| | Igfbp3 | Pdlim4 | |
| | Itga5 | Sdc4 | |
| | Lgals1 | Serpine2 | |
| | Postn | Sparc | |
| | Ptx3 | Timp3 | |
| | Serpine1 | Tnc | |
| | Spp1 | Vcam1 | |
| | Tgfb1 | | |
| | Tgfbi | | |
| | Timp1 | | |
| | Tnfrsf12a | | |

**Supplementary Fig. 6 | Reactivity and EMT gene panels. a** Workflow for generating reactivity gene panel in astrocytes. **b** List of reactivity gene panel (also available in source data). **c.** EMT gene panel. **d.** Overlap of the astrocyte state gene panels.

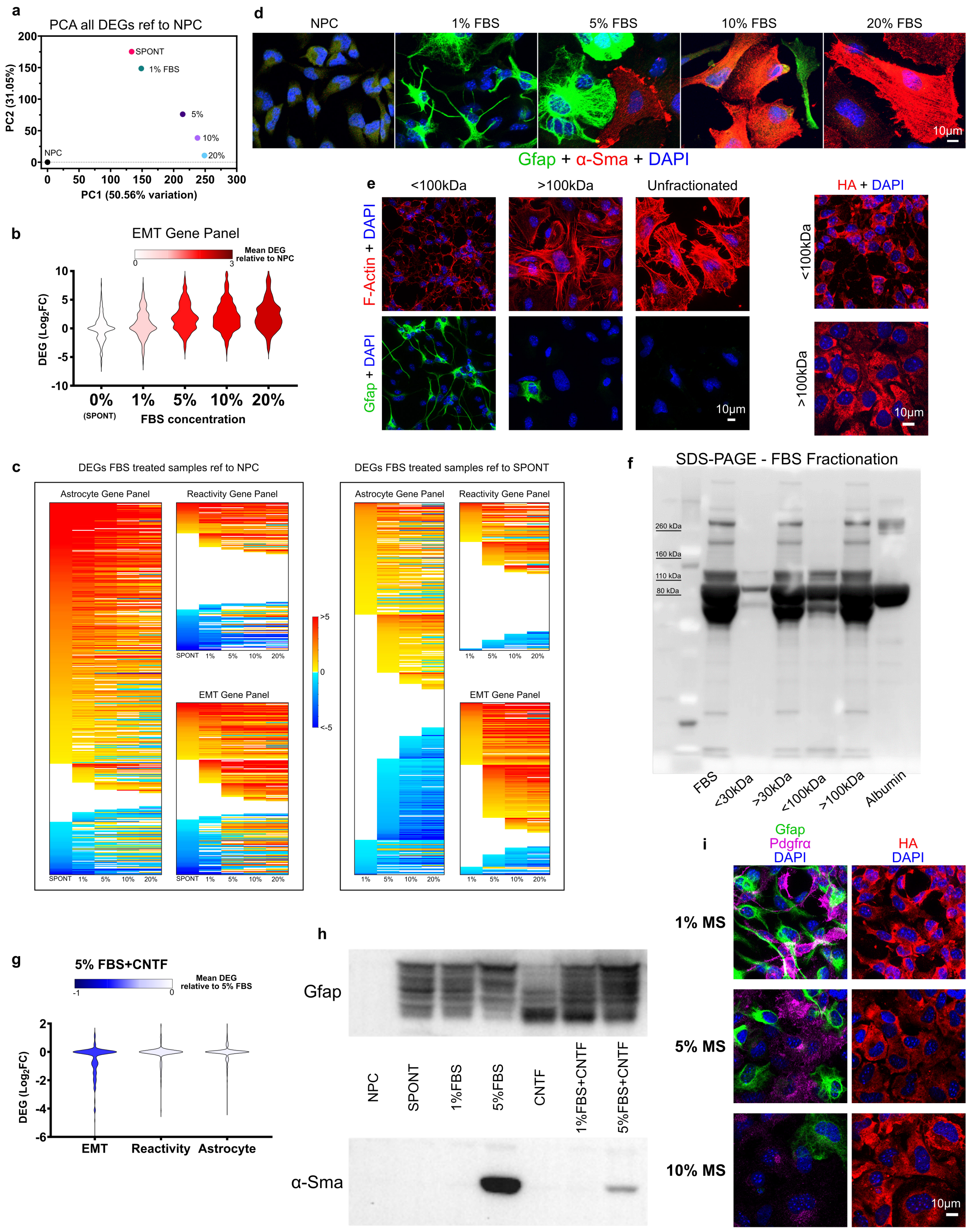


**Supplementary Fig. 7 | Serum exposure evokes myofibroblast-like phenotypes in NPC *in vitro*.** **a** PCA of all DEGs for SPONT and increasing concentrations of FBS referenced to NPC showing increased divergence from NPC with increased FBS concentration. **b** Violin Plot of DEGs for SPONT and increasing concentrations of FBS referenced to NPC for EMT gene panel genes showing a concentration dependent increase in expression of EMT gene panel genes. **c** Heatmaps displaying DEGs for increasing concentrations of FBS referenced to either NPC or SPONT separated into astrocyte, reactivity and EMT gene panels. **d** ICC staining of NPC exposed to increasing concentrations of FBS showing changes in Gfap and α-Sma expression as a function of FBS concentration. Cells counterstained with DAPI. **e** ICC staining of NPC exposed to FBS fractionated by molecular size cut-off centrifugal filters. NPC treated with the <100 kDa fraction, >100kDa fraction and unfractionated FBRS and stained with phalloidin to detect F-Actin expression and counter-stained with Gfap and DAPI. NPC treated with fractionated FBS samples also stained with HA and DAPI. **f** SDS-PAGE of FBS and fractionated samples stained with SYPRO Ruby Protein Gel Stain showing the effectiveness of molecular size cut-off centrifugal filters (30kDa and 100kDa) to separate out higher molecular weight constituents. Unfractionated FBS and albumin used as controls. **g** Violin Plot of DEGs for NPC treated with 5%FBS and CNTF referenced to 5%FBS alone showing the effect of CNTF on attenuating expression of EMT genes. **h** Western blot of FBS and CNTF treated samples stained for Gfap and α-Sma showing CNTF and FBS dependent effects on expression of Gfap Isoforms and α-Sma. **i** ICC staining of NPC exposed to increasing concentration of mouse serum (MS) showing decreased expression of Gfap and Pdgfr-α and altered cell size and morphology indicated by HA staining. Note that the heightened myofibroblast phenotypes caused by increasing concentrations of FBS involved elevated expression of canonical EMT transcription factors (*Snai1*, *Snai2, Twist1*), myofibroblast cytoskeletal proteins (*Acta2, Tagln*), profibrotic growth factors (*Ccn2* (Ctgf), *Tgfbi* (Tgf-β), *Il6*) and fibrotic extracellular matrix proteins (*Fap, Col1a1, Fn1, Thbs1*). The FBS concentration dependent downregulation in astrocyte genes was most apparent for genes associated with healthy astrocyte functions such as molecular transport and included *Slc6a11, Atp1a2, Rlbp1, Slc6a1, Kcnj1, Agt* and *Pla2g7*. Canonical astrocyte gene *Gfap* also showed a strong serum concentration dependent decrease in expression that directly opposed the trend in expression for canonical EMT associated gene *Acta2* (smooth muscle actin (α-Sma)) which increased exponentially.

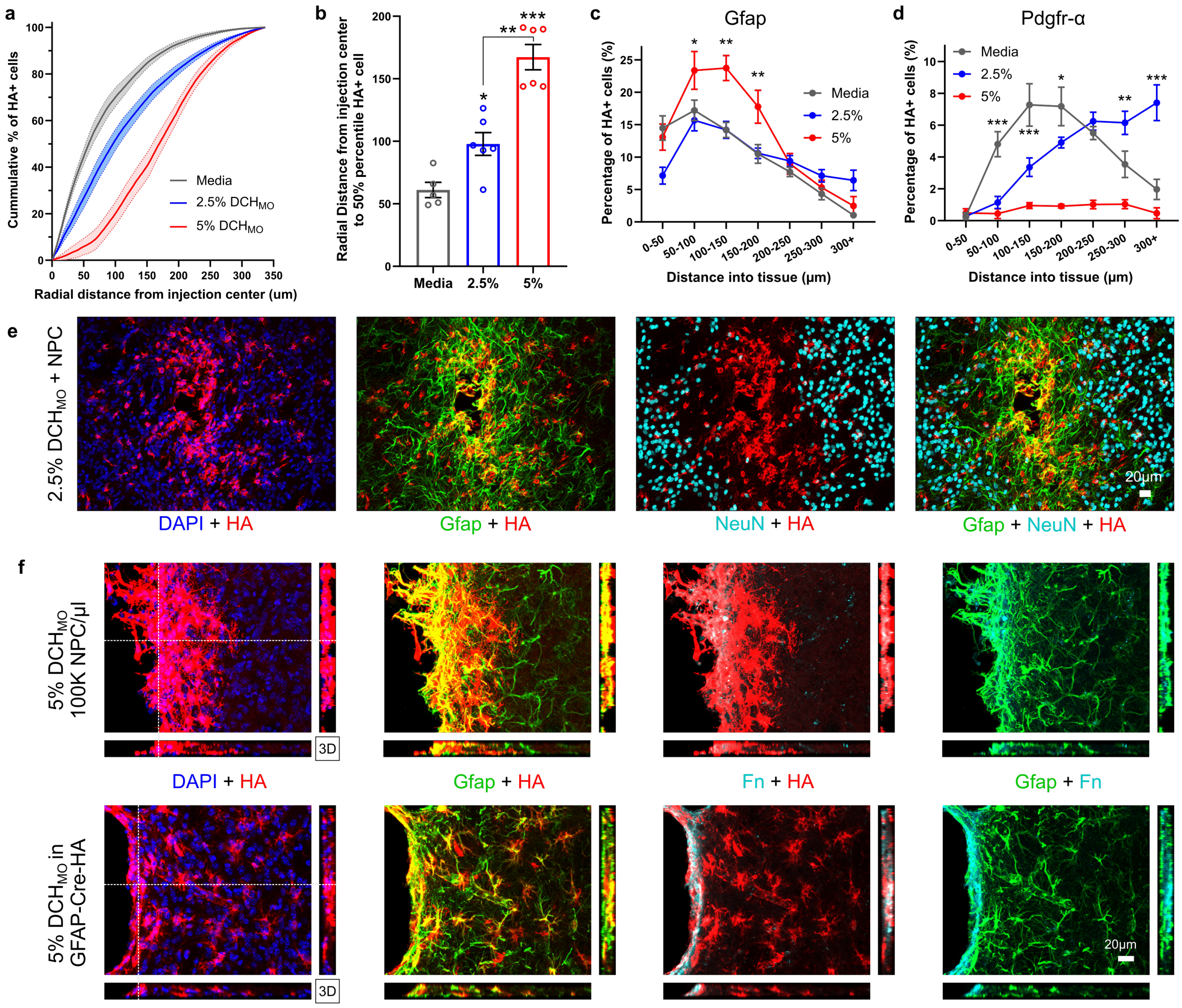


**Supplementary Fig. 8 | NPC grafted into healthy mouse striatum differentiate into astrocyte and oligodendroglia lineage cells in a spatial and carrier dependent manner. a** Quantification of the cumulative spatial distribution of HA+ cells, referenced from the center of injection, for media and $DCH_{MO}$ hydrogel carrier samples. For each sample the darkened line represents the mean and shaded area indicates standard error of mean (SEM) (N=6 for both hydrogels, N=5 for Media). **b** Quantification of the Radial distance to the $50^{th}$ percentile HA+ cell, referenced from the center of injection, for media and $DCH_{MO}$ hydrogel carrier samples. (Mean + SEM) (N=6 for both hydrogels, N=5 for Media). *** p-value < 0.0001, ** p-value < 0.0002, * p-value < 0.032, One way ANOVA with Tukey multiple comparison test. **c** Percentage distribution of HA-positive and Gfap-positive cells within define spatial zones, referenced from the center of injection, for media and $DCH_{MO}$ hydrogel carrier samples. (Mean + SEM) (N=6 for both hydrogels, N=5 for Media). *** p-value < 0.0001, ** p-value < 0.005, * p-value < 0.03, One way ANOVA with Tukey multiple comparison test. **d** Percentage distribution of HA-positive and Pdgfrα-postive cells within define spatial zones, referenced from the center of injection, for media and $DCH_{MO}$ hydrogel carrier samples. (Mean + SEM) (N=6 for both hydrogels, N=5 for Media). *** p-value < 0.0001, ** p-value < 0.005, * p-value < 0.03, One way ANOVA with Tukey multiple comparison test. **e** Survey Immunohistochemistry (IHC) images of grafted HA-positive NPC in healthy striatum stained for astrocytes (Gfap) and neurons (NeuN) at 2 weeks after NPC transplantation. HA-positive grafted NPC do not express NeuN indicating they do not differentiate into Neurons in healthy striatum at 2 weeks. **f** High magnification 3D view showing HA-positive grafted NPC in 5% $DCH_{MO}$ hydrogel carrier showing formation of Gfap-positive borders by grafted NPC that express Fibronectin (FN). Host astrocyte borders are formed around 5% $DCH_{MO}$ hydrogel carrier that are morphologically similar to the borders formed by grafted NPC and also expressed Fibronectin at the border interface.

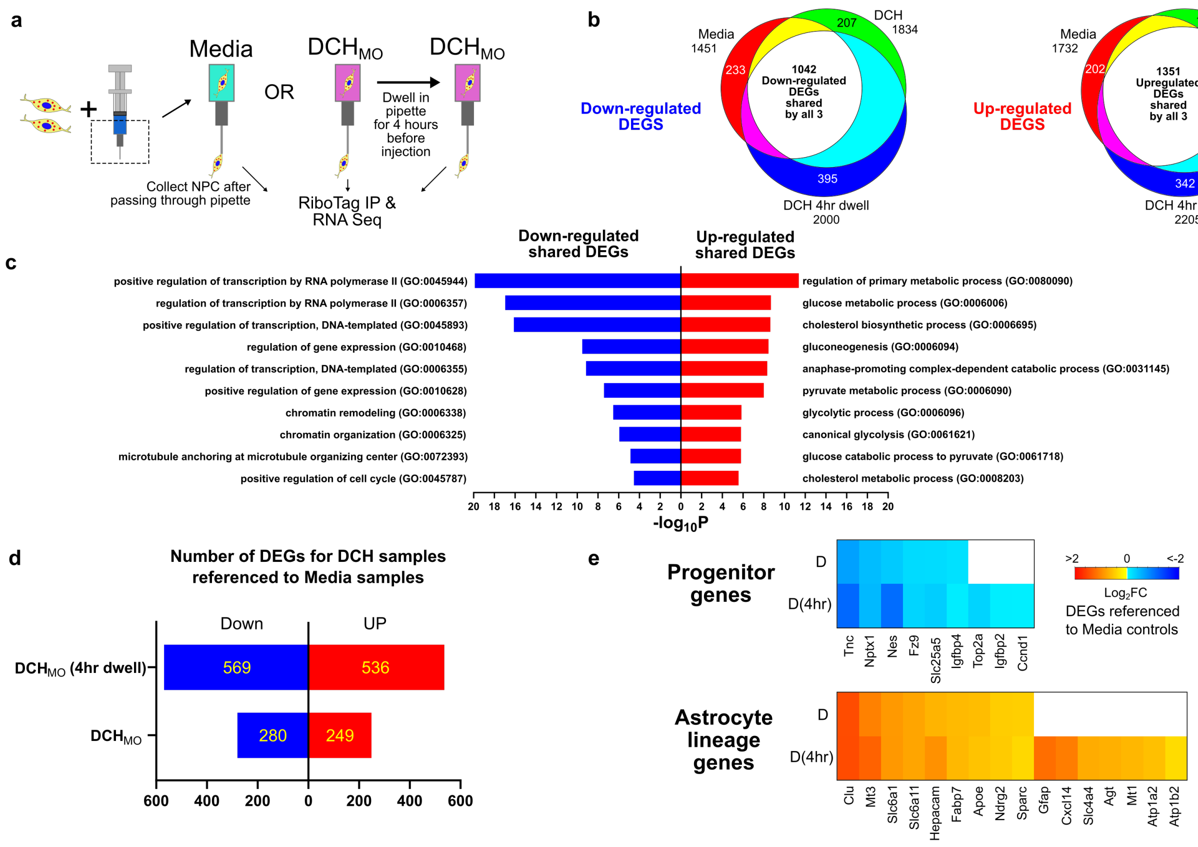


**Supplementary Fig. 9 | NPC injected through glass pipettes are minimally altered by the carrier vehicle used. a** Schematic detailing the experimental paradigm to test the effect of injection through pulled glass pipettes in media or hydrogel carrier on NPC phenotype. In one group, NPC loaded into hydrogel carriers were allowed to dwell in the pipette for 4 hours before being passed through the pipette to evaluate changes to NPC phenotype taking place over a typical transplantation session. **b** Ven diagram of Upregulated and Downregulated DEGs for NPC passed through pipettes in media or hydrogel or allowed to dwell in hydrogel for 4 hours showing high overlap of shared DEGs across the three conditions. **c** Gene Ontology (GO) term analysis of the shared Up-regulate and Down-regulated DEGs. Shared down-regulated DEGs indicate an overall decrease in transcription and cell cycle processes as a result of injection through pipettes. Shared Up-regulated DEGs suggests increased metabolic processes in NPC upon injection through pulled glass pipettes. **d** Total number of hydrogel specific upregulated and downregulated DEGs when referenced to NPC in media samples (FDR <0.01). The 4-hour dwell in the pipette effectively doubles the number of DEGs. **e** Heat map of Astrocyte and Progenitor genes altered in hydrogel samples referenced to NPC in media samples. NPC in hydrogel carriers show an increase in astrocyte gene expression and a decrease in progenitor gene expression with the 4-hour dwell enhancing both effects. Note that there were 1351 shared upregulated genes that were consistent with increased metabolic processes as assessed by Gene Ontology term enrichment, whereas 1042 shared downregulated genes suggested a reduction in cell cycle and transcription processes. Comparisons of hydrogel and media carrier samples identified a modest increase in astrocyte differentiation signature in the hydrogel group with upregulation of astrocyte lineage genes *Clu, Mt3, Slc6a1, Hepacam* among others and a concurrent downregulation of some progenitor genes such as *Igfbp4, Tnc, Nes* and *Slc25a5*. Persistent retention of cells in the pipette for 4 hours further increased the expression of these astrocyte genes, increased expression of other astrocyte genes (including Gfap), and caused further downregulation of progenitor genes which now included *Top2a, Igfbp2* and *Ccnd1*, suggesting that the cells may undergo some differentiation over the course of a standard injection session and that astrocyte phenotypes emerge more quickly than other definable phenotypes. However, despite these apparent DEGs caused by dwell time differences in the pipette, we observed no obvious trend among animals that received transplants first or last in an injection session suggesting these small variations in DEGs may not significantly impact overall graft outcomes *in vivo*.

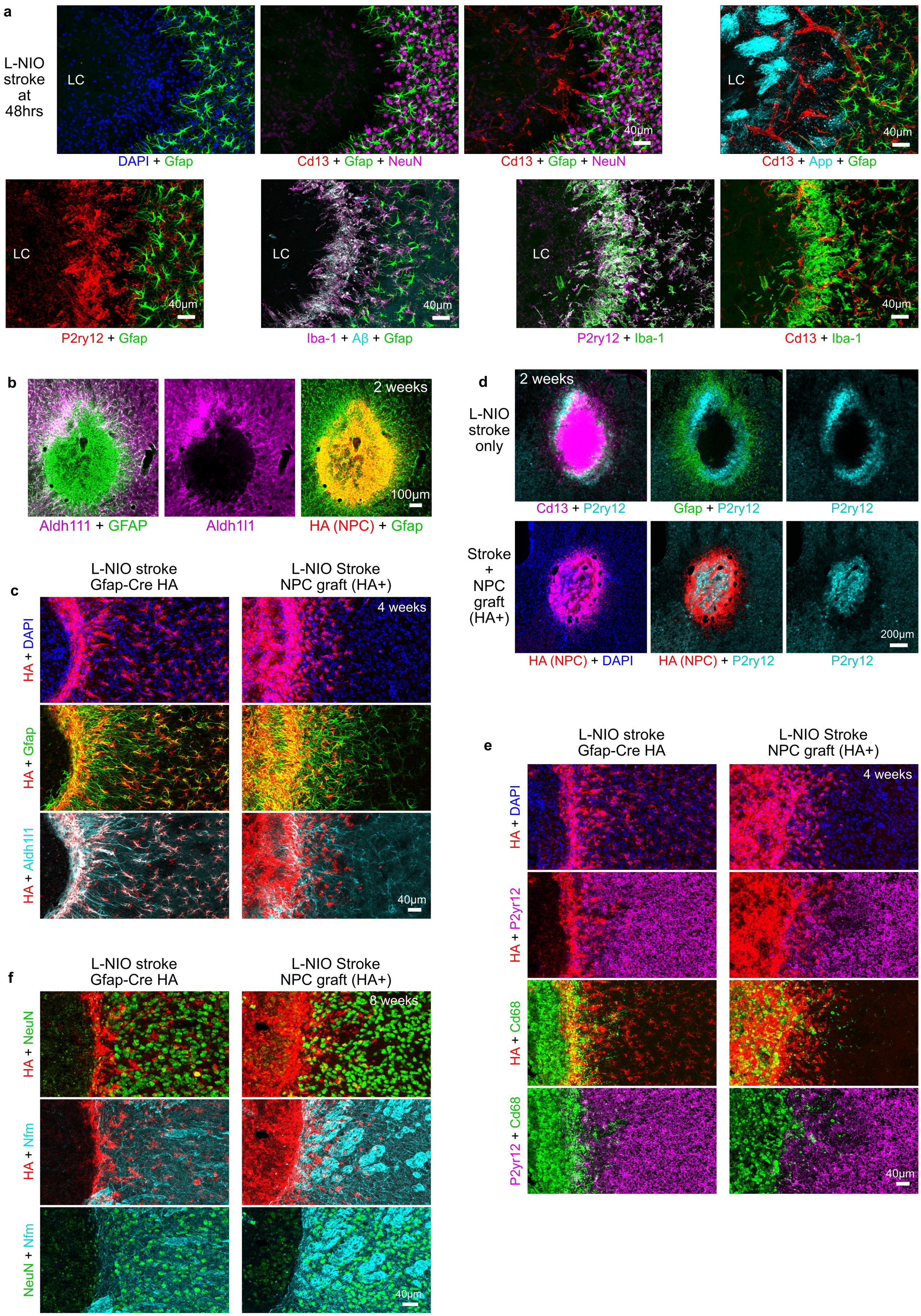
a
L-NIO stroke at 48hrs
LC
DAPI + Gfap
Cd13 + Gfap + NeuN
Cd13 + Gfap + NeuN
40µm
LC
40µm
Cd13 + App + Gfap
LC
40µm
P2ry12 + Gfap
LC
40µm
Iba-1 + Aβ + Gfap
LC
P2ry12 + Iba-1
40µm
Cd13 + Iba-1
b
Aldh111 + GFAP
Aldh1l1
2 weeks
100µm
HA (NPC) + Gfap
d
L-NIO stroke only
2 weeks
Cd13 + P2ry12
Gfap + P2ry12
P2ry12
Stroke + NPC graft (HA+)
200µm
HA (NPC) + DAPI
HA (NPC) + P2ry12
P2ry12
c
L-NIO stroke Gfap-Cre HA
L-NIO Stroke NPC graft (HA+)
4 weeks
HA + DAPI
HA + Gfap
HA + Aldh1l1
40µm
e
L-NIO stroke Gfap-Cre HA
L-NIO Stroke NPC graft (HA+)
4 weeks
HA + DAPI
HA + P2yr12
HA + Cd68
P2yr12 + Cd68
40µm
f
L-NIO stroke Gfap-Cre HA
L-NIO Stroke NPC graft (HA+)
8 weeks
HA + NeuN
HA + Nfm
NeuN + Nfm
40µm

**Supplementary Fig. 10 | NPC grafted into striatal stroke generate astroglia that reduce fibrotic scar and bridge lesions. a** Survey IHC images showing stroke lesion core (LC) at 2 days after stroke induction by L-NIO injection. Stroke lesions are defined by infarcted tissue that shows complete loss of neurons (NeuN) and glia (Gfap, P2ry12), accumulation of axon damage (App), onset of acute inflammation and Aβ demarcated at the border of the forming lesion. **b** Survey IHC images showing that NPC differentiate into Gfap-positive astroglia upon grafting into stroke lesions but do not express mature astrocyte markers such as Aldh1l1. **c** Detailed lesion interface IHC images of L-NIO stroke lesions at 4 weeks after stroke in 73.12 Gfap-Cre HA mice. HA-positive host astrocytes that express Gfap and Aldh1l1 form discrete borders with non-neural lesion cores. NPC grafting into L-NIO strokes comprehensively fill the lesion with Gfap-positive cells, but only grafted cells located at the host neural tissue interface begin to express Aldh1l1 at 4 weeks. **d** Survey IHC image showing NPC treated stroke lesions transiently containing P2yr12-positive CNS derived microglia at 2 weeks whereas untreated stroke lesions do not contain P2yr12-positive microglia in the lesion core with Cd13-positive macrophages and fibroblasts instead predominating in this lesion compartment. **e** Detailed IHC images showing an NPC graft induced change in Cd68-positive immune cells within stroke lesion cores at 4 weeks compared with untreated strokes made in Gfap-Cre HA mice that have HA-positive host astrocytes. P2yr12-positive microglia are no longer detected within the graft. **f** Detailed IHC images showing that NPC grafting into stroke lesions does not provoke neuronal repopulation or striatal axon regrowth into lesion cores by 8 weeks.

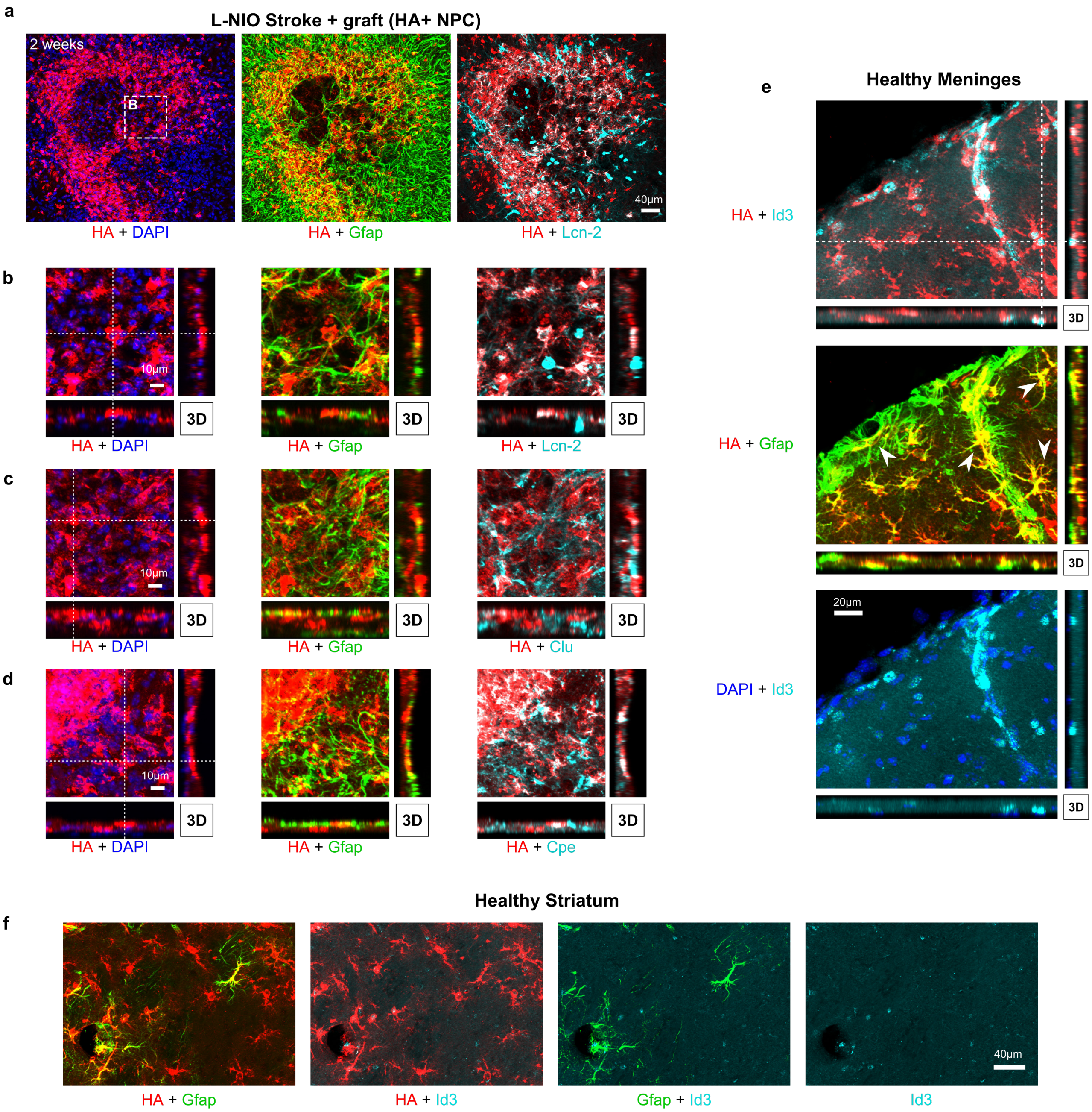


**Supplementary Fig. 11 | Astroglia derived from NPC grafted into CNS lesions become border-forming astroglia. a** Survey IHC imaging of NPC grafted into L-NIO stroke lesions showing Gfap-positive, HA-positive grafted cells expressing the astrocyte reactivity marker Lipocalin-2 (Lcn-2) in cells forming borders around lesion cores. **b** Detailed, higher magnification 3D imaging showing co-localization of Lcn-2 in Gfap-positive, HA-positive grafted cells in stroke. **c** Detailed, higher magnification 3D imaging showing co-localization of astrocyte reactivity marker Clusterin (apolipoprotein J) (Clu) in Gfap-positive, HA-positive grafted cells in stroke. **d** Detailed, higher magnification 3D imaging showing co-localization of astrocyte marker Carboxypeptidase E (Cpe) in Gfap-positive, HA-positive grafted cells in stroke. **e** Detailed, higher magnification 3D imaging showing co-localization of Id3 in HA-positive host astrocytes. Several Id-3, HA and Gfap colocalized host astrocytes are indicated by overlayed arrows. **f** Survey IHC imaging of healthy mouse striatum showing relatively low expression of Id3 in protoplasmic astrocytes in the striatum. Astrocytes that form the glia limitans around blood vessels in the striatum show higher Id3 expression.

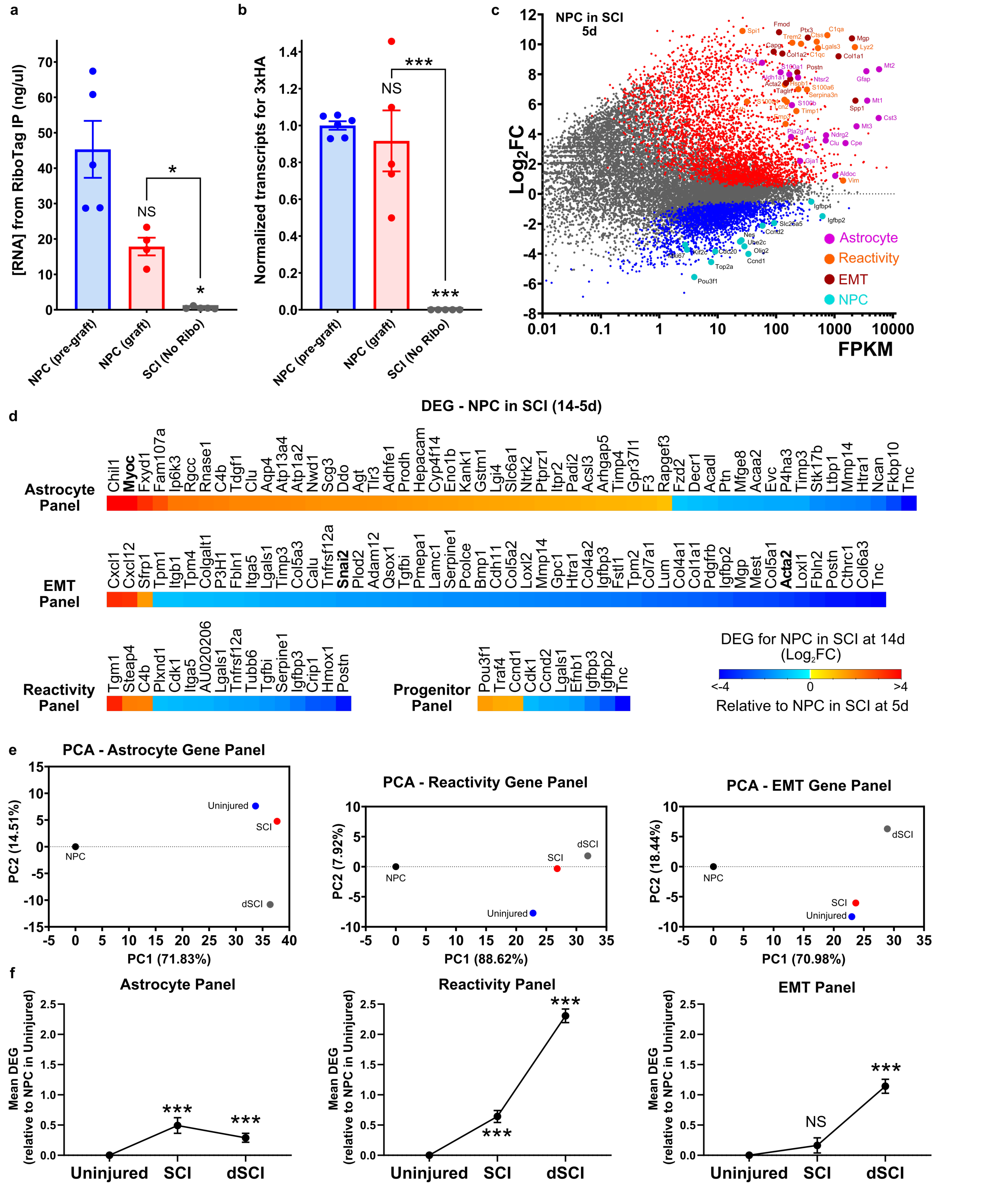


**Supplementary Fig. 12 | RiboTag transcriptomics on NPC transplanted into subacute crush spinal cord injury reveals an astroglial border phenotype that evolves over time and is altered by lesion severity. a** Concentration of RNA recovered following RiboTag Immunoprecipitation and mRNA purification from NPC before grafting and 5 days after grafting into a 2-day crush SCI lesion. RNA recovered from NPC after grafting is not significantly reduced compared to pre-graft levels. There is minimal recovery of RNA from SCI lesions that did not receive a graft of HA-positive cells (SCI (no RiboTag)) suggesting high specificity towards recovery of grafted cell RNA using the RiboTag methods. **b** Normalized counts of 3xHA transcripts from RNA-Seq data sets for pre-graft NPC and NPC recovered at 5 days after grafting into a 2-day crush SCI lesion showing no statistical differences in the frequency of detection of 3xHA transcripts in samples recovered from the graft compared to the pre-graft sample that contains 100% HA-positive cells. No 3xHA transcripts were detected in samples recovered from SCI tissues that did not receive a graft of HA-positive cells. **c** MA plot of FPKM levels versus DEGs for NPC grafted into 2-day crush SCI lesions (N=5 mice). Significant Up (red) and Down (Blue) DEGs have FDR

<0.01. Select DEGs for astrocytes, reactivity, EMT and progenitor genes are indicated. **d** Heatmaps of all astrocyte, reactivity, EMT and NPC/progenitor gene panel DEGs detected in NPC grafts recovered at 14 days after grafting when compared with NPC grafts recovered at 5 days after grafting into 2-day SCI lesions (N=5 for both time points). DEGs have FDR <0.01. NPC recovered at 14 days show overall net increase in expression of astrocyte genes and a decrease in expression of EMT genes compared with the grafts recovered at 5 days. (N=5 mice per group). **e** PCA of astrocyte, reactivity and EMT gene panels for NPC grafted into different tissue environments (uninjured, SCI and double crush SCI (dSCI)). (N=5 mice per group). **f** Plots of Mean DEGs for astrocyte, reactivity and EMT gene panels for NPC grafts recovered from SCI and dSCI referenced to NPC grafts recovered from uninjured spinal cord tissue. (N=5 mice per group) *** p-value < 0.0001, NS = not significant, One way ANOVA with Tukey multiple comparison test.
